# Astrocyte dysfunction distinguishes monozygotic twin *C9orf72* expansion carriers discordant for amyotrophic lateral sclerosis

**DOI:** 10.64898/2026.07.15.738192

**Authors:** Allan C. Shaw, Iris S. Pasniceanu, Rebecca Stevenson, Chloé Moutin, Gergo Erdi-Krausz, Matthew Parker, Matthew Wyles, Cleide Dos Santos Souza, Adrian Higginbottom, Lydia Castelli, Janine Kirby, Guillaume M. Hautbergue, Laura Ferraiuolo, Johnathan Cooper-Knock, Matthew R. Livesey, Pamela J. Shaw

## Abstract

Hexanucleotide repeat expansions in *C9orf72* are the most common genetic cause of amyotrophic lateral sclerosis, yet many carriers remain asymptomatic for decades or never develop disease. This incomplete penetrance suggests that phenoconversion from genetic susceptibility to symptomatic disease onset is governed by epigenetic, environmental and cell-intrinsic modifiers. Astrocytes are key mediators of non-cell-autonomous neurodegeneration in ALS, but whether they undergo disease-associated phenoconversion in C9orf72 expansion carriers remains unclear.

We investigated astrocyte-associated mechanisms of phenoconversion using a unique *C9orf72* pedigree comprising monozygotic twins discordant for ALS and their asymptomatic father. This enabled analysis across a continuum from non-penetrance to late-stage neurodegeneration while controlling for inherited genetic background. Notably, the unaffected twin exhibited hypermethylation of the expanded *C9orf72* allele, identifying an epigenetic correlate of non-penetrance.

To define symptomatic and asymptomatic-associated astrocyte states, fibroblasts from this pedigree were directly reprogrammed into astrocytes and subjected to molecular, functional, electrophysiological and translatome analyses. Astrocytes derived from symptomatic ALS individuals were found to exhibit astrocyte-mediated toxicity towards motor neurons, canonical C9orf72 molecular pathology and connexin-related membrane dysfunction. Specifically, RNA foci and dipeptide repeat protein burden peaked in astrocytes obtained early in disease in the symptomatic twin and declined in astrocytes derived from samples obtained at advanced disease stages, whereas motor neuron toxicity increased progressively, demonstrating a dissociation between aggregate burden and functional neurotoxicity. In contrast, connexin-mediated electrophysiological dysfunction emerged with symptomatic disease and closely tracked with maximal toxicity. Translatome profiling revealed early global translational repression and impaired proteostasis in symptomatic astrocytes, whereas unaffected and non-penetrant C9-carriers retained enrichment of protein homeostasis pathways. These findings define distinct astrocyte states associated with asymptomatic, early symptomatic and late-stage disease.

Notably, the affected twin reported substantially higher lifetime strenuous physical activity compared to the unaffected twin, consistent with a potential role for sustained exercise-related stress in accelerating disease onset in genetically susceptible individuals. Experimentally modelling increased stress induced aberrant upregulation of connexin-mediated currents in *C9orf72* astrocytes, including in asymptomatic carriers, indicating that physiological stress can unmask latent astrocyte-intrinsic vulnerability and precipitate dysfunction in cellular homeostasis.

Together, these findings redefine phenoconversion in *C9orf72*-associated amyotrophic lateral sclerosis to be associated with a significant upregulation of astrocyte toxicity, failure of resilience and demonstrate that the astrocyte disease state can be potentially induced by external stressors.

## Introduction

Amyotrophic lateral sclerosis (ALS) is a rapidly progressive neurodegenerative disorder characterised by degeneration of upper and lower motor neurons, leading to spreading paralysis and death, typically within 2–3 years of symptom onset.^1^ Although multiple converging mechanisms of motor neuron injury have been implicated including: proteostatic failure, mitochondrial dysfunction, oxidative stress, neuroinflammation, excitotoxicity and non–cell-autonomous glial toxicity – the critical biological transition from decades-long presymptomatic vulnerability to clinical disease-onset (phenoconversion) remains poorly understood.^2^

Hexanucleotide repeat expansions in *C9orf72* (C9-HRE) are the most common genetic risk factor of ALS and frontotemporal dementia, accounting for ∼40% of familial ALS and up to 10% of sporadic cases.^3–5^ Despite a shared mutation, clinical penetrance is incomplete and age-of-onset varies widely, even within families.^6–10^ This indicates that genotype alone is insufficient to account for disease emergence, and that epigenetic state, environmental exposures, and cell-intrinsic resilience mechanisms likely govern phenoconversion. Mechanistically, C9-HRE pathology is linked to toxic RNA foci, repeat-associated non-AUG (RAN) translation producing dipeptide repeat proteins (DPRs), C9orf72 haploinsufficiency, and downstream TDP-43 proteinopathy. However, how and when these processes become functionally operational in human brain cells remains unresolved.^11–14^

Astrocytes are emerging as central regulators of motor neuron (MN) viability through metabolic support, neurotransmitter and ion homeostasis, and modulation of neuroinflammation, but also as drivers of non–cell-autonomous neurotoxicity in ALS.^15–28^ Yet, a key unresolved question is whether astrocytes acquire disease-associated toxic states prior to symptom onset, and what triggers this transition. Given that ALS typically manifests in mid-to-late adulthood, astrocytic dysfunction is unlikely to be constitutive, implying dynamic, disease stage-dependent reprogramming of glial states during disease evolution.

Physical exercise and neuronal activity impose metabolic, oxidative, and ionic stress on astrocytes and have been epidemiologically associated with increased ALS risk, particularly in *C9orf72* carriers, suggesting a potential environmental modifier of disease onset.^29–38^ While exercise has been observed as beneficial to astrocytic function, a mechanistic link between activity-dependent astrocyte stress and phenoconversion has not been established.^39^

To address this, we employed direct fibroblast to induced-neural-progenitor-cell (iNPC) reprogramming followed by astrocyte differentiation, a platform that preserves age-associated epigenetic signatures and enables modelling of disease-relevant cell states.^19,40^ We applied this approach to a unique C9-HRE pedigree comprising monozygotic twins discordant for ALS and their asymptomatic father, enabling interrogation of phenoconversion across genetically matched but clinically divergent states. Studies of monozygotic twins have previously been instrumental in demonstrating how environmental exposures and epigenetic variation can shape disease phenoconversion in disorders such as multiple sclerosis, Alzheimer’s disease, schizophrenia, and type 1 diabetes, underscoring the value of this framework for dissecting non-genetic determinants of disease onset.^41,42^

By integrating molecular, electrophysiological, and translatomic analyses across these isogenic astrocyte populations, we identify stage-specific astrocyte dysfunction and demonstrate that astrocyte state transitions, rather than genotype alone, underpin susceptibility to neurotoxicity. Our findings position astrocytes as active determinants of C9-HRE phenoconversion and implicate activity-dependent stress as a potential trigger of disease emergence in genetically predisposed individuals.

## Materials and methods

Detailed methods are provided in the Supplementary material.

### Skin biopsy collection and fibroblast banking

Human skin fibroblast samples were obtained via the AMBRoSIA sampling initiative (Study number STH16573, Ethics Committee reference 12/YH/0330; **Supplementary Table 1**). Informed consent was obtained from all subjects before sample collection.

### Direct conversion of skin fibroblasts into induced neural progenitor cells (iNPCs)

Direct conversion was performed with a modified version of the previously described protocol.^19^ Immunocytochemistry was undertaken staining for Nestin and PAX6 expression to confirm successful iNPC conversion. All experiments were conducted in iNPCs between passage 14-21.

### Differentiation into iAstrocytes

For iAstrocyte differentiation, 200,000 iNPCs were plated in medium composed of DMEM, 10% FBS, 1% P/S, 0.2% N2-supplement in a 10 cm^2^ dish coated with 2.5 µg/mL fibronectin. Differentiation took place over 7 days, with a complete media change on day 3.^19^

### Statistical analysis

Data are presented as mean ± SD (except electrophysiology: mean ± SEM). Statistical comparisons utilised two-sided Wilcoxon rank-sum test, one-way ANOVA, two-way ANOVA and Kruskal-Wallis test (each with appropriate post hoc tests). Translatome data normality was assessed using linear regression modelling, visualised as MA plots. Significance was set at *P* < 0.05.

## Results

### Description of the C9-HRE pedigree with discordant monozygotic twins and their asymptomatic father (Fig. 1)

The affected twin presented at the age of 37 years in November 2014. He had previously been very fit and active, attending a gymnasium daily since the age of 21 and enjoying regular swimming and running activities. He had a past medical history of familial hypercholesterolaemia, well-controlled ulcerative colitis diagnosed 9 years previously and an episode of gout at the age of 30 years. He had a 7-year history of regular calf cramps and a 2-year history of cramps affecting his right hand. This was followed by progressive weakness and wasting of the right hand, with more recent development of a degree of weakness affecting the left hand also. At the time of presentation, he had no symptoms affecting the lower limbs, bulbar or respiratory muscles. He had an identical twin brother and reported no family history of neurological disorders.

**Figure 1.**
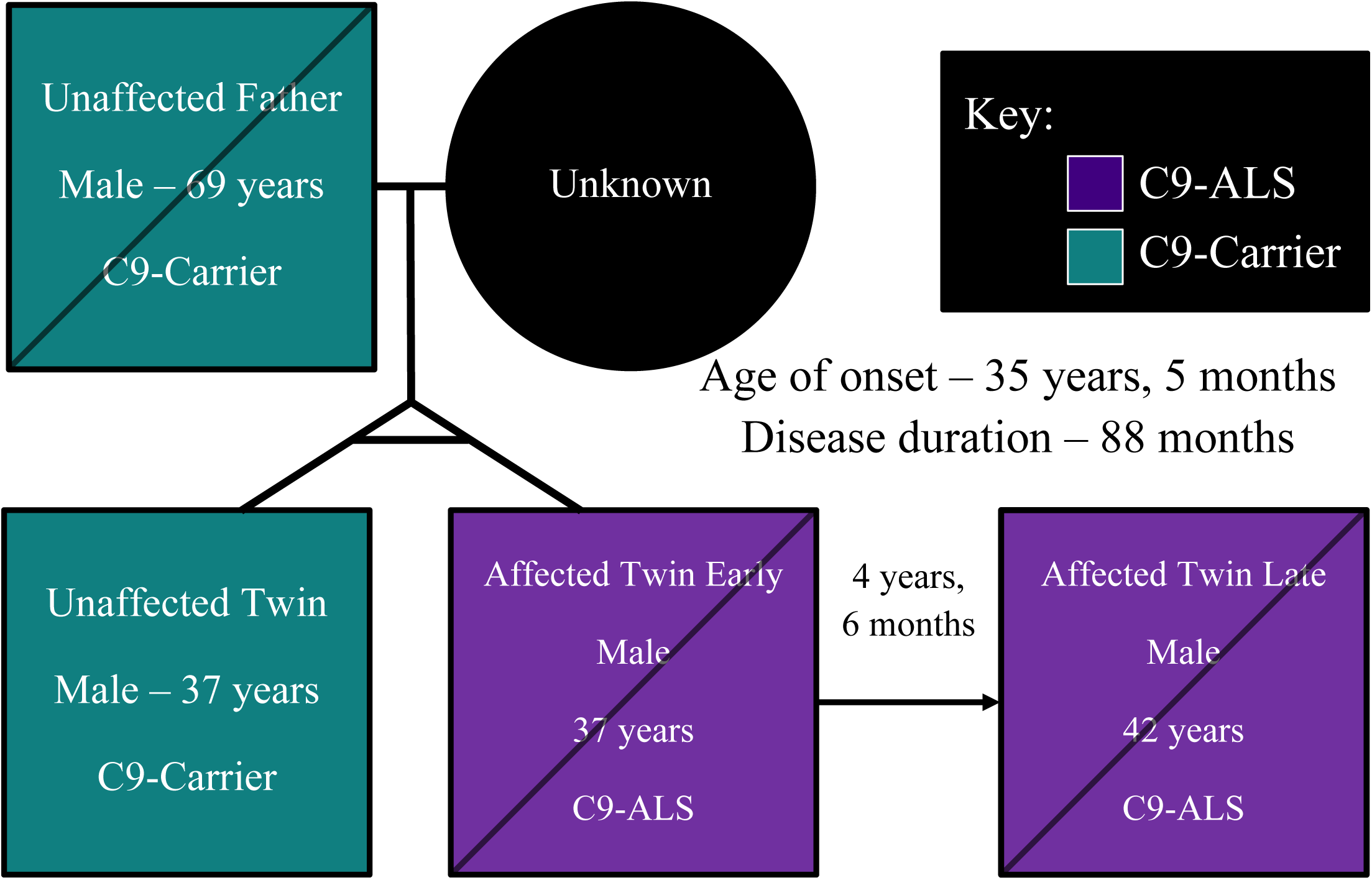
Pedigree of the C9-HRE family. An initial set of biopsies was collected from the proband (affected twin early) 2 years and 4 months after symptom onset, his unaffected twin brother and unaffected father. A second biopsy was collected from the patient (affected twin late) 4 years and 6 months later at the late stage of disease (4 months before death). A familial predisposition to ALS was established on the paternal line, with no such link maternally.

On neurological examination his speech, gait and cognition were normal. There were no abnormalities in the cranial nerve or lower limb territories. In the upper limbs, proximal fasciculations were evident on the right and there was wasting of the right intrinsic hand and forearm muscles, with significant weakness of finger extension and the ulnar and median innervated intrinsic hand muscles. On the left he had mild weakness of abductor pollicis brevis only. There were no clear upper motor neuron signs and clinically the features were those of the progressive muscular atrophy variant of motor neuron disease.

Detailed neurological investigations showed no significant abnormalities on MR imaging of the brain and cervical spinal cord. The initial neurophysiological examination showed diffuse fasciculations in the upper limb, lower limb and abdominal musculature, with active denervation in the muscles of the right upper limb, the left first dorsal interosseous and biceps and the right tibialis anterior. A later neurophysiology examination showed more widespread active and chronic neurogenic changes.

The clinical course was of slowly progressive disability affecting all 4 limbs. He required non-invasive ventilatory support from 2018. He passed away in 2020, 5 years after presenting to neurological attention and ∼7 years after the onset of his symptoms. His identical twin brother remains well 12 years after the onset of his brother’s neurological symptoms. The father of the twins remained neurologically asymptomatic until his death from cancer at the age of 75 years.

Skin biopsies were collected from the unaffected identical twin and the father. The affected twin had an initial biopsy collected 18 months after symptom onset and a second 4 ½ years later at the late stage of disease. The presence of a *C9orf72* repeat expansion was confirmed clinically in the affected twin via repeat primed PCR. We performed repeat primed PCR using material from the asymptomatic biopsies, confirming that both the unaffected twin and father carried C9-HRE expansions.

### The monozygotic twins have identical genetic predisposition for ALS, but different methylation status

First, we assessed whether differences in *C9orf72* repeat length could account for disease discordance within the family. Initially we used Southern blot analysis of fibroblast DNA (**Fig. 2A**). Both monozygotic twins carried a large, expanded allele consistent with a pathogenic repeat size of approximately 800 repeats, alongside a normal allele. In contrast, the asymptomatic father carried a markedly smaller expanded allele, estimated at approximately 70–100 repeats. To obtain higher-resolution repeat sizing, fibroblast DNA was enriched using CRISPR-Cas9 targeting of flanking *C9orf72* sequences and subjected to Oxford Nanopore sequencing. Single-molecule analysis confirmed repeat expansions of ∼800 units in both twins and ∼70 units in the father (**Fig. 2B**). These data indicate a >10-fold intergenerational expansion, which could be consistent with previously reported potential for the *C9orf72* repeat expansion to increase in size across generations.^43,44^

**Figure 2.**
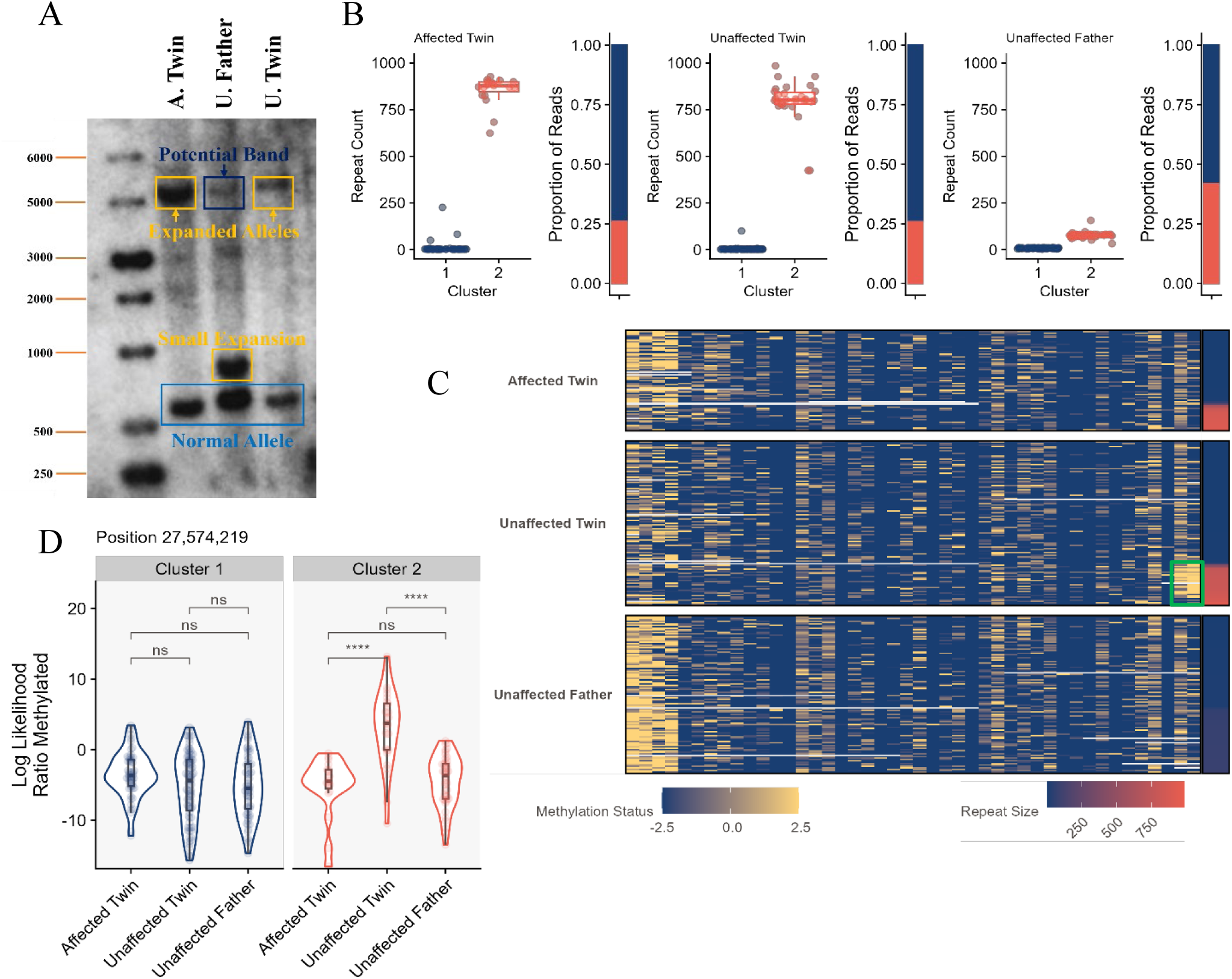
Discordant identical twins carry comparably sized C9-HRE, with hypermethylation observed in the unaffected twin. **A.** *C9orf72* hexanucleotide repeat expansion size was determined using Southern blotting, performed on HMW gDNA extracted from the donated fibroblasts. A large band showing the expansion was clearly seen in both twins (gold boxes), with a fainter band at the same height in the unaffected father’s fibroblasts (dark blue box). A smaller, but still pathologically expanded allele was identified in the father (gold box), while additional normal alleles were observed in all samples (blue box) (A. Twin = affected twin early; U. Father = unaffected father; U. Twin = unaffected twin). **B.** Oxford Nanopore MinIon sequencing was performed after CrispR-Cas9 enrichment to sequence the *C9orf72* repeat expansions in donor fibroblasts. This confirms the result from the Southern blotting (Fig. 2A) with a normal allele in all samples, a small but pathogenic length allele in the father and larger expanded pathogenic alleles in both discordant twins. Two biological repeats per fibroblast line. **C.** Oxford Nanopore MinIon sequencing additionally captures methylation status from each nucleotide passing through the nanopore. This heatmap displays the methylation status for each sequenced DNA strand (blue = unmethylated, yellow = methylated), classified by repeat length (blue = small, red = large). An area of highly methylated DNA was observed in the unaffected twin’s expansion cluster (green box). **D.** Quantification of the methylation status heatmap using violin plotting (Fig. 2C). No significant differences were observed in the methylation status of the normal allele (Cluster 1), though interestingly, both unaffected samples have a greater spread of methylation status, whilst the affected twin has more homogeneous methylation. In the expanded allele (Cluster 2), the unaffected twin has significantly higher methylation than the father and affected twin, with the latter again having a reduced spread of methylation status. Significance test with Two-sided Wilcoxon rank-sum test. ****, *P* < 0.0001.

Together, these analyses demonstrate that the clinically divergent, monozygotic twins harbour equivalent large *C9orf72* repeat expansions, whereas the asymptomatic father carries a substantially smaller expansion, excluding repeat length as a determinant of phenoconversion within the twin pair, but supporting its contribution to reduced disease penetrance in the father.

We next assessed whether epigenetic regulation of the *C9orf72* expansion contributes to phenotypic divergence within the family, focusing on allele-specific DNA methylation as a known modifier of repeat-associated transcription and downstream pathological burden.^45^ To quantify this, we performed Oxford Nanopore sequencing, enabling simultaneous measurement of repeat length and CpG methylation across the expanded locus (**Figs. 2C and D**). This revealed marked hypermethylation of the expanded allele in the unaffected twin compared with both the affected twin and the father (*P* < 0.0001 for both). Methylation changes were allele-specific, with no differences observed in the normal allele across individuals. Thus, the unaffected twin carries an epigenetically silenced *C9orf72* expansion, providing a plausible protective mechanism against downstream molecular pathology, despite an identical repeat length compared to the affected twin.

### Equivalent ALS-Associated Genetic Burden in Discordant Monozygotic Twins

We next asked whether additional known ALS-associated genetic variants could explain the phenotypic discordance between the monozygotic twins. To address this, fibroblasts were directly converted to iNPCs and genomic DNA was extracted for whole genome sequencing (WGS) and targeted analysis using two neurodegenerative gene panels covering 144 ALS-associated loci (**Supplementary Tables 6 and 7**).^19,46,47^ Neither twin carried an additional mutation which has been conclusively linked to ALS.^48,49,50^ These data indicate that the monozygotic twins share an essentially equivalent burden of known ALS-associated genetic risk variants, and that discordant disease expression cannot be explained by identifiable differences in coding genetic variation.

### Astrocyte-mediated motor neuron toxicity increases with disease stage and is present in asymptomatic C9-HRE carriers

Previous studies have demonstrated that *C9orf72* astrocytes exert non-cell-autonomous toxicity to motor neurons.^18–28^ To determine whether this phenotype is present across the clinical spectrum of C9-HRE carriers, we employed an iAstrocyte–motor neuron co-culture system, as previously described, using cells generated from this unique familial pedigree.^19^

Patient fibroblasts were directly converted into iNPCs and differentiated into astrocytes using an established protocol.^19^ Specifically, iAstrocytes were generated from healthy individuals, an unrelated symptomatic C9-HRE individual, asymptomatic father, unaffected twin and the affected twin, where the latter was sampled at an early and late time point in disease progression (**Supplementary Table 1**). The presence of the expanded *C9orf72* allele was retained in iNPCs derived from both twins, whereas no comparable expanded allele was detected in the unaffected father following differentiation (**Supplementary Fig. 1**), confirming preservation of genotype within the model system. In parallel, the specification of *in vitro* iNPCs and iAstrocytes was confirmed by the highly enriched expression of neural progenitor markers PAX6 and Nestin, and astrocyte markers CD44 and Vimentin, respectively (**Supplementary Figs. 2-3**).

To assess non-cell autonomous toxicity, iAstrocytes were co-cultured with healthy control iPSC-derived motor neurons (**Fig. 3A**), and neurotoxicity was quantified by cleaved caspase-3 after 72 hours (**Fig. 3B**). Consistent with previous reports using this approach, iAstrocytes from an unrelated, symptomatic patient with C9-HRE ALS induced marked motor neuron toxicity (413% increase in caspase-3^+^ neurons vs. healthy astrocytes; one-way ANOVA, *P* < 0.0001), validating the assay.^19^ Within the family, astrocytes from the affected twin demonstrated stage-dependent toxicity, increasing from early disease (183.3% increase; *P* = 0.032) to late disease (499% increase; *P* < 0.0001). Strikingly, astrocytes derived from both related asymptomatic carriers were also intrinsically toxic, inducing significant motor neuron death compared with healthy controls (unaffected twin: 211%, *P* = 0.003; unaffected father: 219%, *P* = 0.005).

**Figure 3.**
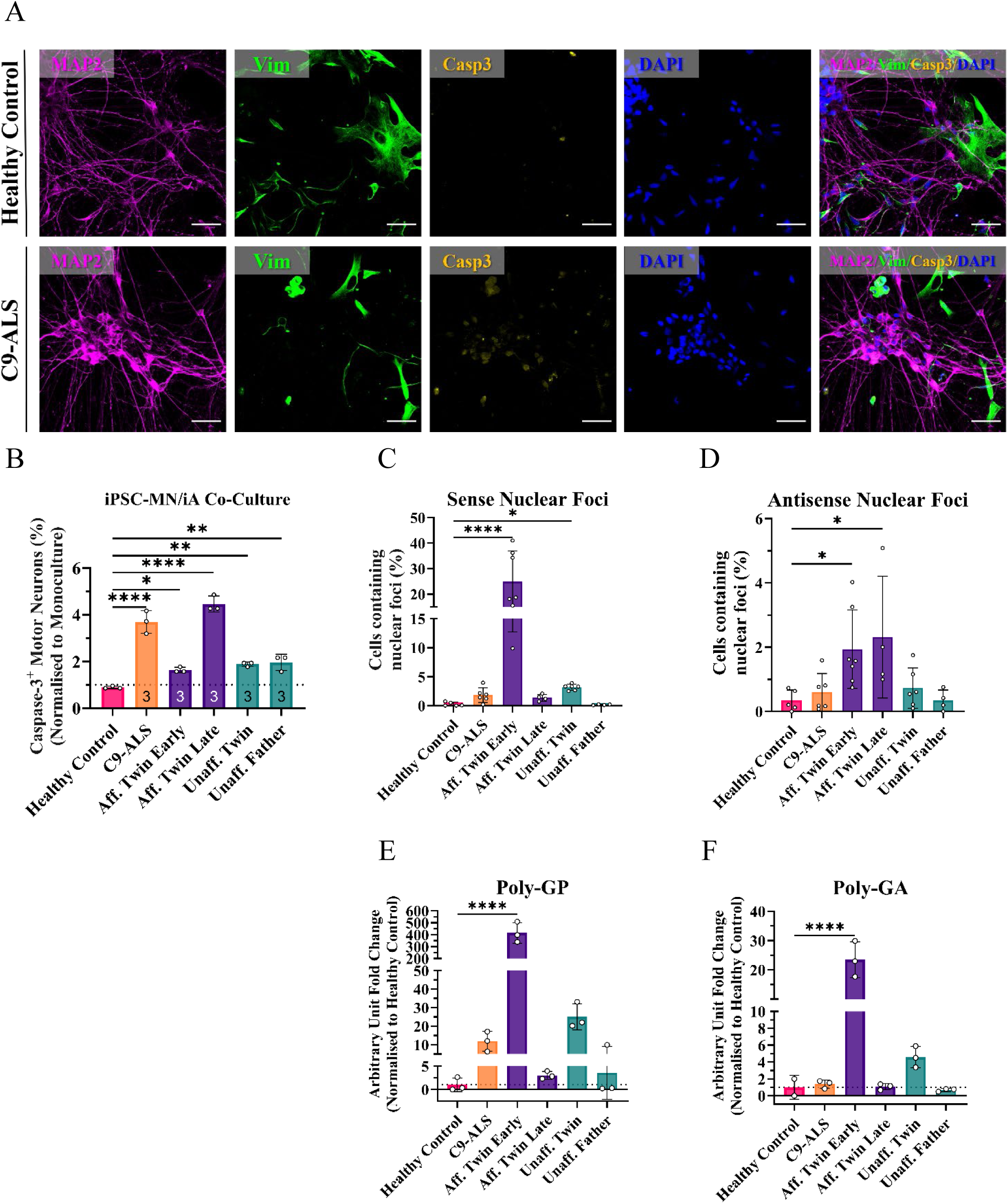
C9-HRE iAstrocytes produce C9orf72 pathological hallmarks and are toxic to motor neurons. **A.** C9-ALS iAstrocytes are toxic when co-cultured with iPSC-derived healthy control motor neurons. Representative images showing MAP2^+^ motor neurons, vimentin^+^ astrocytes and cleaved caspase-3 expression. Cells expressing vimentin were excluded from analysis, with caspase-3^+^ neurons quantified. Images captured on Opera Phenix at 40x magnification. **B.** Cleaved caspase-3^+^ iPSC-derived motor neurons (normalised to monoculture average) after 72 hours of co-culture with iAstrocytes. Significance test with one-way ANOVA with Dunnett’s multiple comparisons test vs. HC. 3 biological replicates per cell line. ****, *P* < 0.0001. **, *P* < 0.01. *, *P* < 0.05. **C&D.** Quantification of iAstrocytes containing nuclear sense (3C) and antisense (3D) RNA foci. Significance test with Kruskal-Wallis test with Dunn’s multiple comparisons test vs. HC. *n* = 4-7 per line per condition. ****, *P* < 0.0001. *, *P* < 0.05. **E&F.** Quantification of poly-GP (3E) and poly-GA (3F) DPRs detected by ELISA. Fold change of arbitrary units normalised to average of Healthy Control. Significance test with one-way ANOVA with Dunnett’s multiple comparison tests vs. HC. *n* = 3 per line per condition (except poly-GA HC – *n* = 2). ****, *P* < 0.0001.

Together, these data demonstrate that astrocyte-mediated motor neuron toxicity is not restricted to symptomatic disease but is present in genetically at-risk individuals prior to clinical onset and increases with disease progression, consistent with progressive amplification of a pre-existing toxic astrocyte state. This contrasts with iAstrocytes derived from neurologically normal individuals which are non-toxic to neurons in co-culture.^19^

### Symptomatic astrocytes display canonical C9orf72 pathology

To investigate whether these features are associated with pathology, fluorescent in situ hybridisation (FISH) was used to assess the production of *C9orf72* RNA foci in the astrocytes from this pedigree (**Supplementary Fig. 4**). Nuclear foci composed of G_4_C_2_-repeat RNA transcribed in the sense direction (hereafter referred to as ‘sense foci’) were detected in both twins, with significant upregulation in astrocytes generated from the affected twin early in disease (Kruskal-Wallis test vs. HC, *P* < 0.0001), with decreased expression later in disease and a smaller, but still significant (Kruskal-Wallis test vs. HC, *P* < 0.05), upregulation in the unaffected twin (**Fig. 3C**). Astrocytes from the unaffected father were comparable with astrocytes from healthy controls. Nuclear G_2_C_4_ foci transcribed in the antisense direction (hereafter referred to as ‘antisense foci’) were also detected.

Astrocytes from both early and late biopsies of the affected twin produced significantly more antisense foci than healthy controls (Kruskal-Wallis test vs. HC, both *P* < 0.05), with both asymptomatic lines comparable to the healthy control baseline. Overall, antisense foci were detected less readily than sense foci (**Fig. 3D**).

We next assessed RAN translation with poly-GP and poly-GA DPR species detectable above baseline using a high-sensitivity MSD ELISA (**Figs. 3E and F**). A similar distribution was seen for both DPRs with significantly higher production observed in the affected twin early biopsy when compared to healthy controls (one-way ANOVA, *P* < 0.0001), but this production was reduced to baseline levels in the late biopsy. The unaffected twin had a higher production than the affected twin’s late biopsy, whilst the levels of DPRs in astrocytes from the unaffected father were comparable to healthy controls. The larger C9-HRE size carried by the unaffected twin may result in the increased production of RNA foci and DPRs when compared to his father, though the reduced expression when compared with the affected twin early astrocytes may be a result of the methylated C9-HRE (**Fig. 2D**). The presence of these pathological hallmarks prior to onset of symptoms (and early in disease) suggests that these features are amongst the earliest changes in the pathophysiology of ALS.

Interestingly when compared to the C9-specific pathological hallmarks, the affected twin has a negative correlation whereby reduced production of RNA foci and DPRs correlated with increased astrocyte toxicity towards co-cultured motor neurons.

### Connexin dysfunction emerges with symptomatic disease and tracks with astrocyte toxicity

Given the critical role of astrocyte membrane physiology in neuron–astrocyte communication and toxicity, we next examined whether altered membrane properties of astrocytes contribute to MN toxicity in ALS.^51^ To examine membrane properties, we performed whole-cell voltage-clamp recordings on individual iAstrocytes, differentiated for 14 days. By evoking changes in membrane potentials, we induced membrane currents in the iAstrocytes, which displayed the expected linear, passive current–voltage relationship characteristic of astrocytes.^52^ However, symptomatic C9-ALS iAstrocytes exhibited markedly larger membrane currents compared to iAstrocytes obtained from healthy individuals. The affected twin iAstrocytes similarly displayed markedly elevated currents, also increasing with disease stage. However, iAstrocytes from the father and unaffected twin showed small currents comparable to the healthy control astrocytes (**Fig. 4A**). Symptomatic iAstrocytes present with a clear membrane dysfunction phenotype. To quantify this, current density analysis confirmed a progressive increase in membrane conductance with disease stage (**Figs. 4B and C**). To rule out increased membrane currents due to loss of membrane integrity, we measured resting membrane potentials. These were determined to be preserved or more hyperpolarised than controls (**Fig. 4D**), indicating that increased currents therefore reflected altered membrane channel function. Together, these data define a disease-associated increase in astrocytic membrane conductance that emerges with symptomatic onset and progresses with disease stage.

**Figure 4.**
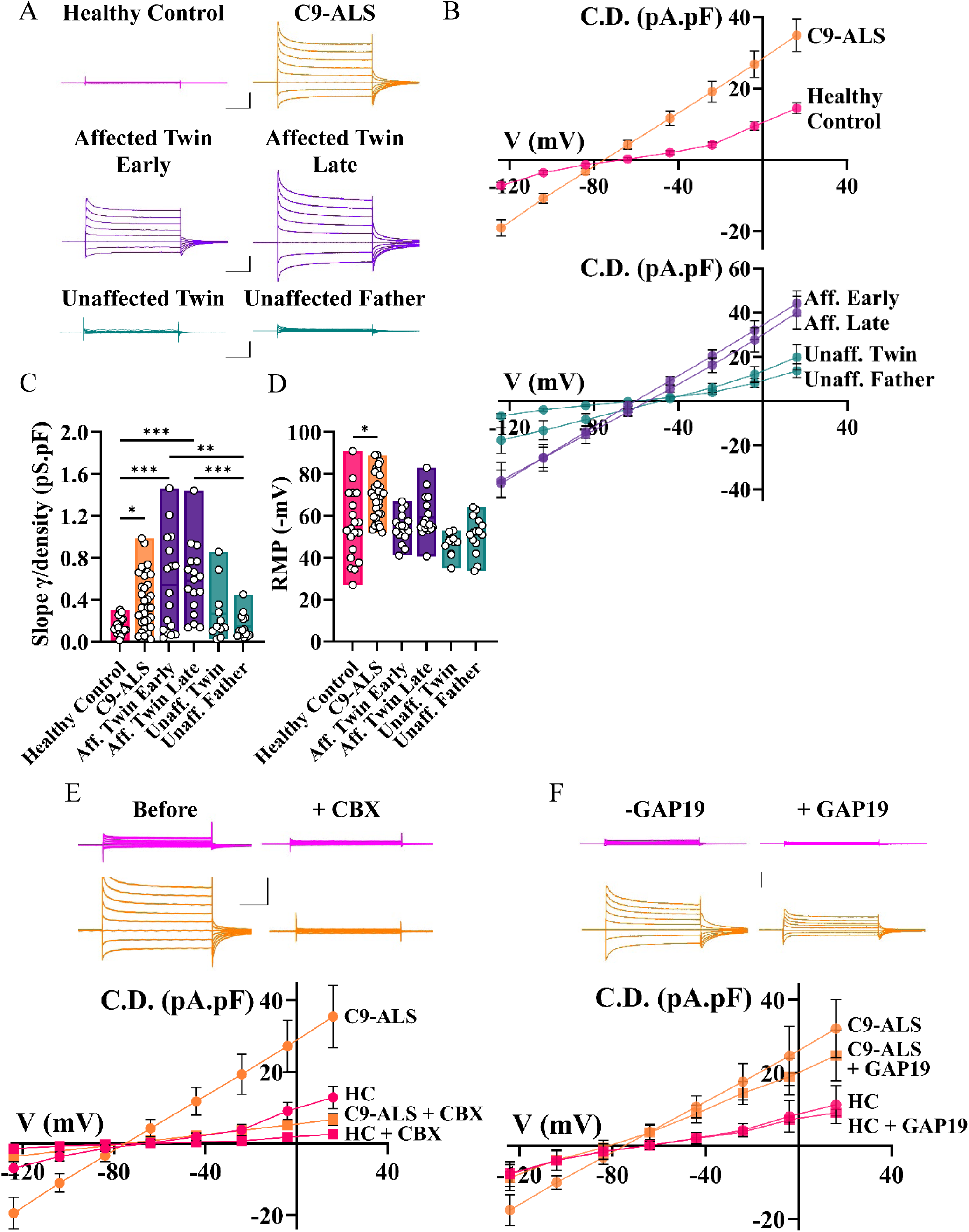
Symptomatic C9-ALS iAstrocytes display enhanced connexin-mediated membrane currents that increase with disease progression. **A.** Individual iAstrocytes were examined by whole cell patch-clamp in the voltage-clamp configuration. Cells were held at a potential of −84 mV and a voltage-step protocol (stepped voltages from −124 mV to +16 mV; 200ms steps applied every 10s) to evoked passive membrane current responses. Representative traces show enhanced membrane currents in the unrelated C9-ALS patient and affected twin. Scale bar; 50ms, 1 nA. **B&C.** Mean ± SEM current density (current normalised to whole cell capacitance) versus voltage for each iAstrocyte background (4B). Slope γ/density was determined from the linear fit for each cell from these plots (4C). Note the increase in membrane conductance in the C9-ALS and affected twin iAstrocytes. **D.** Mean ± SEM resting membrane potential (RMP) of iAstrocytes. The RMP is not depolarised in C9-ALS astrocytes indicating no loss of membrane integrity. **E.** Recordings, performed as in Fig. 4A, before and after the addition of connexin blocker carbenoxolone (CBX) (10 μM) for healthy and C9-ALS iAstrocytes. The large current dysfunction in C9-ALS astrocytes is removed by CBX and quantified in the mean ± SEM current density versus voltage plots below. **F.** Recordings from healthy iAstrocytes and C9-ALS in the absence or presence of Cx43 blocker Gap19 within the intracellular patch pipette solution. Data obtained from 11-32 cells, from >3 de novo differentiations. Significance test with one-way ANOVA with Tukey’s post hoc test. ***, *P* < 0.001. **, *P* < 0.01. *, *P* < 0.05.

To identify the underlying mechanism of membrane dysfunction, we assessed connexin (Cx) channels. Connexin 43 (Cx43), the predominant astrocytic connexin, is upregulated in ALS and its inhibition confers neuroprotection.^53,54^ Application of the connexin blocker carbenoxolone (CBX, 10 µM) completely abolished the enhanced currents in C9-HRE iAstrocytes, restoring current density–voltage relationships to control levels (**Fig. 4E**). Application of the Cx43-selective hemichannel partial blocker Gap19 (100 µM) to the patch pipette solution produced a reduction in current density-voltage-relationship (**Fig. 4F**), consistent with Cx43 hemichannel involvement.^55,56^ These data are consistent with the membrane dysfunction being driven by Cx43 hemichannel dysfunction.^57^ Critically, however, the notable lack of membrane dysfunction in the unaffected C9-HRE twin and father means that the C9-HRE cannot be the sole driver of the membrane dysfunction.

Astrocytic Cx43 has been highlighted as a key mediator of MN toxicity.^53,54^ Cx43 channels are highly permeable to several intracellular molecules including ATP and reactive oxygen species (ROS), which are toxic to MNs at high levels. To test whether the enhanced Cx43 hemichannel currents contributed to toxicity, astrocytes treated with either CBX or the licenced neuroprotective agent riluzole (RIL) were co-cultured with HB9-GFP^+^ mouse motor neurons over the course of 72 hours. CBX-treatment alone had a negligible effect on motor neuron survival, contrasting with reduced toxicity following RIL-treatment, particularly in the highly toxic affected twin late astrocytes. As neither treatment reduced survival, we tested both in combination, observing a synergistic effect, with a significant rescue when affected twin late astrocytes were treated with each of CBX (30 µM) and RIL (30 µM) compared with RIL (30 µM) alone (**Supplementary Figs. 5-7**). When compared with unaffected twin astrocytes, affected twin early (Two-way ANOVA, *P* = 0.02) and late astrocytes (*P* = 0.06) responded with reduced toxicity when treated with RIL (30 µM). This rescue effect was enhanced further in affected twin late astrocytes when co-treated with CBX (30 µM) and RIL (30 µM) when compared with unaffected twin (*P* = 0.0004) or affected twin early astrocytes (*P* = 0.008) (**Figs. 5A and B**). The observed rescue correlates with the level of Cx43 membrane current impairment (**Fig. 4A**).

**Figure 5.**
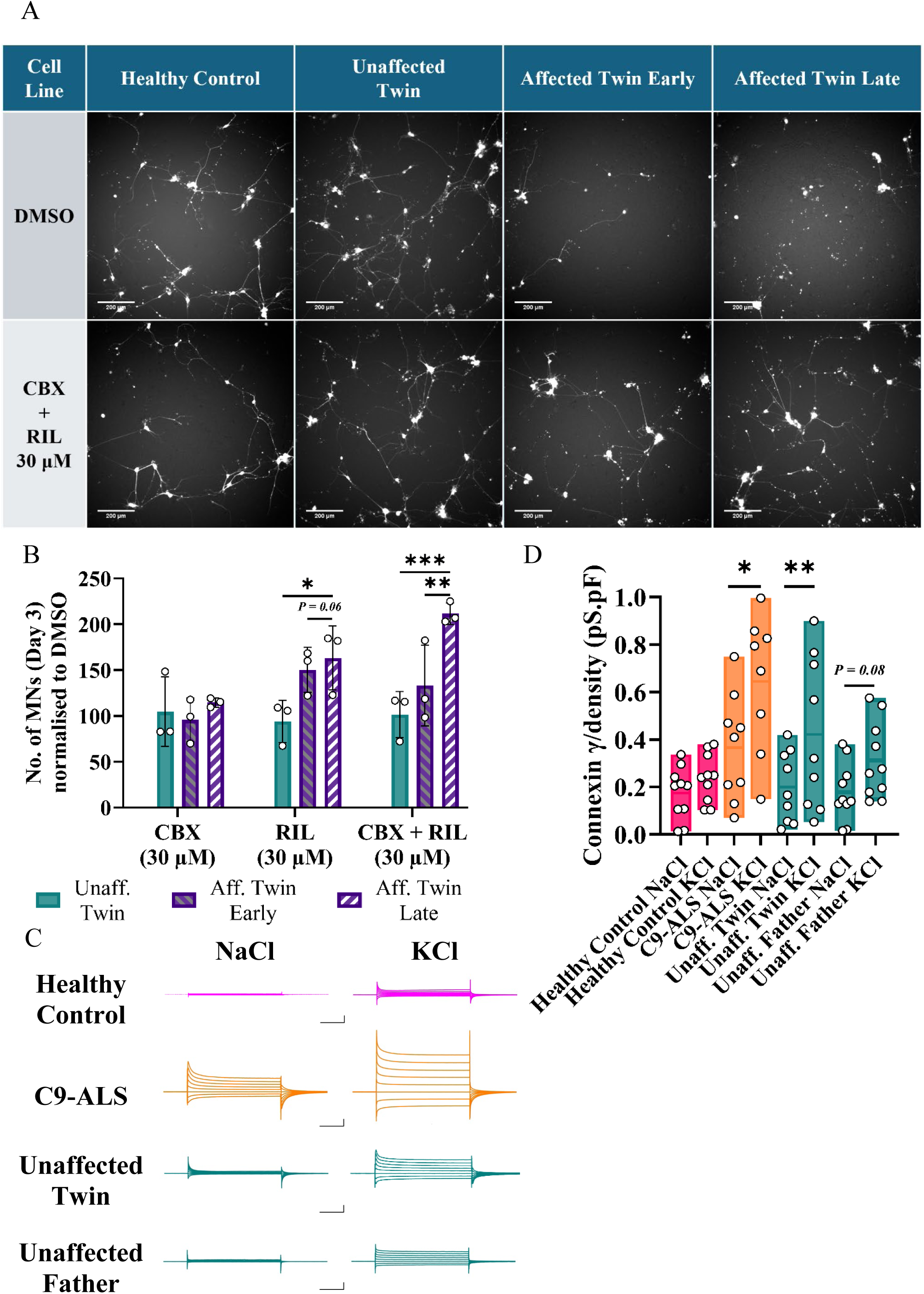
Rescue of astrocyte toxicity to MNs via carbenoxolone and riluzole in combination. **A.** Representative images of HB9-GFP^+^ MN following co-culture with iAstrocytes treated with either DMSO (top) or 30 μM CBX and 30 μM RIL in combination. Motor neurons cultured with healthy control or unaffected twin iAstrocytes are supported, with no toxicity observed following treatment. Motor neurons cultured with affected twin iAstrocytes are reduced in the DMSO baseline and are rescued with CBX and RIL co-treatment. **B.** Twin iAstrocytes were treated with either 30 μM CBX (left), 30 μM RIL (middle), or both in combination (right). When compared with the DMSO baseline, minimal rescue was observed in the CBX-treated iAstrocytes. Motor neuron survival was increased in RIL-treated iAstrocytes from both biopsies of the affected twin. The largest rescue was observed in the affected twin late biopsy iAstrocytes when treated with both carbenoxolone and riluzole in combination. Significance test with two-way ANOVA with Tukey’s multiple comparisons test per treatment. *n* = 3 per line per condition. ***, *P* < 0.001. **, *P* < 0.01. *, *P* < 0.05. **C.** Depicts connexin (CBX-sensitive)-mediated currents from individual iAstrocytes obtained by subtracting CBX-insensitive currents from the total overall current for each cell (see Fig. 4E). The connexin current after the incubation with KCl is highly pronounced versus the NaCl osmotic control. Scale bar; 50ms, 1 nA. **D.** Mean ± SEM connexin γ/density for each iAstrocyte background after KCl or NaCl incubation. Note the pronounced increase in the presence of KCl versus NaCl osmotic control. Data obtained from 8-10 cells, from 3-4 de novo generations of cells. Significance test with Students t-test. **, *P* < 0.01. *, *P* < 0.05.

Collectively, these data identify progressive Cx43-dependent membrane dysfunction as a hallmark of symptomatic disease onset and a functional driver of astrocyte-mediated neurotoxicity in *C9orf72-*associated ALS.

### Discrepancy in life-style factors between the twins: levels of strenuous physical activity

Quantification of historical leisure-time physical activity in the twins, using the previously validated HAPAQ questionnaire (Supplementary Methods), revealed a significant difference between energy expenditure in the affected and unaffected twin. Physical activity in the 10 years prior to the age of symptom onset in the affected twin was >4-fold higher compared to the unaffected twin (average leisure-time activity 13.70 kJ/kg/day versus 3.06 kJ/kg/day). Previously we have linked physical activity in early adulthood to the age of ALS onset. Performing a similar analysis here reveals a substantially higher level of physical activity in the affected twin (19.51 kJ/kg/day versus 1.53 kJ/kg/day).^58^

### Mimicking the effects of physical exercise in cultured astrocytes

Astrocytes carrying the C9-HRE have been observed to show reduced metabolic flexibility, potentially limiting the neuroprotective response to the increased synaptic activity during exercise.^59^ Exposing these genetically predisposed astrocytes to intense exercise may hasten the development of ALS disease phenoconversion.^60^

Physical exercise is associated with adaptive remodelling of astrocyte physiology, reflecting a homeostatic response to heightened central nervous system demand.^60^ During periods of intense neuronal activity and relative hypoxia, both elevated during exercise, expression of Cx43 is increased.^61^ One proposed mechanism underlying this response is enhanced extracellular K⁺ buffering due to increased neuronal K+ extrusion, where connexin channels facilitate spatial K⁺ redistribution, limiting periaxonal K⁺ accumulation, thus preventing sustained depolarization and reducing the risk of pathological hyperexcitability.^62^ However, given that elevated Cx43 expression is toxic, we hypothesised that C9-HRE astrocytes may display an abnormal dysregulation of connexins in response to elevated extracellular K+, modelling increased neuronal activity.^51,53,62^ To model sustained elevations in extracellular K⁺ associated with high levels of neuronal activity, we exposed iAstrocytes derived from healthy controls, C9-HRE carriers and unaffected family members to 30 mM KCl for 72 hours (osmotic control: 30 mM NaCl). Consistent with a homeostatic response, KCl treatment induced a modest increase in connexin (CBX-sensitive) membrane current density in healthy iAstrocytes. In contrast, connexin current density was markedly elevated in C9-HRE iAstrocytes under the same conditions. Strikingly, unaffected family member iAstrocytes also exhibited a pronounced upregulation of connexin-mediated current density following KCl exposure compared to the osmotic NaCl control (**Figs. 5C and D**). Taken together, these findings demonstrate that elevated extracellular K⁺ unmasks a dysregulated connexin-mediated response in C9-HRE astrocytes and suggests that connexin dysfunction can be precipitated or accelerated by aberrant homeostatic signalling under conditions of increased neuronal demand, such as during strenuous exercise.

### Single-subject translatome sequencing analysis reveals impairments in cellular homeostasis

To elucidate the mechanisms behind these divergent phenotypes, iAstrocytes were subjected to translatome RNA-sequencing (**Fig. 6A**).^63^ Expression of transcripts was assessed between sample pairs as MA plots, the linear regression R^2^ values explaining less than 1% of the variation, showing comparable distribution in each (**Supplementary Table 8, Supplementary Figs. 8-10**). To compare between these genetically identical samples we utilised N-of-1-*pathways* analysis, developed to assess dysregulation in cancer patients by comparing tissue from tumours with that sequenced from a comparatively healthy tissue in the same subject.^64,65^

**Figure 6.**
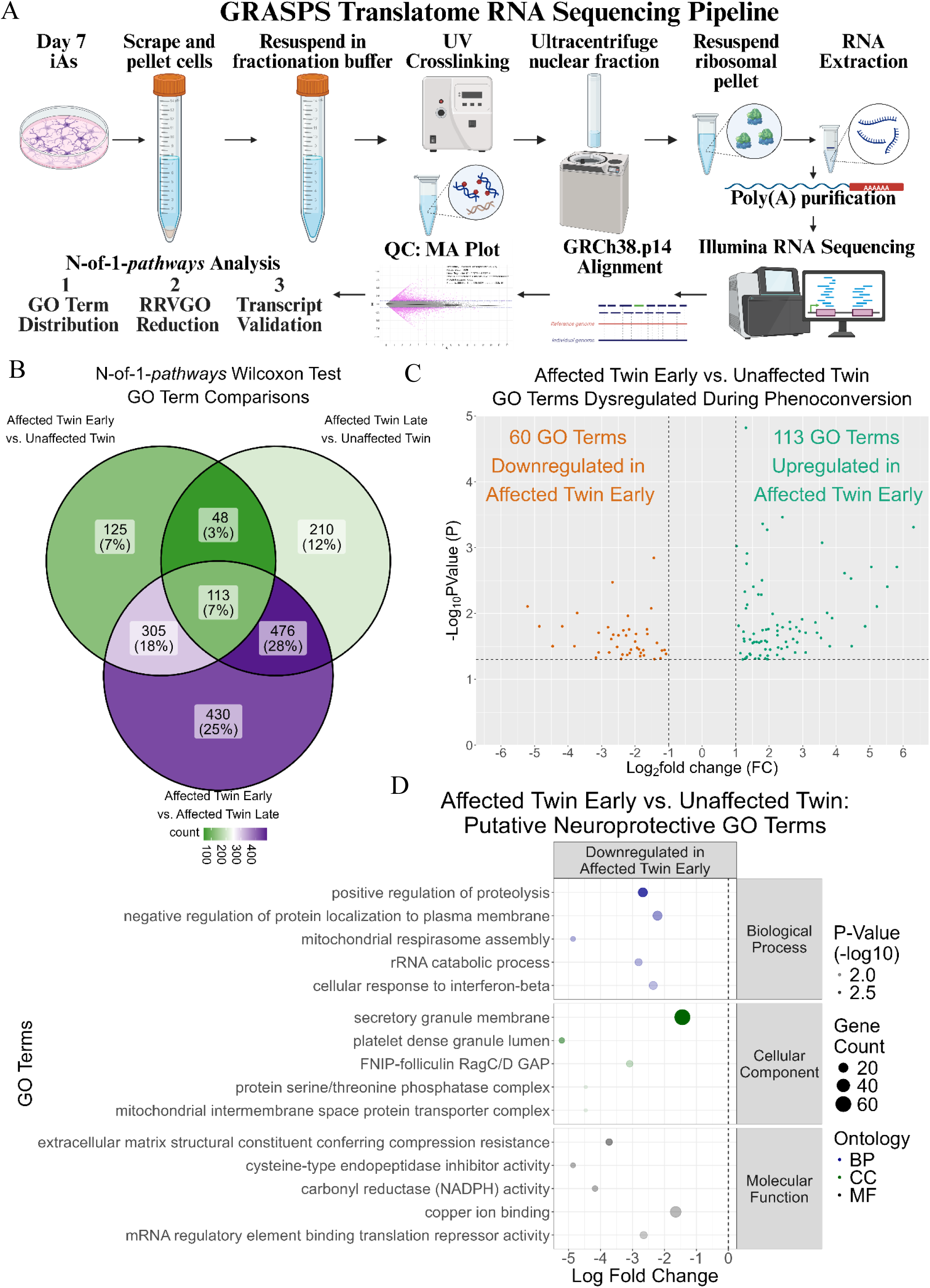
Identification of neuroprotective genes weakened during phenoconversion. **A.** Schematic of GRASPS Translatome technology and subsequent bioinformatics analysis pipeline via Illumina RNA sequencing and N-of-1-*pathways* single subject analysis.^50–52^ Generated using Biorender. **B.** Comparison of significantly dysregulated GO clusters identified with N-of-1-*pathways* Wilcoxon Test. 78% of the identified GO terms were found in the affected twin early vs. affected twin late enrichment. 7% were common to all three comparisons, a further 7% unique in the affected twin early vs. unaffected twin enrichment and 12% in the affected twin late vs. unaffected twin enrichment, with 3% common between these two comparisons. **C.** Volcano plot of GO clusters dysregulated between the affected twin early and unaffected twin during phenoconversion, comprising 125 GO terms uniquely dysregulated and a further 48 common with the comparison between the affected twin late and unaffected twin as seen in Fig. 6B. 113 GO terms were upregulated in affected twin early and 60 downregulated. **D.** GO Terms downregulated in affected twin early in Fig. 6C were hypothesised as potentially neuroprotective. The five most significant terms from each gene ontology were arranged by increasing significance (-log10 *p*-value). The five most significantly dysregulated GO terms overall were assessed further with contributing genes extracted in Supplementary Fig. 17.

Firstly, the asymptomatic and symptomatic samples were compared. 168 significantly upregulated and 426 downregulated GO terms were observed in the affected twin early compared to unaffected twin, while 809 significantly upregulated and 43 downregulated GO terms were observed in the affected twin late compared to unaffected twin (**Supplementary Figs. 11-12**). When comparing between the symptomatic samples, 114 significantly upregulated and 1221 downregulated GO clusters were observed in affected twin early compared with the affected twin late astrocytes (**Supplementary Fig. 13**). GO clusters enriched in the unaffected twin related to translation, proteasome complex and GTP binding (vs. affected twin early), and vacuolar acidification, proton transport and transmembrane signalling (vs. affected twin late) (**Supplementary Figs. 14-15**). GO clusters upregulated in the affected twin early related to synapse assembly, cell-cell junction and plasma membrane (vs. unaffected twin), and sphingolipid, plasma membrane and transmembrane signalling receptor activity (vs. affected twin late) (**Supplementary Figs. 14 and 16**). Finally, the affected twin late, containing the most abundant and significantly dysregulated clusters, was enriched for translation and DNA repair (vs. affected twin early), and DNA regulation and apoptosis (vs. unaffected twin) (**Supplementary Figs. 15-16**).

When the identified GO terms were compared, 7% were common to the three comparisons, with 25% unique between affected twin early vs. affected twin late, 12% unique between affected twin late vs. unaffected twin and 7% (125) unique between affected twin early vs. unaffected twin (**Fig. 6B**). Overall, the observed pattern of enrichment and the downregulation of translation-related terms in the affected twin early, suggests that an impairment in translation occurs between the asymptomatic and early symptomatic phases, which is then reversed and exacerbated by the end stage of disease. A reduction in canonical translation could explain the increase in non-canonical RAN translation resulting in increased DPR production in this early-stage disease sample (**Fig. 3D**).

To investigate if the translation reduction was due to impairment in the translation initiation machinery, elongation factor transcript counts were assessed. The master regulator eIF2α, which is phosphorylated to inhibit translation, had reduced expression in affected twin early. Four kinases (HRI, PKR, PERK and GCN2) are responsible for the phosphorylation of eIF2α, each in response to a different stimulus.^66^ When compared to the average of the unaffected and affected samples, PKR was reduced by ∼20% in affected twin early, HRI was comparable to the average, with PERK and GCN2 elevated. This suggests that eIF2α phosphorylation is not inhibited, to explain translation repression in affected twin early (**Supplementary Table 9**).

Next, we aimed to identify if the electrophysiological membrane impairment was reflected in the translatome. GO terms for cation channels, potassium ion homeostasis and gap junctions were used to develop a relevant gene list (**Supplementary Table 10**). Of the five most abundant transcripts in the dataset, four were upregulated in the affected twin early, with the largest upregulation observed in Cx43 (216% upregulated compared to unaffected twin and 166% upregulated compared to affected twin late). This increase from asymptomatic to symptomatic astrocytes supports the increased impairment previously identified (**Fig. 4A**). Interestingly, the second most abundant transcript in this list, CCN3, had negligible expression in the affected twin early (0.1% of the average). CCN3 is a matricellular protein expressed by astrocytes as part of the extracellular matrix, regulating cell adhesion, migration, differentiation and survival. CCN3 has been shown to play a role in inflammation by upregulating CCL2 and CXCL1 and as an interaction partner of Cx43.^67,68^

Finally, we looked to identify novel neuroprotective targets using the GO terms uniquely dysregulated between the unaffected twin and affected twin early, together with those that overlapped with the comparison between unaffected twin and affected twin late (**Fig. 6B**). The 173 GO terms were split, with 113 upregulated in affected twin early and 60 downregulated in affected twin early (**Fig. 6C**). The downregulated GO terms (i.e. upregulated in the unaffected twin) were assessed further as putative neuroprotective ontologies. These most significant of these GO terms included regulation of proteolysis and protein localization to plasma membrane (Biological Process), platelet dense granule lumen and secretory granule membrane (Cellular Component) and extracellular matrix structure (Molecular Function) (**Fig. 6D**). The contributing genes from these five GO terms were extracted and 75 transcripts with negative Log2FC (i.e. increased in unaffected twin) were taken forward (**Supplementary Table 11, Supplementary Fig. 17**).

Overall, this network of transcripts upregulated in the unaffected twin is related to protein homeostasis, via the mechanisms of degradation, transportation, lysosomal function and cellular remodelling in response to stress. This may represent compensatory neuroprotective mechanisms allowing for controlled response to and turnover of pathological hallmarks. In contrast, the impairment of these processes could result in accumulation of harmful inclusions, such as the RNA foci and DPRs observed in the affected twin early (**Figures 3C and D**).

To investigate this hypothesis of an impairment in protein degradation, we assessed TDP-43 mislocalisation using immunocytochemistry. In the healthy control astrocytes, ∼2% of cells contained TDP-43 cytoplasmic inclusions following blinded manual quantification. Each of the cell lines carrying the C9-HRE had increased levels of TDP-43 inclusions, with the highest burden observed in unaffected twin astrocytes (13.6%) and the lowest in affected twin late (2.9%). 7.5% of affected twin early astrocytes contained TDP-43 inclusions, with <5% in the unaffected father (who did not develop disease before death) and C9-ALS (66 years old at biopsy collection, within end-stage of disease) astrocytes. Together, these data suggest that TDP-43 aggregation occurs throughout the disease course from pre-symptomatic to end-stage, though interestingly, at a reduced rate as the disease progresses (**Supplementary Table 12**). Furthermore, this increase in TDP-43 aggregation does not correlate with non-cell autonomous toxicity, with the two most toxic lines (affected twin late and unrelated C9-ALS) having lower aggregation, resulting in a non-significant negative correlation across the C9-HRE^+^ lines (**Supplementary Figure 18**).

## Discussion

Using direct conversion of fibroblasts, we have generated a cohort of iNPCs from biopsies representing the C9-ALS disease continuum from asymptomatic, early symptomatic, late symptomatic and finally, a non-penetrant *C9orf72* carrier. The differential disease-specific phenotypes from the three genetically identical biopsies expand on our previous observation that these directly converted iNPCs retain the epigenetic features of ageing.^40^

Previous studies of monozygotic twins discordant for ALS have identified differences in methylation suggesting that affected individuals undergo accelerated ageing during disease development.^69,70^ What is less clear is whether this acceleration acts as a driver of disease (i.e. following an environmental stimulus) or represents a consequence of disease development. While most studies have assessed twin discordance using DNA isolated directly from biosamples (usually whole blood), iPSCs have been derived from monozygotic twins discordant for sporadic ALS and differentiated into motor neurons.^71^ However, the observed phenotypes did not significantly differ between these twins, likely due to the epigenetic resetting that occurs during iPSC-reprogramming.^72^

As expected, both monozygotic twins in the present study carried a C9-HRE, the size of which was confirmed to be comparable by Southern blotting and Nanopore sequencing at 800 repeats. Interestingly, the unaffected father carried a much smaller repeat size of only 70 repeats, larger than the current at-risk threshold of 30 repeats, but not sufficient to induce disease at the time of biopsy collection. The continued challenge of accurately sizing the C9-HRE, including somatic mosaicism, has hindered sufficiently powered studies to understand the rate and mechanisms leading to inter-generational increases in expansion length. There is greater consensus on anticipation whereby age of onset decreases with subsequent generations.^44,73,74^ Our data support this consensus and warrant further study in additional pedigrees to confirm if increasing repeat length is a mechanism leading to disease anticipation.

Assessing methylation at genomic level was beyond the scope of this study, but hypermethylation of the expanded C9-HRE was observed in the unaffected twin. This potentially protective mechanism may explain the reduced expression of C9orf72 pathological hallmarks observed in the unaffected twin’s astrocytes.^45^

In addition to the C9-HRE, a further seven variants of uncertain significance (VUS) were identified in the twin’s iNPCs, with five shared with their father. Further experimentation would be required to assess any functional effect on disease development, but importantly for this study, the identical inheritance in the discordant twins confirms comparable genetic predisposition to ALS. This suggests the effect of additional environmental factor(s) resulting in the observed discordance.

In the original study of 100 patients using the ALS/FTD panel, 21% of patients carried a variant conferring pathogenic or likely pathogenic risk, a further 21% carried a VUS and 13% carried more than one variant (including VUS).^46^ This latter group had a significantly earlier age of onset compared with patients carrying a single variant. Of note, C9-HREs were identified in 10 patients, with four of those individuals carrying an additional variant (40%). These data, combined with the VUS identified in the twin pedigree, suggest that polygenic risk burden may be more common in C9-HRE carriers, potentially contributing to the more aggressive nature of C9-ALS.

In iAstrocytes, sense RNA foci were more readily detected than antisense, with increasing disease burden correlating with increased detection of sense DPRs poly-GP and poly-GA. Interestingly, this contrasts with motor neurons and microglia, both observed with more widespread expression of antisense foci.^75,76^

When assessing the motor neuron toxicity of lines with comparable C9-HRE length, increasing toxicity was negatively correlated with the abundance of RNA foci and DPR burden. In the clinical trial of BIIB078, an antisense oligonucleotide targeting sense C9orf72 repeat RNA, poly-GP was reduced but this did not lead to clinical benefits or reduction in TDP-43 pathology.^77^

This observed dynamism of sense RNA foci and DPRs may indicate a role in disease development at the prodromal and early symptomatic stages of ALS. Importantly, DPR species from the antisense C9orf72 repeat RNA continue to be challenging to detect. While this may be due to reduced abundance – inferred from the reduced expression of antisense RNA foci in this astrocyte model – differences in expression may contribute to astrocyte toxicity to motor neurons.

Connexin-mediated membrane dysfunction was minimal in iAstrocytes derived from asymptomatic individuals but became pronounced in symptomatic iAstrocytes and increased with disease progression. This stage-dependent phenotype, observed in monozygotic twins carrying a comparable C9-HRE, indicates that the mutation alone is insufficient to drive connexin dysregulation and instead supports its emergence as a feature of symptomatic disease onset. The absence of a comparable phenotype in genetically identical asymptomatic carriers suggests that additional modifiers, potentially epigenetic or environmental, are required to facilitate this transition. Consistent with this, directly converted astrocytes retain age-related epigenetic signatures. Accelerated epigenetic ageing and altered neuronal and immune signalling have been reported in discordant ALS twins, providing a plausible mechanism underlying phenoconversion.^73,78,79^

The functional upregulation of Cx43 hemichannels provides a direct route by which astrocytes can acquire a neurotoxic phenotype. Connexin channels are permeable, not only to ions, but also to a wide range of cytoplasmic factors, including ATP, glutamate, reactive oxygen species (ROS) and inflammatory mediators such as TNF-α, IL-1β and IL-6. While gap junctions normally support intercellular homeostasis within the glial syncytium, pathological opening of Cx43 hemichannels enables the release of these factors into the extracellular space. This results in accumulation of neurotoxic mediators, particularly glutamate and ATP, driving excitotoxicity, an established pathophysiological feature of ALS, and contributing to motor neuron injury and cell death.^80–83^ In parallel, connexin dysfunction has been associated with reduced expression of glutamate transporter GLT-1, further impairing glutamate clearance and exacerbating synaptic toxicity.^84^

Beyond mediator release, Cx43 upregulation may also contribute to metabolic dysfunction in ALS iAstrocytes. Connexins have been implicated in mitochondrial coupling, a process that enhances ATP production but also increases ROS generation.^85^ Elevated Cx43 activity may therefore promote both the production and release of toxic metabolites, linking astrocyte metabolic stress to motor neuron injury. This is consistent with evidence that Cx43 is upregulated under inflammatory conditions and drives pathological calcium signalling and excitotoxicity in ALS models, as well as with the broader metabolic and oxidative stress signatures observed in ALS astrocytes.^53,54,86^

These findings support a model in which astrocyte connexin dysfunction represents a key pathological transition associated with symptomatic disease onset. In this framework, Cx43 hemichannel upregulation does not initiate disease but amplifies pathology by coupling astrocyte metabolic, inflammatory and excitotoxic mechanisms to motor neuron degeneration. This positions connexin dysfunction as both a marker of phenoconversion and a potential therapeutic target for modifying disease progression in ALS.

The marked difference in lifetime strenuous physical activity between the monozygotic twins provides a compelling environmental factor that may contribute to disease phenoconversion. The affected twin exhibited substantially higher levels of sustained physical activity, consistent with epidemiological evidence linking strenuous exercise to increased ALS risk and earlier onset, particularly in *C9orf72* expansion carriers.^33^ These observations suggest that heightened neuronal activity during strenuous physical activity may act as a critical environmental modifier in genetically predisposed individuals.

Neuronal activity is tightly coupled to astrocyte function through extracellular K⁺ dynamics. Increased neuronal firing leads to elevated K⁺ extrusion, requiring astrocyte-mediated buffering via connexin channels. Connexin expression and activity are themselves regulated by neuronal activity, forming a feedback system that maintains homeostasis under physiological conditions.^61,62,87^ However, our data indicate that this adaptive mechanism becomes maladaptive in the context of the C9-HRE. Elevation of extracellular K⁺, modelling sustained neuronal activity, induced a modest increase in connexin-mediated currents in control astrocytes, but elicited a markedly enhanced response in C9-HRE astrocytes, including those derived from the unaffected twin and father. This suggests that genetically predisposed astrocytes are primed for an abnormal, activity-dependent upregulation of connexin channels.

These findings provide a mechanistic link between two central features of ALS pathophysiology: motor neuron hyperexcitability and astrocyte-mediated toxicity. Increased neuronal activity and consequent K⁺ release may drive pathological Cx43 hemichannel activation in astrocytes, leading to enhanced release of neurotoxic factors and amplification of disease processes. In this context, sustained high levels of physical activity may accelerate this transition by chronically increasing neuronal demand, thereby exacerbating connexin dysfunction in susceptible individuals. Interestingly, a combination of carbenoxolone and riluzole led to a significant decrease in the toxicity of C9-astrocytes towards co-cultured motor neurons.

These data support a model in which strenuous physical activity acts not as a primary cause of ALS, but as a modifier that interacts with genetic susceptibility to promote astrocyte dysfunction. This activity-dependent mechanism may represent a key pathway linking environmental exposure to disease phenoconversion, highlighting astrocyte connexin signalling as a convergence point between neuronal activity and neurodegeneration.

Leveraging the genetic homogeneity of the iAstrocytes generated from three genetically identical biopsies, we performed translatome RNA-sequencing and single-subject pathways analysis. The affected twin early showed consistent downregulation of GO terms which, combined with targeted analysis showing a reduction in much of the translation initiation machinery, suggests that these early symptomatic astrocytes exhibit translation repression. PERK-mediated eIF2α phosphorylation has been observed to induce reactive astrocytes, displaying altered secretome and reduced synaptic functionality, ultimately leading to increased neuronal toxicity.^88^ In mice, mutations of eIF2α have been associated with vanishing white matter disease, with alterations in the astrocytic translatome identified as a key disease mechanism.^89^ Another key regulator of translation initiation, eIF4E, was also reduced in affected twin early astrocytes. Interestingly, this protein is phosphorylated by mammalian target of rapamycin (mTOR) signalling which is enhanced by exercise but identified as a system that becomes aberrant following extreme exercise, potentially contributing to early onset development of ALS.^58,90^

Within the context of C9-ALS, repression of canonical translation via eIF2α phosphorylation can lead to enhanced non-canonical RAN translation, increasing DPR production. Our data support this, whereby the unaffected twin has moderate DPR expression, substantially increased in the early symptomatic astrocytes, before reducing to the healthy control baseline in the late symptomatic astrocytes, with disease-stage expression correlating with the suspected alterations in canonical translation.^91^ A recent study has highlighted eIF1A and eIF5B as potential repressors of RAN translation and these proteins merit further investigation as neuroprotective targets.^92^

Additionally, GO terms uniquely upregulated in unaffected twin compared with affected twin early were used to identify potential novel neuroprotective targets. The top terms related to protein homeostasis, suggesting an impairment in the early symptomatic astrocytes, which may contribute to the observed accumulation of nuclear RNA foci. Interestingly, the unaffected twin astrocytes had the highest proportion of cells containing TDP-43 inclusions, while being only mildly toxic, suggesting that these asymptomatic cells are effectively dealing with this pathological burden.

In conclusion, astrocytes directly converted from the fibroblasts of a C9-ALS pedigree covering multiple disease stages display phenotypes representative of these differential disease states. This reinforces the power of this model to maintain ageing features through retention of epigenetic alterations. Through this, we observed pathological hallmarks in asymptomatic astrocytes, which then accumulated in the early symptomatic phase, before reducing at the late disease stage. Interestingly, this accumulation negatively correlated with non-cell autonomous toxicity to motor neurons, suggesting that these features exist as early events contributing to disease development. A worsening connexin-mediated electrophysiological impairment did correlate with toxicity and further experimentation assessing the secretome of these astrocytes may elucidate additional toxic factors in the late-stage symptomatic astrocytes, beyond DPR and TDP-43 pathology. It is important to note the unaffected twin remains healthy 12 years after biopsy collection, though is still younger than the average age of onset of ALS and may remain susceptible to disease in later life.

## Data availability

Materials used in this study can be made available subject to Material Transfer agreements (MTAs). RNA Sequencing data will be made available on an appropriate public database following publication of the manuscript. All other data are available in the manuscript main text and supplementary materials.

## Acknowledgements

The authors thank the ALS patients and healthy control subjects who kindly contributed time and biosample donations to support this project.

Kind appreciation to Yves A. Lussier for sharing bioinformatic scripts for N-of-1-pathways analysis, and to Sam Haldenby for preparing the translatomic sequencing data for bioinformatic analysis.

Many thanks also to Rachel Patel and Helen Hipperson for assistance with fibroblast phenol-chloroform DNA extraction. A final acknowledgement to Allison P. Berg, Amrutha Pattamatta, Joanne Berghout, Rob Moccia, and Christine Bulawa for contributions to the pilot study prior to this project.

## Author contributions

**Conceptualization:** PJS, JC-K, MRL

**Methodology**: ACS, ISP, CM, MP, MW, CDSS, AH, LC, JK, GMH, LF, MRL, JK, PJS

**Investigation**: ACS, ISP, CM, GEK, RS, MP, MW, CDSS, MRL, JK, PJS

**Visualization**: ACS, CM, MP, CDSS, MRL

**Funding acquisition**: PJS, JC-K, JK

**Project administration**: PJS, JC-K, LF, MRL, JK

**Supervision**: PJS, JC-K, AH, LC

**Writing** – original draft: ACS, MRL, JC-K, PJS

**Writing** – manuscript review & editing: All authors

## References

1. Nógrádi B, Nógrádi-Halmi D, Erdélyi-Furka B, Kádár Z, Csont T, Gáspár R. Mechanism of motoneuronal and pyramidal cell death in amyotrophic lateral sclerosis and its potential therapeutic modulation. Cell Death Discovery. 2024;10:291.

2. Schweingruber C, Hedlund E. The Cell Autonomous and Non-Cell Autonomous Aspects of Neuronal Vulnerability and Resilience in Amyotrophic Lateral Sclerosis. Biology. 2022;11(8):1191.

3. DeJesus-Hernandez M, Mackenzie IR, Boeve BF, et al. Expanded GGGGCC hexanucleotide repeat in noncoding region of C9ORF72 causes chromosome 9p-linked FTD and ALS. Neuron. 2011;72:245–256.

4. Renton AE, Majounie E, Waite A, et al. A hexanucleotide repeat expansion in C9ORF72 is the cause of chromosome 9p21-linked ALS-FTD. Neuron. 2011;72:257–268.

5. Majounie E, Renton AE, Mok K, et al. Frequency of the C9orf72 hexanucleotide repeat expansion in patients with amyotrophic lateral sclerosis and frontotemporal dementia: a cross-sectional study. Lancet Neurol. 2012;11(4):323–30.

6. Murphy NA, Arthur KC, Tienari PJ, et al. Age-related penetrance of the *C9orf72* repeat expansion. Sci Rep. 2017;7:2116.

7. Ryan M, Doherty MA, Al Khleifat A, et al. *C9orf72* Repeat Expansion Discordance in 6 Multigenerational Kindreds. Neurol Genet. 2023;10(1):e200112.

8. Van Langenhove T, Piguet O, Burrell JR, et al. Predicting Development of Amyotrophic Lateral Sclerosis in Frontotemporal Dementia. J Alzheimers Dis. 2017;58(1):163–170.

9. Fratta P, Poulter M, Lashley T, et al. Homozygosity for the C9orf72 GGGGCC repeat expansion in frontotemporal dementia. Acta Neuropathol. 2013;126(3):401–9.

10. Cooper-Knock J, Higginbottom A, Connor-Robson N, et al. C9ORF72 transcription in a frontotemporal dementia case with two expanded alleles. Neurology. 2013;81(19):1719–21.

11. Belzil VV, Bauer PO, Prudencio M, et al. Reduced C9orf72 gene expression in c9FTD/ALS is caused by histone trimethylation, an epigenetic event detectable in blood. Acta Neuropathol. 2013;126(6):895–905.

12. Zampatti S, Peconi C, Campopiano R, Gambardella S, Caltagirone C and Giardina E. C9orf72-Related Neurodegenerative Diseases: From Clinical Diagnosis to Therapeutic Strategies. Front. Aging Neurosci. 2022;14:907122.

13. Balendra R, Isaacs AM. C9orf72-mediated ALS and FTD: multiple pathways to disease. Nat Rev Neurol. 2018;14(9):544–558.

14. Benatar M, Wuu J, McHutchison C, et al. Preventing amyotrophic lateral sclerosis: insights from pre-symptomatic neurodegenerative diseases. Brain. 2022;145(1):27–44.

15. Maugeri G, D’Agata V, Magrì B, et al. Neuroprotective Effects of Physical Activity via the Adaptation of Astrocytes. Cells. 2021;10(6):1542.

16. Won W, Bhalla M, Lee J, Lee CJ. Astrocytes as Key Regulators of Neural Signaling in Health and Disease. Annu Rev Neurosci. 2025;48(1):251–276.

17. Augusto-Oliveira M, Arrifano G, Leal-Nazaré CG, et al. Lifestyle Drives Astroglial Plasticity Toward Cognitive Improvement: Roles of Physical Exercise, Environmental Enrichment, Diet, and Sleep. Molecular Neurobiology. 2026;63:42.

18. Haidet-Phillips AM, Hester ME, Miranda CJ, et al. Astrocytes from familial and sporadic ALS patients are toxic to motor neurons. Nat Biotechnol. 2011;29(9):824–8.

19. Meyer K, Ferraiuolo L, Miranda CJ, et al. Direct conversion of patient fibroblasts demonstrates non-cell autonomous toxicity of astrocytes to motor neurons in familial and sporadic ALS. Proc Natl Acad Sci USA. 2014;111(2):829–32.

20. Ferraiuolo L, Meyer K. Astrocyte Toxicity In Motor Neuron Disease: Progress and Future Hopes. Future Neurology. 2014;9:149–161.

21. Varcianna A, Myszczynska MA, Castelli LM, et al. Micro-RNAs secreted through astrocyte-derived extracellular vesicles cause neuronal network degeneration in C9orf72 ALS. eBioMedicine. 2019;40:626–635.

22. Allen SP, Hall B, Castelli LM, et al. Astrocyte adenosine deaminase loss increases motor neuron toxicity in amyotrophic lateral sclerosis. Brain. 2019;142(3):586–605.

23. Birger A, Ben-Dor I, Ottolenghi M, et al. Human iPSC-derived astrocytes from ALS patients with mutated C9ORF72 show increased oxidative stress and neurotoxicity. eBioMedicine. 2019;50: 274–289.

24. Zhao C, Devlin AC, Chouhan AK, et al. Mutant C9orf72 human iPSC-derived astrocytes cause non-cell autonomous motor neuron pathophysiology. Glia. 2020;68(5):1046–1064.

25. Arredondo C, Cefaliello C, Dyrda A, et al. Excessive release of inorganic polyphosphate by ALS/FTD astrocytes causes non-cell-autonomous toxicity to motoneurons. Neuron. 2022;110(10):1656–1670.e12.

26. Stoklund Dittlau K, Terrie L, Baatsen P, et al. FUS-ALS hiPSC-derived astrocytes impair human motor units through both gain-of-toxicity and loss-of-support mechanisms. Molecular Neurodegeneration. 2023;18(1):5.

27. Stoklund Dittlau K, Van Den Bosch L. Why should we care about astrocytes in a motor neuron disease?. Front. Mol. Med. 2023;3:1047540.

28. Provenzano F, Torazza C, Bonifacino T, Bonanno G, Milanese M. The Key Role of Astrocytes in Amyotrophic Lateral Sclerosis and Their Commitment to Glutamate Excitotoxicity. Int J Mol Sci. 2023;24(20):15430.

29. Chiò A, Mora G. Untangling the knot: Lifetime physical exercise and amyotrophic lateral sclerosis. EBioMedicine. 2021;69:103438.

30. Chiò A, Benzi G, Dossena M, et al. Severely increased risk of amyotrophic lateral sclerosis among Italian professional football players. Brain. 2005;128:472–476.

31. Lehman EJ, Hein MJ, Baron SL, Gersic CM. Neurodegenerative causes of death among retired National Football League players. Neurology. 2012;79:1970–1974.

32. Fang F, Hållmarker U, James S, et al. Amyotrophic lateral sclerosis among cross-country skiers in Sweden. Eur J Epidemiol. 2016;31(3):247–53.

33. Julian TH, Glascow N, Fisher Barry AD, et al. Physical exercise is a risk factor for amyotrophic lateral sclerosis: Convergent evidence from Mendelian randomisation, transcriptomics and risk genotypes. EBioMedicine. 2021;68:103397.

34. Chapman L, Cooper-Knock J, Shaw PJ. Physical activity as an exogenous risk factor for amyotrophic lateral sclerosis: a review of the evidence. Brain. 2023;146(5):1745–1757.

35. Visser AE, Rooney JPK, D’Ovidio F, et al. Multicentre, cross-cultural, population-based, case-control study of physical activity as risk factor for amyotrophic lateral sclerosis. J Neurol Neurosurg Psychiatry. 2018;89:797–803.

36. Vaage A, Meyer HE, Landgraff IK, Myrstad M, Holmøy T, Nakken O. Physical Activity, Fitness, and Long-Term Risk of Amyotrophic Lateral Sclerosis. Neurology. 2024;103(2):e209575.

37. Li S, Zheng F, Liu S, et al. Physical exercise protects neurons from energy deficit and improves cognitive function by upregulating astrocytic Slc2a1 in Alzheimer’s disease. Experimental Neurology. 2026;395:115493.

38. Fabricio de Souza R, Lopes Augusto R, Arruda de Moraes SR, et al. Ultra-Endurance Associated With Moderate Exercise in Rats Induces Cerebellar Oxidative Stress and Impairs Reactive GFAP Isoform Profile. Front. Mol. Neurosci. 2020;13:157.

39. Li Y, Luo Y, Tang J, et al. The positive effects of running exercise on hippocampal astrocytes in a rat model of depression. Transl Psychiatry 2021;11:83.

40. Gatto N, Dos Santos Souza C, Shaw AC, et al. Directly converted astrocytes retain the ageing features of the donor fibroblasts and elucidate the astrocytic contribution to human CNS health and disease. Aging Cell. 2021;20(1):e13281.

41. Castillo-Fernandez J, Spector T, Bell J. Epigenetics of discordant monozygotic twins: implications for disease. Genome Med. 2014;6(7):60.

42. Ingelfinger F, Gerdes L, Kavaka V, et al. Twin study reveals non-heritable immune perturbations in multiple sclerosis. Nature. 2022;603:152–158.

43. Xi Z, van Blitterswijk M, Zhang M, et al. Jump from Pre-mutation to Pathologic Expansion in C9orf72. The American Journal of Human Genetics. 2015;96:962–970.

44. Van Mossevelde S, van der Zee J, Gijselinck I, et al. Clinical Evidence of Disease Anticipation in Families Segregating a C9orf72 Repeat Expansion. JAMA Neurol. 2017;74(4):445–452.

45. Liu EY, Russ J, Wu K, et al. C9orf72 hypermethylation protects against repeat expansion-associated pathology in ALS/FTD. Acta Neuropathol. 2014;128(4):525–41.

46. Shepheard SR, Parker MD, Cooper-Knock J, et al. Value of systematic genetic screening of patients with amyotrophic lateral sclerosis. J Neurol Neurosurg Psychiatry. 2021;92(5):510–518.

47. Iacoangeli A, Al Khleifat A, Sproviero W, et al. ALSgeneScanner: a pipeline for the analysis and interpretation of DNA sequencing data of ALS patients. Amyotroph Lateral Scler Frontotemporal Degener. 2019;20(3-4):207–215.

48. van der Spek RAA, van Rheenen W, Pulit SL, Kenna KP, van den Berg LH, Veldink JH; Project MinE ALS Sequencing Consortium. The project MinE databrowser: bringing large-scale whole-genome sequencing in ALS to researchers and the public. Amyotroph Lateral Scler Frontotemporal Degener. 2019;20(5-6):432–440.

49. Neuenschwander AG, Thai KK, Figueroa KP, Pulst SM. Amyotrophic Lateral Sclerosis Risk for Spinocerebellar Ataxia Type 2 *ATXN2* CAG Repeat Alleles: A Meta-analysis. JAMA Neurol. 2014;71(12):1529–1534.

50. Hop PJ, Kooyman M, Kenna BJ, et al. Large-scale exome analyses reveal new rare variant contributions in amyotrophic lateral sclerosis. Nat Genet. 2026;58:717–725.

51. Verkhratsky A, Parpura V, Vardjan N, Zorec R. Physiology of Astroglia. Adv Exp Med Biol. 2019;1175:45–91.

52. Zheng J, Xie Y, Ren L, et al. GLP-1 improves the supportive ability of astrocytes to neurons by promoting aerobic glycolysis in Alzheimer’s disease. Mol Metab. 2021;47:101180.

53. Almad A, Doreswamy A, Gross S, et al. Connexin 43 in astrocytes contributes to motor neuron toxicity in amyotrophic lateral sclerosis. Glia. 2016;64(7):1154–1169.

54. Almad A, Taga A, Joseph J, et al. Cx43 hemichannels contribute to astrocyte-mediated toxicity in sporadic and familial ALS. Proc Natl Acad Sci USA. 2022;119(13);e2107391119.

55. Abudara V, Bechberger J, Freitas-Andrade M, et al. The connexin43 mimetic peptide Gap19 inhibits hemichannels without altering gap junctional communication in astrocytes. Front Cell Neurosci. 2014;8:306.

56. Lissoni A, Wang N, Nezlobinskii T, et al. Gap19, a Cx43 Hemichannel Inhibitor, Acts as a Gating Modifier That Decreases Main State Opening While Increasing Substate Gating. Int J Mol Sci. 2020;21(19):7340.

57. Hansen DB, Braunstein TH, Nielsen MS, MacAulay N. Distinct permeation profiles of the connexin 30 and 43 hemichannels. FEBS Letters. 2014;588(8):1446–1457.

58. O’Brien D, Alhathli E, Harwood C, et al. Extreme exercise in males is linked to mTOR signalling and onset of amyotrophic lateral sclerosis, Brain. 2025;148(10):3652–3664.

59. Allen SP, Hall B, Woof R, et al. *C9orf72* expansion within astrocytes reduces metabolic flexibility in amyotrophic lateral sclerosis, Brain, 2019;142,12:3771–3790.

60. Li F, Geng X, Yun HJ, Haddad Y, Chen Y, Ding Y. Neuroplastic Effect of Exercise Through Astrocytes Activation and Cellular Crosstalk. Aging Dis. 2021;12(7):1644–1657.

61. Rouach N, Glowinski J, Giaume C. Activity-dependent neuronal control of gap-junctional communication in astrocytes. J Cell Biol. 2000;149(7):1513–26.

62. Giaume C, Naus CC, Sáez JC, Leybaert L. Glial Connexins and Pannexins in the Healthy and Diseased Brain. Physiol Rev. 2021;101(1):93–145.

63. Lin Y, Dodd J, Cutillo L, et al. GRASPS: a simple-to-operate translatome technology reveals omics-hidden disease-associated pathways in TDP-43-related amyotrophic lateral sclerosis. bioRxiv. [Preprint] doi:10.1101/2024.03.04.583294.

64. Gardeux V, Achour I, Li J, et al. ’N-of-1-pathways’ unveils personal deregulated mechanisms from a single pair of RNA-Seq samples: towards precision medicine. J Am Med Inform Assoc. 2014;21(6):1015–1025.

65. Schissler AG, Gardeux V, Li Q, et al. Dynamic changes of RNA-sequencing expression for precision medicine: N-of-1-pathways Mahalanobis distance within pathways of single subjects predicts breast cancer survival. Bioinformatics. 2015;31(12):i293–302.

66. Donnelly N, Gorman AM, Gupta S, Samali A. The eIF2α kinases: their structures and functions. Cell Mol Life Sci. 2013;70(19):3493–3511.

67. Le Dréau G, Kular L, Nicot AB, et al. NOV/CCN3 upregulates CCL2 and CXCL1 expression in astrocytes through beta1 and beta5 integrins. Glia. 2010;58(12):1510–21.

68. Fu CT, Bechberger JF, Ozog MA, Perbal B, Naus CC. CCN3 (NOV) interacts with connexin43 in C6 glioma cells: possible mechanism of connexin-mediated growth suppression. J Biol Chem. 2004;279(35):36943–36950.

69. Tazekaar GHP, Hop PJ, Seelen M, et al. Whole genome sequencing analysis reveals post-zygotic mutation variability in monozygotic twins discordant for amyotrophic lateral sclerosis. Neurobiology of Aging. 2023;122:76–87.

70. Tarr IS, McCann EP, Benyamin B, et al. Monozygotic twins and triplets discordant for amyotrophic lateral sclerosis display differential methylation and gene expression. Scientific Reports. 2019;9:8254.

71. Seminary ER, Santarriaga S, Wheeler L, et al. Motor Neuron Generation from iPSCs from Identical Twins Discordant for Amyotrophic Lateral Sclerosis. Cells. 2020;9(3):571.

72. Papp B, Plath K. Epigenetics of reprogramming to induced pluripotency. Cell. 2013;152(6):1324–43.

73. Gijselinck I, Van Mossevelde S, van der Zee J, et al. The *C9orf72* repeat size correlates with onset age of disease, DNA methylation and transcriptional downregulation of the promoter. Molecular Psychiatry. 2016;21:1112–1124.

74. Esselin F, Mouzat K, Polge A, et al. Clinical Phenotype and Inheritance in Patients With C9ORF72 Hexanucleotide Repeat Expansion: Results From a Large French Cohort. Front Neurosci. 2020;28(14):316.

75. Cooper-Knock J, Higginbottom A, Stopford MJ, et al. Antisense RNA foci in the motor neurons of C9ORF72-ALS patients are associated with TDP-43 proteinopathy. Acta Neuropathol. 2015;130(1):63–75.

76. Vahsen BF, Nalluru S, Morgan GR, et al. C9orf72-ALS human iPSC microglia are pro-inflammatory and toxic to co-cultured motor neurons via MMP9. Nature Communications. 2023;14:5898.

77. McEachin ZT, Chung M, Stratton SA, et al. Molecular impact of antisense oligonucleotide therapy in *C9orf72*-associated ALS. Cell. 2025;188(23):6424–6435.

78. Zhang M, Zhengrui X, Mahdi G, et al. Genetic and epigenetic study of ALS-discordant identical twins with double mutations in *SOD1* and *ARHGEF28*. Journal of Neurology, Neurosurgery & Psychiatry. 2016;87:1268–1270.

79. Young P, Kum Jew S, Buckland ME, Pamphlett R, Suter CM. Epigenetic differences between monozygotic twins discordant for amyotrophic lateral sclerosis (ALS) provide clues to disease pathogenesis. PLOS One. 2017;12(8):e0182638.

80. Orellana JA, Froger N, Ezan P, et al. ATP and glutamate released via astroglial connexin 43 hemichannels mediate neuronal death through activation of pannexin 1 hemichannels. J Neurochem. 2011;118(5):826–840.

81. Takeuchi H, Jin S, Wang J, et al. Tumor Necrosis Factor-α Induces Neurotoxicity via Glutamate Release from Hemichannels of Activated Microglia in an Autocrine Manner. Journal of Biological Chemistry, 2006;281:21362–21368.

82. Van Den Bosch L, Vandenberghe W, Klaassen H, Van Houtte E, Robberecht W. Ca^2+^-permeable AMPA receptors and selective vulnerability of motor neurons. Journal of the Neurological Sciences. 2000;180(1-2):29–34.

83. Van Den Bosch L, Van Damme P, Bogaert E, Robberecht W. The role of excitotoxicity in the pathogenesis of amyotrophic lateral sclerosis. Biochim Biophys Acta. 2006;1762(11-12):1068–1082.

84. Figiel M, Allritz C, Lehmann C, Engele J. Gap junctional control of glial glutamate transporter expression. Molecular and Cellular Neuroscience. 2007;35(1):130–137.

85. Zhang J, Riquelme M, Hua R, Acosta F, Gu S, Jiang J. Connexin 43 hemichannels regulate mitochondrial ATP generation, mobilization, and mitochondrial homeostasis against oxidative stress. Elife. 2022;11:e82206.

86. Vandoorne T, De Bock K, Van Den Bosch L. Energy metabolism in ALS: an underappreciated opportunity? Acta Neuropathol. 2018;135(4):489–509.

87. Charvériat M, Mouthon F, Rein W, Verkhratsky A. Connexins as therapeutic targets in neurological and neuropsychiatric disorders. Biochim Biophys Acta. 2021;1867(5):166098.

88. Smith HL, Freeman OJ, Butcher AJ, et al. Astrocyte Unfolded Protein Response Induces a Specific Reactivity State that Causes Non-Cell-Autonomous Neuronal Degeneration. Neuron. 2020;105(5):855–866.

89. Mandelboum S, Lev-Ari L, Atzmon A, et al. The translational landscape of reactive astrocytes reveals the impact of eIF2B-mediated dysregulation in VWM disease. NAR Molecular Medicine. 2025;2(2):ugaf009.

90. Maracci C, Motta S, Romagnoli A, Constantino M, Perego P, Di Marino D. The mTOR/4E-BP1/eIF4E Signalling Pathway as a Source of Cancer Drug Targets. Curr Med Chem. 2022;29(20):3501–3529.

91. Green KM, Glineburg MR, Kearse MG, et al. RAN translation at C9orf72-associated repeat expansions is selectively enhanced by the integrated stress response. Nat Commun. 2017;8(8):2005.

92. Ito H, Machida K, Fujino Y, et al. Canonical translation factors eIF1A and eIF5B modulate the initiation step of repeat-associated non-AUG translation. Nucleic Acids Research. 2025;53(18):gkaf994.

