## Supplementary Material for "Astrocyte dysfunction distinguishes monozygotic twin *C9orf72* expansion carriers discordant for amyotrophic lateral sclerosis"

#### Supplementary Methodology

##### Skin biopsy collection and fibroblast banking

The biopsy was cut into several small pieces and placed into a T25 flask, which was pre-coated with FBS. This was fed with 0.5 ml (50:50) mix of FBS and EMEM, supplemented with 10% FBS, 2 mM glutamine, 1% uridine, 1% MEM vitamins, 1% MEM non-essential amino acids, 1% P/S and 1 mM sodium pyruvate in humid incubators at 37°C with 5% CO2. On day two, 50% of the media was removed and replaced with fresh solution. This feeding continued for the first week, gradually increasing the volume of EMEM up to

6 ml total as cells began to grow out of the skin plugs. When cells became confluent around the biopsies (19-23 days), the flask was split into a T75 flask using trypsin and with the media saved for mycoplasma testing. A further split occurred 5-7 days after and fibroblasts were frozen after this point in a freeze mix containing 80% EMEM media, 10% FBS and 10% dimethylsulfoxide (DMSO; Sigma) and stored in liquid nitrogen.

#### Direct conversion of skin fibroblasts into iNPCs (induced Neural Progenitor Cells)

Fibroblasts between passage 5-10 were grown in fibroblast medium consisting of DMEM (Gibco Life Tech) with 10% FBS and 1% P/S. Fibroblasts were dissociated using trypsin and plated in a six-well plate coated with 2.5 µg/mL human fibronectin (Merck) at the following cell densities: 80,000, 120,000, and 160,000. After incubating at 37°C overnight, plates were checked for optimal density of 70% with evenly spread cells. The optimal well was transduced in the late afternoon of this day, using a cocktail of retroviral vectors expressing OCT3/4, SOX2, c-MYC, and KLF4 (ALSTEM). The following morning, the vectors were removed and fresh fibroblast medium added. The next day, the medium was switched to iNPC conversion medium, consisting of DMEM/F12 (Gibco), 1% N2-supplement (Gibco), 1% B27-supplement (Gibco), 40 ng/mL epidermal growth factor (EGF; Peprotech) and 20 ng/mL fibroblast growth factor (FGF; Peprotech). The morphology changed after a further 2-4 days, with cells becoming spherical and forming rosettes. 6-7 days after induction, the cells were lifted with Accutase (StemPro® Accutase® Cell Dissociation Reagent; Gibco) and re-plated into two wells of a six-well plate coated with 10 µg/mL fibronectin. Culturing continued over the course of 18 days until cells were in a 10 $cm^2$ dish coated with 5 µg/mL fibronectin. After the culture reached this stage, the medium was switched to iNPC medium (DMEM/F12, 1% N2, 1% B27) supplemented with 40 ng/mL FGF, with EGF removed.

#### Fibroblast DNA Extraction

Fibroblasts were cultured as before, with pellets generated from 3x confluent T75 flasks to ensure sufficient starting material. Pellets were then processed using a modified phenol chloroform extraction protocol to extract high molecular weight genomic DNA (HMW gDNA).[1] Pre-cut pipette tips and gentle mixing with no vortexing was used throughout the protocol to prevent DNA shearing. First, SET buffer was prepared (0.15 M NaCl, 0.05 M Tris base (Tris-(hydroxymethyl) aminomethane), 0.001 M EDTA, pH 8.0) and the pellet thawed in 500 µL of this solution. 10 µL RNAse (100 mg/mL) was added and incubated for 2 minutes at room temperature. 13 µL of 20% SDS was added and incubated for 30 minutes at 37°C. 7.5 µL proteinase K (20 mg/mL) was added, the samples mixed well and spun down, before incubating overnight in a 55°C water bath (with an additional mix after the first hour).

The next day, the samples were spun down, 50 µL 5 M NaCl added, mixed well, and spun again. An equal volume of phenol was added to the sample (500-600 µL) and mixed well to create a homogeneous mix.

The samples were left for 60 minutes in the fume hood with regular mixing, before spinning for 15 minutes at 10,000 rpm. In a new set of labelled tubes, 250 µL phenol and 250 µL chloroform/isoamyl alcohol (24:1) were added. With a pre-cut 1,000 µL tip, the DNA containing supernatant of the samples was transferred into the new tubes, mixed, and spun for 15 minutes at 10,000 rpm. Another set of tubes containing phenol and chloroform/isoamyl alcohol were prepared, the supernatant transferred, mixed, and spun for 15 minutes at 10,000 rpm.

In a further set of fresh tubes, 500 µL chloroform/isoamyl alcohol (24:1) was added, the sample supernatant transferred, mixed, and spun for 15 minutes at 10,000 rpm. 50 µL of NaAc (3 M) was added to a new set of tubes and the sample supernatant transferred. 1,000 µL of ice cold 95% EtOH was added, and the samples mixed well until the DNA precipitated to create 'fluff'. The DNA fluff was retrieved using a glass rod by carefully spinning the rod around the fluff, this was then rinsed three times in ice cold 70% EtOH (three separate tubes) and left to dry in the air until the EtOH evaporated. Finally, the DNA was dissolved in 50-200 µL (adjusted to the amount of DNA) 0.1x TE (pH 7.8) with the rod left in the tubes until clean and further dissolved at 4°C overnight. The DNA purity was assessed using a Nanodrop spectrophotometer (Fisher Scientific, Hampton, NH, USA) and quantified using a Qubit fluorometer using the double stranded DNA Broad Range Assay Kit (Fisher Scientific, Hampton, NH, USA), diluted to desired concentration and run on a 0.8% agarose gel (50 V for 90 minutes) for pulse-field size determination (ideally 120-150 kb), with lambda DNA as the control marker. The samples were stored at -80°C until needed.

#### Whole Genome Sequencing (WGS)

Fibroblast HMW gDNA samples were shipped on dry ice to Fulgent Therapeutics. Libraries were prepared using the TruSeq Nano DNA Protocol (Illumina) and sequenced on a HiSeq X system using paired-end 150 bp reads (Illumina). Raw sequence reads were aligned to Human Reference Genome GRCh38 and variants identified using the HaplotypeCaller package in the Genome Analysis Tooklit.[2] Variants were filtered and annotated using the Functional Annotator package, with additional information provided by the Ensembl Variant Effect Predictor and the LOFTEE package.[3,4]

#### *C9orf72*-repeat sizing using Southern blotting

C9-HRE length in *C9orf72* fibroblasts was determined by Southern blotting using a previously described protocol.[5] Firstly, a probe was generated specific for the flanking regions of the *C9orf72* hexanucleotide repeat (**Supplementary Table 2**). A reaction mix was generated containing PCR grade $H_2O$, Phire II buffer (Fisher), Betaine (Sigma), dNTPs (Fisher), the forward and reverse C9orf72 probe, Phire Taq II (Fisher), 7-deaza-dGTP (Roche), and fibroblast HMW gDNA.

For C9-HRE assessment in *C9orf72* iNPCs, gDNA was extracted using DNeasy Blood & Tissue Kit (Qiagen) according to the manufacturer's instructions, with C9-HRE length determined using a previously published Southern blotting protocol.[6]

#### *C9orf72*-repeat sizing using Nanopore sequencing

Fibroblast HMW gDNA samples with native DNA fragment lengths of 120 kb were fragmented by passing the DNA through a 25-gauge needle 3 times to achieve optimal fragments lengths of 20-30 kb. 5 µg of DNA in a volume of 53.5 µL was combined with 6.5 µL of FFPE DNA Repair Buffer and 2.0 µL of FFPE DNA repair Mix (New England Biolabs). The reaction was incubated at 20°C for 30 minutes. A 1.8x Agencourt AMPure XP (Beckman Coulter) bead clean-up was performed to allow buffer exchange prior to target enrichment and sequencing. The repaired DNA was combined with 111.6 µL of beads and incubated at room temperature for 15 minutes. Beads were captured on a magnet and the supernatant discarded. The bead pellet was washed two times with 80% ethanol then allowed to dry for 1 minute at room temperature. The beads were re-suspended in 30 µL of nuclease free water (heated to 37°C) and incubated for 15 minutes. Beads were then captured and the supernatant retained in a fresh tube.

DNA recovery was assessed using a Nanodrop Spectrophotometer and DNA was quantified using a Qubit double stranded DNA Broad Range Assay Kit. crRNAs (Integrated DNA Technologies) were pooled in equimolar amounts at a concentration of 100 µM (**Supplementary Table 3**). A crRNA-tracrRNA duplex was formed by combining 1 µL of the 100 µM crRNA pool, 1 µl of 100 µM Alt-R® CRISPR-Cas9 tracrRNA and 8 µl of duplex buffer (both Integrated DNA Technologies) and incubating at 95°C for 5 minutes. Ribonucleoprotein complexes (RNPs) were created by combining 10 µl of 10x CutSmart® buffer (New England Biolabs), 79.2 µl nuclease-free water, 10 µl of the annealed crRNA-tracrRNA duplex (10 µM) and 0.8 µl of Alt-R® S.p. HiFi Cas9 V3 (62 µM) (Integrated DNA Technologies), which was incubated at room temperature for 30 minutes.

Genomic DNA was dephosphorylated by combining 3 µl of 10x CutSmart® buffer, 23 µl of DNA and 4 µl Quick CIP (New England Biolabs) followed by incubation at 37°C for 10 min then 80°C for 2 minutes to inactivate the enzyme. Cleavage and dA-tailing of DNA was performed simultaneously by combining the 30 µL of dephosphorylated genomic DNA with 10 µL of Cas9 RNPs, 1 µL of 10 mM dATP and 1 µL of Taq polymerase (both New England Biolabs) and incubating at 37°C for 15 minutes followed by 72°C to inactivate the Cas9 nuclease.

Nanopore sequencing adapters were then ligated by combining 20 µL LNB (Oxford Nanopore Technologies), 10 µL T4 DNA Ligase (New England Biolabs), 3 µL nuclease-free water and 5 µL AMX (Oxford Nanopore Technologies), followed by incubation for 10 minutes at room temperature. A 0.3x Agencourt AMPure XP bead clean-up was performed to remove excess unligated sequencing adapters.

Short fragments of DNA were also removed by washing the beads twice with 250 µL of LFB (Oxford Nanopore). The final library was eluted by adding 20 µL of elution buffer (pre-warmed to 65˚C) and incubating for 10 minutes followed by bead capture.

A MinION R9.4.1 flow cell (Oxford Nanopore) was primed for sample loading by preparing the priming mix (1170 µL FB and 30 µL FLT, both Oxford Nanopore) and adding 800 µL of this to the priming port and incubating for 5 minutes. Immediately prior to sample loading a further 200 µL of priming mix was added to the priming port, with the SpotON sample port in the open position. A sequencing mix was prepared by combining 20 µL of the eluted library, 30 µL SQB and 20 µL of LB (both Oxford Nanopore). The mixture was then added to the SpotON sample port dropwise to allow the mix to disperse across the sensor array. The SpotON and priming ports were then closed and the sequencing run initiated using MinKNOW software (Oxford Nanopore Technologies).

Reads were mapped to the human genome version 38 (HG38) using minimap2, allowing assessment of *C9orf72* locus coverage.[7] Reads that covered this locus were then further assessed using STRique allowing for estimation of DNA methylation in each called repeat.[8]

#### Targeted Sequencing Panel

The gDNA previously extracted from iNPCs was sent to the Sheffield Diagnostic Genetics Service (SDGS) laboratory for in-depth screening using a panel of 44 ALS and FTD risk genes and a larger pan-neurodegenerative panel of 153 risk genes (**Supplementary Table 4**).[9,10] Samples underwent targeted next-generation sequencing using the SureSelectXT (Design ID: 0836801) automated library preparation, followed by sequencing using an Illumina HiSeq 2500, at a depth of at least 100x.

Additionally, the presence of *C9orf72* repeat expansions was confirmed using fluorescent repeat primed polymerase chain reaction (PCR) with fragment size analysis performed in GeneMapper v3.5 and expansions in excess of 30 repeats reported as potentially pathogenic.[11,12] Finally, Ataxin-2 CAG repeat expansions were investigated using standard PCR with a fluorescently labelled primer, with size analysis via GeneMapper v3.5. CAG expansions of 14-28 repeats were reported as normal, 29-34 associated with ALS, and above this as a risk factor for spinocerebellar ataxia type 2 (SCA2).[13] Discovered variants were returned with information such as the specific mutation (and protein change if applicable) and the number of counts for the variant and the reference allele to assess heterozygosity. Assessment of pathogenicity was performed according to the previously published criteria.

#### Whole-cell patch clamp electrophysiology

iAstrocytes (30,000 cells/well) were plated onto Thermanox 13 mm coverslips placed into 24-well plates for electrophysiological assessment. Coverslips were pre-coated for 5 minutes at room temperature with

2.5 μg/mL fibronectin diluted in PBS. For experiments performed at day 7, iAstrocytes were plated on day 5, and for day 14, iAstrocytes were plated at day 7. iAstrocytes were lifted from the plates using Accutase, which was inactivated by adding PBS. Cells were then centrifuged for 4 minutes at 200 x g. Media changes were performed at day 3 and 10.

The whole-cell patch configuration was used to record macroscopic currents from *in vitro* derived human glial cells as previously described.[14] Reported potential values are corrected for liquid junction potential (+14 mV). Current measurements were typically low-pass filtered online at 2 kHz, digitized at 10 kHz via a BNC-2090A (National Instruments) interface, and recorded to computer using the WinEDR V2 7.6 (J. Dempster, Department of Physiology and Pharmacology, University of Strathclyde, UK). Slope conductance was determined by fitting a linear line to each data point for each cell and taking the slope (dI/dV). Conductance density was determined by taking the slope conductance and dividing this by the measured whole cell capacitance.

#### iAstrocyte Immunocytochemistry (ICC)

10,000 iAstrocytes were plated per well in 96-well plates at day 5 of differentiation and fixed 24 hours after seeding with 4% paraformaldehyde (PFA, Gibco) for 15 minutes at room temperature (RT). Cells were washed 3 times for 5 minutes with phosphate-buffered saline (PBS) and then blocked for 1 hour in a solution containing PBS, 5% horse serum (Gibco) and 0.05% Triton X-100 (Sigma). After blocking, 70 µL of primary antibody, diluted in blocking solution was added per well and incubated overnight at 4°C. The next day, cells were washed once with PBS containing 5% Tween-20 (Sigma) and 2 times in PBS, before incubating with secondary antibody diluted in blocking solution for 1 hour in the dark at RT (**Supplementary Table 5)**. Hoechst (Gibco) diluted 1:6000 was then incubated for 5 minutes in dark conditions at RT as a nuclear counterstain. Finally, cells were washed twice with PBS and images were acquired with the Opera Phenix high-content imager (Perkin Elmer) at x 40 objective magnification (NA 1.0) using the Hoechst 33342, Cy5 (Alexa 647), Alexa 568 and Alexa 488 laser lines.

#### Fluorescent in situ Hybridisation for *C9orf72* RNA Foci (C9-FISH)

The methodology utilised in this study was adapted from an in-house protocol optimised to detect RNA foci in patient tissue, and later, *in vitro* cell models.[15-17] iAstrocytes were plated at a density of 10,000 cells per well in 96-well plates at day 5 of differentiation. Cells were fixed 48 hours after seeding with 4% paraformaldehyde (PFA) in diethyl pyrocarbonate-treated (DEPC, Sigma) PBS (DEPC-PBS) for 15 minutes at RT.

The hybridisation oven (Thermo Fisher Hybrid Oven, Fisher) was turned on and allowed to heat to 68°C, with a foil sheet preventing light-exposure. During this time hybridisation buffer was prepared consisting of 350 µL/mL DEPC-$H_2O$, 0.1 g/mL dextran sulphate (Generon), 50 µL/mL sodium phosphate (Sigma) and 100 µL/mL 20% saline sodium citrate buffer (SSC, containing 3 M NaCl, and 0.3 M sodium citrate at pH 7.0; Sigma) mixed in a covered beaker for 30 minutes. Finally, 500 µL/mL formamide (Sigma) was added after the other reagents had completely mixed. Cells were then permeabilised with 4% DEPC-PFA with 0.1% Triton X-100 (RNAse-Free stock, Sigma) for 15 minutes at RT. For RNAse-treatment control wells, 10 µg/mL RNAse-A (Fisher) in DEPC-PBS was added and incubated for 30 minutes at 37°C, with the remaining wells containing DEPC-PBS. After RNAse-treatment, solutions were removed, 30 µL hybridisation buffer added per well and incubated for 1 hour in the hybridisation oven. Locked Nucleic Acid™ RNA probes (Exiqon) designed to recognise the *C9orf72* Sense (CCCCGGCCCCGGCCCC) and Antisense (GGGGCCGGGGCCGGGG) strands and tagged to TYE563 were heated in Eppendorf tubes on a PCR heat block for at least 75 seconds at 80°C for denaturing. Upon removal from the block, the probes were immediately snap frozen on ice and then diluted 1:400 in hybridisation buffer. 30 µL per well of the Probe-Buffer mix was then added per well and the plate incubated in the oven overnight.

Two washes were then prepared for the second day, the first, containing 2% SSC with 0.1% Triton X-100 (Fisher) and 1:10,000 Hoescht (Gibco) in DEPC-$H_2O$ (wash one) and stored at RT. The second consisting of 0.1% SSC in DEPC-$H_2O$ (wash two) was prepared in duplicate falcon tubes and stored in the oven to heat up.

The next day, the probe was removed and 100 µL per well of wash one was added and incubated for 10 minutes at RT. This wash was removed and 100 µL per well of wash two added and incubated for 10 minutes in the oven at 65°C, this second wash was repeated. The UV Translinker Transilluminator (AnalytikJena) was turned on and a box prepared with packed ice. The final wash was removed, and the plate placed firmly into the ice before placing into the machine. The cross-linker was set to 3000 (x100 $\mu J/cm^2$) and ran for the final sterilisation. The plate was then stored in DEPC-PBS and imaged with the Opera Phenix high-content imager (Perkin Elmer) using the 568 nm laser line (TYE563) and 415 nm laser line (Hoescht) at x40 magnification. Analysis was performed on the Columbus. Thresholds were set based on the $99^{th}$ percentile of RNAse-A treated wells Cy3 expression.

#### Detection of Dipeptide Repeat Proteins using Meso Scale Discovery (MSD) ELISA Immunoassay

Dipeptide repeat proteins (DPRs) were measured using a modified version of the protocol developed and kindly shared by the laboratory of Professor Adrian Isaacs.[18] Lysis buffer was prepared using RIPA Buffer (containing 150 mM NaCl, 1.0% IGEPAL® CA-630, 0.5% sodium deoxycholate, 0.1% sodium dodecyl sulfate (SDS), 50 mM Tris, pH 8.0; Sigma) with 1x cOmplete, Mini, EDTA-free Protease Inhibitor Tablet per 5 mL buffer (Roche) and SDS to a 2% final concentration.

2x 10 cm plates of iAstrocytes were prepared per line and used when confluent at day 7 of differentiation. Media was removed and cells washed in 1x PBS at room temperature. 100-200 µL Lysis Buffer was added per plate and left for 7 minutes, gently swirling to ensure coverage. Using a cell scraper, lysed cells were collected into an Eppendorf tube and kept on ice. Samples were sonicated using the Vibra-Cell Sonicator (Sonics & Materials) with 3x 5 second pulses at 80% power. Samples were then centrifuged at 17,000 x g, room temperature for 20 minutes. Following this, the supernatant was transferred into a new tube and protein content measured using a BCA assay (Bio-Rad). Lysates were diluted to a final concentration of 2 mg/mL at a minimum and stored at -80°C until the ELISA was performed.

To determine concentration of DPRs, an ELISA using electrochemiluminescence (ECL) for detection was performed. Firstly, an MSD plate was coated with 4 µg/mL unconjugated antibody in TBS (ensuring the plate was covered) and shaken at 700 rpm for 2 hours at room temperature. The next day, the standards and samples were prepared by diluting 1:1 in EC Buffer (Meso Scale Diagnostics). The untagged antibody was removed from the plate by flicking, followed by 3x washes in TBS with 0.2% tween (TBST) (washes are fast with immediate removal and blotting to remove excess liquid). After this the plate was blocked with 150 µL/well TBST with 3% milk whilst shaking at 700 rpm for 2 hours at room temperature, followed by a further 3x washes in TBST. Standards and samples added in duplicate with 50 µL max volume, followed by shaking at 700 rpm for a further 2 hours at room temperature. After 3x TBST washes, biotinylated detection antibody (2 µg/mL) prepared and 25 µL added per well, followed by shaking at 700 rpm for 2 hours at room temperature. After a final set of TBST washes, 25 µL MSD Read Buffer T was added per well and protein concentration detected using the MESO QuickPlex SQ 120 (Meso Scale Diagnostics).

#### Murine Hb9-GFP Co-Culture with iAstrocytes

Murine Hb9-GFP$^{+}$ motor neurons were differentiated from mouse embryonic stem cells (mESC; kind gift from Professor Thomas Jessell, Columbia University, New York) containing stable GFP expression controlled by the motor neuron-specific promoter Hb9, were utilised in an astrocyte-motor neuron co-culture model as previously described.[19]

mESC were cultured on primary mouse embryonic fibroblasts (Merck) in mESC medium containing KnockOut DMEM (Gibco), 15% (*v/v*) embryonic stem-cell FBS (Gibco), 2 mM L-glutamine (Gibco), 1% (*v/v*) nonessential amino acids (Gibco), 0.00072% (*v/v*) 2-mercaptoethanol (Sigma). For differentiation mESC were split using trypsin and resuspended in EB medium containing DMEM/F12 (Gibco), 10% (*v/v*) knockout serum replacement (Gibco), 1% N2 (Gibco), 1 mM L-glutamine (Gibco), 0.5% (*w/v*) glucose (Sigma) and 0.0016% (*v/v*) 2-mercaptoethanol (Sigma). Cells were seeded in non-adherent Petri dishes and grew as embryoid bodies, with EB media replenished daily, supplemented with 2 µM retinoic acid (Sigma) and 0.5 µM smoothened agonist (Sigma) from days 2 to 7 to induce MN differentiation. iAstrocytes were plated at day 5 of differentiation at a density of 2,500 cells per well of a 384-well plate (six technical repeats per cell line). The following day, plates were drugged using Echo 550 (Beckman Coulter). On day 7 of differentiation, EBs were dissociated with 200 U/mL papain (Sigma) and plated on top of the iAstrocytes at a density of 10,000 cells per well.

MNs were imaged using a INCELL Analyzer 2000 (GE Healthcare) three days after plating (day 10 of differentiation), with nine fields imaged per well. Subsequent analysis utilised the Columbus™ Data Storage and Analysis System (Perkin Elmer) to identify motor neuron cell bodies and neuronal axons as previously described.[19] A motor neuron was counted as viable if it had at least one intact process attached. MN survival was normalised to the DMSO baseline per biological repeat in each cell line.

#### iPSC-MN Co-Culture with iAstrocytes

Undifferentiated induced pluripotent stem cells (iPSCs) derived from healthy control fibroblasts were maintained in complete mTeSR-Plus Medium (STEMCELL Technologies) in Matrigel® growth factor reduced-coated plates (Corning) according to the manufacturer's recommendations. iPSCs were used between passage 20-28 and cultured in 5% $O_2$ and 5% $CO_2$ at 37°C. Motor neuron differentiation was performed as previously described.[20]

iPSCs were plated into wells of a matrigel-coated six-well plate and grown in basal media supplemented with 3 µM CHIR (Tocris), 2 µM DMH1 (Tocris) and 2 µM SMADi (Tocris) between days 1 to 6. Between days 7 to 12, the cells were fed with basal media supplemented with 1 µM CHIR, 2 µM DMH1, 2 µM SMADi, 0.1 µM RA (STEMCELL Technologies), and 0.5 µM Purmorphamine (PMN, Tocris). On day 12, the resultant NPCs were re-plated onto new matrigel-coated six-well plates. The following day the media was changed to basal media supplemented with 0.5 µM RA and 0.1 µM PMN with daily feeding until day 18. Terminal plating occurred on day 19 with 15,000 cells plated per well of Matrigel-coated 96-well plates in basal media supplemented with 0.5 µM RA, 0.1 µM PMN, 0.1 µM Compound-E (Tocris), 10 ng/mL Brain-derived neurotrophic factor (BDNF; Fisher), 10 ng/mL Ciliary neurotrophic factor (CNTF; Fisher) and 10 ng/mL Insulin-like growth factor 1 (IGF-1; Fisher). From day 28 to 40 the medium was switched to Neurobasal Medium (Gibco) supplemented with 1% B27, 1% P/S, 10 ng/mL BDNF, 10 ng/mL CNTF and 10 ng/mL IGF-1, with feeding every 72 hours.

Mature motor neurons were observed from day 40, after which a co-culture was assembled with 8,000 iAstrocytes per well seeded on top of the mature motor neurons and cultured for 72 hours before fixing. Co-cultured cells were then stained for cleaved caspase-3 (Merck) activation and analysed using the Harmony® Analysis Software (Perkin Elmer). In brief, a mask was used to ensure that only caspase activation in motor neurons was captured and used as a surrogate for toxicity.

#### Exercise quantification using the Historical Adulthood Physical Activity Questionnaire (HAPAQ)

The HAPAQ exercise questionnaire was administered as previously described.[21] From the results of the HAPAQ, durations of various types of physical activity were calculated for each decade of life. Inputting the metabolic equivalent value (MET) for each activity undertaken then enabled conversion into a value of average physical activity energy expenditure (PAEE) in kJ/kg/day. Leisure-time physical exercise was derived by summing the physical activity for all strenuous sport and exercise, and more casual sports and exercise.

#### Single subject RNA-sequencing using Genome-wide RNA Analysis of Stalled Protein Synthesis and N-of-1-pathways analysis

mRNA from iNPC-derived astrocytes undergoing active translation was isolated from ribosomes following fractionation and ultracentrifugation as previously described.[22] All buffers were prepared using DEPC-treated $dH_2O$. Following mRNA isolation, RNA was extracted using Direct-zol RNA miniprep kit according to the manufacturer's instructions. 10-20 µg of total RNA was taken forward on the same day for Poly-A purification using the NEBNext® Poly(A) mRNA Magnetic Isolation Module. The Agilent Bioanalyzer RNA 6000 Pico Chip was used to assess the distribution of mRNA and ensure that no contaminating rRNA remained in the sample.

Poly-A purified samples were shipped to the Centre for Genomic Research at the University of Liverpool for RNA sequencing. Libraries were prepared using the NEBNext Ultra II Directional RNA library preparation kit according to the manufacturer's instructions. The prepared libraries were then sequenced on the Illumina NovaSeq 6000 system, using a single lane of an S4 Flow Cell, generating paired end reads in the 2 x 150 bp configuration. Following completion of sequencing, the raw FASTQ files were trimmed using Cutadapt version 1.2.1, removing reads containing Illumina adapter sequences with a match of 3 bp or more at the 3' end.[23] Additional trimming used Sickle version 1.200 using a minimum window quality score of 20, removing reads shorter than 15 bp. The trimmed reads were then aligned to the human reference genome version GRCh38.p14 using the Salmon and Sleuth packages in R.[24,25] The four samples were normalised with raw transcript counts converted to transcripts per million (TPM) and averaged, with any transcripts below 0.5 TPM removed to normalise for sequencing depth and gene length. 35851 commonly expressed transcripts were retained.

As a final QC measure, MA plots were generated for each of the putative comparisons, the log fold change shown on the y-axis (M) against the average expression level of the two compared samples on the x-axis (A). The values for a given transcript were calculated as: M = log2(Case + 1) - log2(Contrast + 1) and A = (log2(Contrast + 1) + log2(Case + 1)) / 2, with a pseudocount of 1 added to avoid null errors. Differentially expressed (DE) transcripts were selected by an absolute value >1 and the percentage DE calculated (Range: 16.7-22.2%). A correlation between the M and A values was determined and a linear regression slope fitted to each comparison. A maximum $R^2$ value of 0.006 was determined from MA plots, equating to less than 1% of the observed variance and suggesting that the datasets were normally distributed and suitable for further analysis (**Supplementary Table 8**).

N-of-1-*pathways* analysis was computed using the Wilcoxon signed-rank test model.[26,27] A reference list of genes contained in the Gene Ontology knowledgebase was used.[28,29] TPM filtered transcript counts were log transformed as: log(TPM + 10) and a vector generated containing the gene names ordered by row. GO enrichment was computed with the n.of.1.pathways module using the following options: row.genes.names = row.genes.names_Fam, model = "wilcoxon", genes.1 = contrast_TPM, genes.2 = case_TPM, maxSizePathway = 500, minSizePathway = 15, parallelize = TRUE, pathways.set = pathways.set_GO, is.exact = TRUE, seed.value = 123, nb.repeat = 1000, p.dereg.cutoff = 0.05. (Genes.1 and genes.2 altered for each comparison as noted in **Supplementary Table 8**).

The outputs of each enrichment were processed to calculate the difference in ranks (GO_Enrichment$rank_difference <- GO_Enrichment$ranks.pos - GO_Enrichment$ranks.neg), absolute difference (GO_Enrichment$abs_rank_difference <- abs(GO_Enrichment$rank_difference)), and log fold change (GO_Enrichment$log_fold_change <- log2((GO_Enrichment$ranks.neg + 1) / (GO_Enrichment$ranks.pos + 1))). Significant GO terms were determined by a log fold change >1 and P-Value <0.05 (Up-Regulated) or log fold change <1 and p-value <0.05 (Down-Regulated). Volcano plot visualisation was performed using ggplot2. Significant GO terms were retained in a new dataframe and Venn Diagrams generated using ggVennDiagram, with constituent elements extracted for targeted analysis. The generated lists of GO Terms were condensed using the R Shiny application of Reduce + Visualize Gene Ontology (RRVGO) using the following options: Organism – Human, Ontology – Biological Process, Molecular Function, Cellular Component (each separate), Stringency – Medium (0.7), Distance measure – Rel. Each list was condensed with up- and down-regulated GO Terms separated. The reduced terms were filtered by termDispensability with only parent terms retained (value = 0). The condensed lists were annotated with ontology and direction, with the 5 most significant terms for each option (by P-Value) used for visualisation (30 terms max). Finally, constituent genes from GO Terms of interest were extracted to identify dysregulated transcripts for subsequent validation.

### Supplementary Tables

***Supplementary Table 1*** *– Clinical characteristics of donor fibroblasts utilised in this project. (fALS = Familial ALS).*

| **Cell Line** | **Ethnicity** | **Gender** | **Diagnosis** | **Mutation** | **Age at collection (y)** | **RRID** |
|---|---|---|---|---|---|---|
| Healthy control | Caucasian | Male | Non-ALS control | - | 40 | RRID:CVCL_UF81 |
| C9-ALS | Caucasian | Male | fALS | *C9orf72* | 66 | RRID:CVCL_UF84 |
| Affected twin early biopsy | Caucasian | Male | fALS | *C9orf72* | 37 | RRID:CVCL_ZC74 |
| Affected twin late biopsy | Caucasian | Male | fALS | *C9orf72* | 43 | RRID:CVCL_ZC75 |
| Unaffected twin | Caucasian | Male | Asymptomatic | *C9orf72* | 37 | RRID:CVCL_ZC77 |
| Unaffected father | Caucasian | Male | Asymptomatic | *C9orf72* | 69 | RRID:CVCL_ZC76 |

***Supplementary Table 2*** *– C9orf72 repeat expansion DNA probe sequences.*

| Name | Sequence |
|---|---|
| *C9orf72* Probe AF (Forward) | 5’-AGAACAGGACAAGTTGCC-3’ |
| *C9orf72* Probe AR (Reverse) | 5’-AACACACACCTCCTAAACC-3’ |

***Supplementary Table 3*** *– crRNA guides designed for Cas9 target enrichment.*

| Name | Start | End | Sequence |
|---|---|---|---|
| 2027 | 27572027 | 27572046 | TAACGTAGAATAGAACCCGA |
| 4383 | 27574383 | 27574402 | CCTGCAGACCAAAAGACGCA |

***Supplementary Table 4*** *– List of genes in targeted screening panels. ANG = Genes found in NECTAR panel. ALS2 = Genes found in both panels. All other genes are unique to the ALSgenescanner panel.*

| Gene Name | | | | | | |
|---|---|---|---|---|---|---|
| *1p34-rs3011225* | *CAPN1* | *ERBB4* | *HNRNPA2B1* | *NEK1* | *SIRT1* | *TNFRSF10B* |
| *8p23.2* | *CASP1* | *ERH* | *HSP90B2P* | *NGF* | *SIRT2* | *TRAF3IP2* |
| *ABCG2* | *CAST* | *ERLIN2* | *HSPB8* | *NIPA1* | *SLC12A2* | *TREM2* |
| *ADCY10* | *CAT* | *EWSR1* | *IFNAR1* | *OPTN* | *SLC1A2* | *TTBK2* |
| *AFMID* | *CCNF* | *FAS* | *IGKV5-2* | *PFN1* | *SLC33A1* | *TUBA4A* |
| *ALPK1* | *CDK5* | *FGFR1* | *IL12A* | *PIKFYVE* | *SLC44A2* | *TXNRD1* |
| *ALS2* | *CEP126* | *FGGY* | *IL17A* | *PKN1* | *SLC6A4* | *UBQLN2* |
| *ALS2CR11* | *CHCHD10* | *FIG4* | *INF2* | *PLCG2* | *SMN1* | *UCHL1* |
| *ALS3* | *CHMP2A* | *FOXA2* | *KIF5A* | *PON1* | *SMN2* | *VAC14* |
| *ANG* | *CHMP2B* | *FUS* | *LARGE* | *PPARG* | *SNCA* | *VAPB* |
| *ANXA11* | *CLEC4C* | *G6PD* | *LMNB1* | *PPARGC1A* | *SOD1* | *VCP* |
| *APEX1* | *CNR2* | *GBA2* | *MAL* | *PRH1* | *SPAST* | *VPS51* |
| *ARHGEF28* | *Cyclin* | *GFM1* | *MAPT* | *PRMT1* | *SPG11* | *VPS54* |
| *ATP1A3* | *CYP27A1* | *GRIA2* | *MATR3* | *PRPH* | *SPG20* | *VRK1* |
| *ATXN2* | *CYSLTR1* | *GRN* | *MED13* | *PTGS2* | *SQSTM1* | *VTN* |
| *ATXN3* | *DAO* | *GST* | *MED20* | *RAB5A* | *SS18L1* | *VWA8* |
| *BAG3* | *DCTN1* | *GSTO1* | *MIR206* | *S100B* | *SUGP1* | *XRCC1* |
| *BAX* | *DNA* | *HDAC1* | *MMP9* | *SELP* | *SUMO3* | |
| *BBS9* | *DPP6* | *HDAC6* | *MRLN* | *SEPT9* | *TAF15* | |
| *C5orf42* | *ELP3* | *HFE* | *NAV3* | *SETX* | *TARDBP* | |
| *C9orf72* | *ENPP2* | *HNRNPA1* | *NEFH* | *SIGMAR1* | *TBK1* | |

***Supplementary Table 5*** *– Primary and secondary antibodies for immunocytochemistry.*

| Antibody | Species | Dilution | Supplier | RRID |
|---|---|---|---|---|
| PAX6 | Rabbit | 1:400 | Abcam | AB_305110 |
| Nestin | Mouse | 1:400 | Abcam | AB_444246 |
| CD44 | Rabbit | 1:200 | Abcam | AB_2847859 |
| Vimentin | Chicken | 1:1000 | Millipore | AB_11212377 |
| MAP2 | Guinea Pig | 1:1000 | Synaptic Systems | AB_2138181 |
| Active Caspase 3 | Rabbit | 1:500 | Millipore | AB_91556 |
| Alexa Fluor™ 568 Anti-Rabbit IgG (H+L) | Donkey | 1:1000 | Thermo Fisher Scientific | AB_2534017 |
| Alexa Fluor™ 488 Anti-Mouse IgG (H+L) | Donkey | 1:1000 | Thermo Fisher Scientific | AB_2556542 |
| Cy5 Chicken IgY (H+L) | Goat | 1:1000 | Abcam | AB_10679551 |
| Alexa Fluor™ 647 Anti-Guinea Pig IgG (H+L) | Goat | 1:1000 | Thermo Fisher Scientific | AB_141882 |

***Supplementary Table 6*** *– Targeted sequencing in directly converted iNPCs showing the retention of the C9-HRE in all lines, as well as an intermediate Ataxin-2 repeat expansion (identified both through ATXN2-specific sequencing and through detection of a Poly-Q Indel at protein position Q184 via NECTAR sequencing). A benign synonymous FUS mutation was also identified using the ALS/FTD-specific gene panel. With the larger pan-neuronal mutation panel, an additional five missense mutations were detected in both twins (including the affected twin's late biopsy). Three of these mutations were also observed in the unaffected father, suggesting that the additional mutations in APP and ATP7A were maternally inherited.*

| Family Member | ATXN2/ SCA2 | C9-HRE | Gene | NM Ref | Chromosome | Mutation | Consequence | ClinVar | Project MinE Impact | Risk Score |
|---|---|---|---|---|---|---|---|---|---|---|
| Affected twin/ Unaffected twin | 27 – Intermediate Expansion | 2 + EXP | AARS | 001605.3 | 16 | c.2791G>A p.Gly931Ser | Missense | Benign/Likely benign | Moderate | 25.8 |
| | | | APP | 000484.4 | 21 | c.592T>C p.Ser198Pro | Missense | Conflicting interpretations of pathogenicity: Uncertain significance (1); Benign (2); Likely benign (1) | Moderate | 25.5 |
| | | | ANXA11 | 145868.2 | 10 | c.688C>T p.Arg230Cys | Missense | Benign | Moderate | 33.9 |
| | | | ATP7A | 000052.7 | X | c.2299G>C p.Val767Leu | Missense | Benign | N/A | 25.5 |
| | | | ATXN2 | 002973 | 12 | c.551_552ins ACAGCAGCAGCA p.Q184delinsQQQQQ | Indel | Likely benign | N/A | 0 |

| | | | | | | | | | | |
|---|---|---|---|---|---|---|---|---|---|---|
| | | | FBXO38 | 205836.3 | 5 | c.1394G>A p.Arg465His | Missense | Benign/Likely benign | Moderate | 32.7 |
| | | | FUS | 004960 | 16 | c.66G>A p.Gly22Gly | Synonymous | Benign | Low | 0 |
| Unaffected father | 27 – Intermediate Expansion | 10 + EXP | APP | 000484.4 | 21 | c.592T>C p.Ser198Pro | Missense | Conflicting interpretations of pathogenicity: Uncertain significance (1); Benign (2); Likely benign (1) | Moderate | 25.5 |
| | | | ANXA11 | 145868.2 | 10 | c.688C>T p.Arg230Cys | Missense | Benign | Moderate | 33.9 |
| | | | ATXN2 | 002973 | 12 | c.551_552ins ACAGCAGCAGCA p.Q184delinsQQQQQ | Indel | Likely benign | N/A | 0 |
| | | | FBXO38 | 205836.3 | 5 | c.1394G>A p.Arg465His | Missense | Benign/Likely benign | Moderate | 32.7 |
| | | | FUS | 004960 | 16 | c.66G>A p.Gly22Gly | Synonymous | Benign | Low | 0 |

***Supplementary Table** 7 – Putative risk factors for ALS identified from Illumina whole genome sequencing. 6 discordant variants with a link to neurodegeneration were identified after manual review of ~100 variants of interest.*

| Gene | Variant | Concordance | Disease Link/Function | Risk |
|---|---|---|---|---|
| *TBC1D2B* | F802F (Frameshift) | Discordant | Neurodevelopmental impairment | Predicted as benign |
| *AP3B1* | P863S | Discordant | Protein trafficking to lysosomes | Medium risk |
| *ARSB* | N213D | Discordant | SOD1 neuronal death | Medium risk |
| *DUSP18* | H123R | Discordant | Protein trafficking and ER stress, neuronal disease | Medium risk |
| *PRKCG* | S577A | Discordant | SC ataxia-14 & FTD/ALS | Medium risk |
| *CUL7* | R1411K | Discordant | Ubiquitin-proteasome system | Predicted as benign |

***Supplementary Table 8*** *– Variance of linear regression and number of differentially expressed transcripts in each enrichment comparison. Coefficient of determination ($R^2$) values indicate that less than 1% of the variance is explained by the linear regression, and as such the input data from each comparison is normally distributed and not significantly biased in either direction.*

| Comparison ID | Case | Contrast | $R^2$ | Number of DE | Percentage DE |
|---|---|---|---|---|---|
| AE_vs_UT | affected twin early | unaffected twin | 0.6% | 6869 | 19.2 |
| AE_vs_AL | affected twin early | affected twin late | 0.3% | 6006 | 16.7 |
| AL_vs_UT | affected twin late | unaffected twin | 0.1% | 7983 | 22.2 |

***Supplementary Table 9*** *– Transcript counts (normalised as TPM) for translation initiation factors. The average of the counts from the unaffected twin, affected twin early biopsy and affected twin late biopsy was calculated for each gene. The percentage expression of this average was calculated for each sample, showing a reduction in expression in the affected twin early biopsy.* ***Key*** *=* ***Highest****,* ***Medium****,* ***Lowest****,* ***U % (Unaffected Twin)****,* ***AE % (Affected Twin Early)****,* ***AL % (Affected Twin Late)****.*

| Gene | Symbol | Unaffected Twin | Affected Twin Early | Affected Twin Late | Average | U % | AE % | AL % |
|---|---|---|---|---|---|---|---|---|
| eIF1 | EIF1 | 308.9 | 216.8 | 281.9 | 269.2 | 114.8 | 80.5 | 104.7 |
| Haponin | EIF1AD | 4.2 | 4.4 | 4.1 | 4.2 | 99.6 | 103.6 | 96.7 |
| eIF1α | EIF1AX | 40.8 | 31.6 | 36.1 | 36.2 | 112.8 | 87.4 | 99.8 |
| eIF4C | EIF1AY | 20.7 | 16.8 | 22.9 | 20.1 | 102.9 | 83.6 | 113.5 |
| eIF1B | EIF1B | 53.1 | 32.9 | 38.7 | 41.6 | 127.7 | 79.2 | 93.1 |
| eIF2A | EIF2A | 30.2 | 18.1 | 40.1 | 29.5 | 102.4 | 61.5 | 136 |
| HRI | EIF2AK1 | 31.1 | 33.8 | 39.8 | 34.9 | 89.1 | 96.9 | 114 |
| PKR | EIF2AK2 | 2.6 | 2.1 | 3.2 | 2.6 | 98.9 | 78.5 | 122.6 |
| PERK | EIF2AK3 | 6.1 | 10.4 | 4.9 | 7.2 | 85.3 | 145.7 | 69 |

| | | | | | | | | |
|---|---|---|---|---|---|---|---|---|
| GCN2 | EIF2AK4 | 5.8 | 14 | 17.6 | 12.5 | 46.5 | 112.5 | 141 |
| eIF2Bα | EIF2B1 | 55.2 | 34.2 | 44.3 | 44.6 | 123.9 | 76.7 | 99.4 |
| eIF2Bβ | EIF2B2 | 26.4 | 15.4 | 24.3 | 22 | 119.8 | 69.8 | 110.4 |
| eIF2Bγ | EIF2B3 | 43.3 | 25.7 | 27.8 | 32.3 | 134.2 | 79.7 | 86.2 |
| eIF2Bδ | EIF2B4 | 35.6 | 27.4 | 32.5 | 31.8 | 112 | 86 | 102.1 |
| eIF2Bε | EIF2B5 | 34.2 | 25.5 | 42.8 | 34.2 | 100.1 | 74.8 | 125.1 |
| Ligatin | EIF2D | 59.2 | 40.8 | 63.7 | 54.6 | 108.6 | 74.8 | 116.7 |
| eIF2α | EIF2S1 | 108.7 | 54.6 | 84.9 | 82.7 | 131.4 | 66 | 102.6 |
| eIF2β | EIF2S2 | 171.3 | 120.6 | 143.1 | 145 | 118.2 | 83.2 | 98.7 |
| eIF2γ | EIF2S3 | 61.1 | 63.7 | 81 | 68.6 | 89.1 | 92.9 | 118.1 |
| eIF3θ | EIF3A | 33.9 | 41.4 | 57.4 | 44.2 | 76.6 | 93.6 | 129.8 |
| eIF3S9 | EIF3B | 77.3 | 55.8 | 88.8 | 74 | 104.5 | 75.5 | 120.1 |
| eIF3S8 | EIF3C | 386.9 | 238 | 146.8 | 257.2 | 150.4 | 92.5 | 57.1 |
| eIF3ζ | EIF3D | 241.6 | 175.7 | 274.7 | 230.7 | 104.7 | 76.2 | 119.1 |
| eIF3S6 | EIF3E | 138.5 | 109.4 | 198.3 | 148.7 | 93.1 | 73.6 | 133.3 |
| eIF3ε | EIF3F | 322.3 | 235.6 | 347.5 | 301.8 | 106.8 | 78.1 | 115.2 |
| eIF3δ | EIF3G | 290.6 | 228.9 | 362.2 | 293.9 | 98.9 | 77.9 | 123.2 |
| eIF3γ | EIF3H | 166.1 | 134.8 | 212.6 | 171.2 | 97 | 78.7 | 124.2 |

| | | | | | | | | |
|---|---|---|---|---|---|---|---|---|
| eIF3β | EIF3I | 399.8 | 289.7 | 462.8 | 384.1 | 104.1 | 75.4 | 120.5 |
| eIF3α | EIF3J | 60.9 | 51.1 | 52.1 | 54.7 | 111.4 | 93.4 | 95.3 |
| eIF3S12 | EIF3K | 494.1 | 319.6 | 442.5 | 418.8 | 118 | 76.3 | 105.7 |
| eIF3S6IP | EIF3L | 208.1 | 185.8 | 238.5 | 210.8 | 98.7 | 88.1 | 113.1 |
| PCID1 | EIF3M | 148.3 | 87.2 | 118.3 | 117.9 | 125.8 | 73.9 | 100.3 |
| DDX2A | EIF4A1 | 1088.5 | 891.7 | 1269.7 | 1083.3 | 100.5 | 82.3 | 117.2 |
| DDX2B | EIF4A2 | 23 | 109.9 | 178.5 | 103.8 | 22.2 | 105.9 | 171.9 |
| DDX48 | EIF4A3 | 145.9 | 189.3 | 192.3 | 175.8 | 83 | 107.7 | 109.4 |
| eIF4B | EIF4B | 128.2 | 112.8 | 190.9 | 144 | 89.1 | 78.3 | 132.6 |
| eIF4E | EIF4E | 27.9 | 18.4 | 23.9 | 23.4 | 119.1 | 78.8 | 102.1 |
| eIF4E2 | EIF4E2 | 93.2 | 71.3 | 86.3 | 83.6 | 111.5 | 85.3 | 103.2 |
| eIF4E3 | EIF4E3 | 0.4 | 1.1 | 1.4 | 1 | 38.4 | 118.9 | 142.7 |
| eIF4γ1 | EIF4G1 | 140 | 98.9 | 52.2 | 97 | 144.3 | 101.9 | 53.8 |
| eIF4γ2 | EIF4G2 | 97.9 | 159.5 | 255.5 | 171 | 57.3 | 93.3 | 149.4 |
| eIF4γ3 | EIF4G3 | 5.2 | 11.7 | 15.4 | 10.8 | 48.4 | 108.8 | 142.7 |
| eIF4H | EIF4H | 200.9 | 171.2 | 250.7 | 207.6 | 96.8 | 82.5 | 120.7 |
| eIF5 | EIF5 | 59.3 | 27.6 | 34.4 | 40.4 | 146.7 | 68.3 | 85 |
| eIF5α | EIF5A | 1234.8 | 859 | 1168.7 | 1087.5 | 113.5 | 79 | 107.5 |

| | | | | | | | | |
|---|---|---|---|---|---|---|---|---|
| eIF5α2 | EIF5A2 | 2.1 | 1.2 | 1.4 | 1.6 | 136.5 | 74.3 | 89.1 |
| IF2 | EIF5B | 70.2 | 40.8 | 36.2 | 49.1 | 143 | 83.2 | 73.8 |
| P27BBP | EIF6 | 186.7 | 124.7 | 210.5 | 174 | 107.3 | 71.7 | 121 |
| GCN1 Activator Of EIF2AK4 | GCN1 | 12.3 | 14.2 | 17.2 | 14.6 | 84.6 | 97.3 | 118.1 |
| Poly(A) Binding Protein Cytoplasmic 1 | PPP1R15A | 24 | 18.9 | 30.3 | 24.4 | 98.3 | 77.5 | 124.2 |
| Poly(A) Binding Protein Cytoplasmic 4 | PABPC1 | 517.7 | 979.9 | 980.2 | 825.9 | 62.7 | 118.6 | 118.7 |
| Poly(A) Binding Protein Nuclear 1 | PABPC4 | 71 | 100.2 | 117.1 | 96.1 | 73.9 | 104.3 | 121.8 |
| GADD34 | PABPN1 | 48.9 | 46.4 | 77.5 | 57.6 | 84.9 | 80.5 | 134.6 |

***Supplementary Table 10*** *– Transcript counts (normalised as TPM) for ion homeostasis factors. A gene list was developed from 5 GO terms: GO:0006812 (monoatomic cation transport), GO:0030003 (cellular cation homeostasis), GO:0005261 (cation channel activity), GO:0006813 (potassium ion transport), and GO:0005921 (gap junction). The average of the counts from the unaffected twin, affected twin early biopsy and affected twin late biopsy was calculated for each gene. The percentage expression of this average was calculated for each sample and transcripts were ordered from highest average expression.* ***Key*** *=* ***Highest****,* ***Medium****,* ***Lowest****,* ***U % (Unaffected Twin), AE % (Affected Twin Early), AL % (Affected Twin Late)****.*

| Gene | Symbol | Unaffected Twin | Affected Twin Early | Affected Twin Late | Average | U % | AE % | AL % | GO Term |
|---|---|---|---|---|---|---|---|---|---|
| Dysadherin | FXYD5 | 728.0 | 886.7 | 325.2 | 646.6 | 112.6 | 137.1 | 50.3 | GO:0006813 |
| Cellular Communication Network Factor 3 | CCN3 | 325.2 | 0.4 | 688.1 | 337.9 | 96.2 | 0.1 | 203.6 | GO:0005921 |
| Drebrin 1 | DBN1 | 119.6 | 292.8 | 228.8 | 213.7 | 56.0 | 137.0 | 107.1 | GO:0005921 |

| | | | | | | | | | |
|---|---|---|---|---|---|---|---|---|---|
| Solute Carrier Family 39 Member 1 | SLC39A1 | 126.1 | 157.2 | 107.9 | 130.4 | 96.7 | 120.6 | 82.8 | GO:0006812 |
| Connexin-43 | GJA1 | 59.9 | 129.8 | 78.0 | 89.2 | 67.1 | 145.5 | 87.5 | GO:0005921 |
| Multiple Inositol-Polyphosphate Phosphatase 1 | MINPP1 | 148.2 | 36.3 | 34.4 | 73.0 | 203.1 | 49.8 | 47.1 | GO:0030003 |
| Solute Carrier Family 39 Member 6 | SLC39A6 | 112.0 | 51.8 | 49.8 | 71.2 | 157.3 | 72.8 | 69.9 | GO:0030003 |
| Potassium Calcium-Activated Channel Subfamily M Alpha 1 | KCNMA1 | 108.3 | 26.7 | 66.3 | 67.1 | 161.4 | 39.8 | 98.8 | GO:0006813 |
| Mucolipin-1 | MCOLN1 | 85.6 | 62.2 | 42.8 | 63.5 | 134.7 | 97.9 | 67.4 | GO:0005261<br>GO:0006812 |
| Piezo Type Mechanosensitive Ion Channel Component 1 | PIEZO1 | 64.1 | 66.9 | 12.1 | 47.7 | 134.3 | 140.3 | 25.4 | GO:0005261<br>GO:0006812 |
| Solute Carrier Family 39 Member 14 | SLC39A14 | 35.7 | 52.3 | 40.0 | 42.7 | 83.7 | 122.6 | 93.7 | GO:0030003 |
| Calcineurin B homologous protein 1 | CHP1 | 44.8 | 25.9 | 34.7 | 35.1 | 127.5 | 73.8 | 98.8 | GO:0006813 |
| N-Ethylmaleimide Sensitive Factor | NSF | 17.7 | 21.9 | 49.8 | 29.8 | 59.4 | 73.5 | 167.1 | GO:0006813 |
| Calcium Homeostasis Modulator Family Member 2 | CALHM2 | 23.0 | 17.5 | 27.4 | 22.6 | 101.7 | 77.2 | 121.1 | GO:0005261 |
| Vacuolar Protein Sorting 4 Homolog B | VPS4B | 23.7 | 18.5 | 24.9 | 22.4 | 105.9 | 82.7 | 111.4 | GO:0006813 |
| Connexin-31 | GJB3 | 50.3 | 1.3 | 5.9 | 19.2 | 262.5 | 6.7 | 30.8 | GO:0005921 |
| Small G Protein Signalling Modulator 3 | SGSM3 | 14.3 | 24.5 | 16.8 | 18.5 | 77.3 | 132.0 | 90.7 | GO:0005921 |
| ATPase Cation Transporting 13A2 | ATP13A2 | 12.6 | 30.6 | 11.2 | 18.1 | 69.6 | 168.6 | 61.8 | GO:0030003 |
| Pannexin 1 | PANX1 | 15.5 | 16.9 | 15.9 | 16.1 | 96.1 | 105.2 | 98.7 | GO:0005921<br>GO:0006812 |
| Polycystin 2 | PKD2 | 9.2 | 20.4 | 14.4 | 14.7 | 62.9 | 139.1 | 98.0 | GO:0005261 |

| | | | | | | | | | |
|---|---|---|---|---|---|---|---|---|---|
| Solute Carrier Family 39 Member 8 | SLC39A8 | 31.5 | 3.6 | 8.2 | 14.4 | 218.2 | 24.8 | 57.0 | GO:0030003 |
| Solute Carrier Family 39 Member 10 | SLC39A10 | 7.3 | 19.2 | 14.7 | 13.7 | 53.1 | 139.8 | 107.1 | GO:0030003 |
| Cytochrome C Oxidase Copper Chaperone | COX11 | 19.8 | 8.0 | 12.4 | 13.4 | 148.0 | 59.6 | 92.4 | GO:0030003 |
| Solute Carrier Family 39 Member 4 | SLC39A4 | 7.1 | 16.5 | 14.0 | 12.5 | 56.4 | 131.8 | 111.8 | GO:0030003 |
| TWIK-2 | KCNK6 | 11.0 | 19.3 | 4.9 | 11.7 | 94.0 | 164.5 | 41.5 | GO:0006813 |
| Cyclin-Dependent Kinase Inhibitor 1B | CDKN1B | 8.3 | 7.0 | 10.6 | 8.6 | 96.1 | 81.4 | 122.5 | GO:0006813 |
| Cyclin-dependent kinase 2 | CDK2 | 7.5 | 6.8 | 9.8 | 8.0 | 93.3 | 84.9 | 121.8 | GO:0006813 |
| Potassium Calcium-Activated Channel Subfamily N Member 4 | KCNN4 | 4.6 | 1.8 | 16.7 | 7.7 | 59.8 | 23.2 | 217.0 | GO:0006813 |
| Cytospin A | SPECC1L | 6.4 | 6.7 | 7.9 | 7.0 | 91.5 | 95.2 | 113.3 | GO:0005921 |
| Transient Receptor Potential Vanilloid 2 | TRPV2 | 5.9 | 10.5 | 0.9 | 5.8 | 102.1 | 182.1 | 15.8 | GO:0005261 |
| Transient Receptor Potential Vanilloid 4 | TRPV4 | 8.2 | 4.9 | 3.8 | 5.6 | 145.7 | 86.8 | 67.5 | GO:0005261 |
| Connexin-26 | GJB2 | 14.8 | 0.6 | 0.9 | 5.4 | 272.6 | 10.3 | 17.1 | GO:0005921 |
| Solute Carrier Family 4 Member 11 | SLC4A11 | 6.8 | 5.9 | 2.8 | 5.2 | 131.1 | 114.3 | 54.6 | GO:0030003 |
| Calcium Homeostasis Modulator 3 | CALHM3 | 3.4 | 6.1 | 4.1 | 4.5 | 73.9 | 134.8 | 91.3 | GO:0005261 |
| Polycystin 1 | PKD1 | 3.4 | 5.3 | 4.3 | 4.3 | 79.1 | 121.0 | 99.8 | GO:0005261 |
| Nitric Oxide Synthase 3 | NOS3 | 7.3 | 3.7 | 1.8 | 4.3 | 171.5 | 86.3 | 42.1 | GO:0006813 |
| Tight Junction Protein ZO-1 | TJP1 | 5.3 | 3.0 | 3.5 | 3.9 | 134.2 | 76.8 | 89.0 | GO:0005921 |
| ATPase Phospholipid Transporting 10D | ATP10D | 2.3 | 3.8 | 5.0 | 3.7 | 62.4 | 102.7 | 134.9 | GO:0006812 |
| Connexin-47 | GJC2 | 3.1 | 4.5 | 2.7 | 3.4 | 91.4 | 129.8 | 78.9 | GO:0005921 |

| | | | | | | | | | |
|---|---|---|---|---|---|---|---|---|---|
| Testis Expressed Metallothionein Like Protein | TESMIN | 2.8 | 3.2 | 3.4 | 3.1 | 88.3 | 104.1 | 107.6 | GO:0030003 |
| FXYD Domain Containing Ion Transport Regulator 3 | FXYD3 | 6.2 | 0.0 | 3.0 | 3.1 | 202.2 | 0.0 | 97.8 | GO:0006813 |
| Cyclin And CBS Domain Divalent Metal Cation Transport Mediator 4 | CNNM4 | 2.1 | 3.4 | 2.0 | 2.5 | 83.5 | 137.0 | 79.5 | GO:0030003 |
| Potassium Voltage-Gated Channel Subfamily D Member 3 | KCND3 | 0.3 | 2.5 | 3.6 | 2.1 | 11.7 | 118.5 | 169.8 | GO:0006813 |
| Potassium Voltage-Gated Channel Modifier Subfamily V Member 1 | KCNV1 | 0.4 | 3.0 | 2.3 | 1.9 | 22.5 | 155.2 | 122.3 | GO:0006813 |
| Potassium Two Pore Domain Channel Subfamily K Member 1 | KCNK1 | 2.1 | 2.7 | 0.5 | 1.8 | 119.2 | 152.9 | 27.9 | GO:0006813 |
| Transient Receptor Potential Cation Channel Subfamily C Member 1 | TRPC1 | 2.0 | 1.7 | 1.6 | 1.7 | 112.8 | 95.0 | 92.2 | GO:0005261 |
| Tuberous sclerosis 1 | TSC1 | 1.7 | 2.1 | 1.2 | 1.7 | 100.2 | 127.4 | 72.4 | GO:0006813 |
| Aquaporin 1 | AQP1 | 0.4 | 0.7 | 3.8 | 1.6 | 22.0 | 42.1 | 236.0 | GO:0006813 |
| Cholinergic Receptor Nicotinic Beta 1 | CHRNB1 | 1.4 | 1.4 | 1.7 | 1.5 | 94.0 | 94.0 | 111.9 | GO:0006812 |
| Transient Receptor Potential Cation Channel Subfamily C Member 6 | TRPC6 | 0.5 | 2.5 | 0.9 | 1.3 | 40.9 | 190.9 | 68.2 | GO:0005261<br>GO:0006812 |
| Connexin-45 | GJC1 | 2.2 | 0.8 | 0.8 | 1.2 | 174.2 | 61.3 | 64.5 | GO:0005921 |
| Pannexin 2 | PANX2 | 0.8 | 2.1 | 0.8 | 1.2 | 62.8 | 171.1 | 66.1 | GO:0006812 |
| Potassium Inwardly Rectifying Channel Subfamily J Member 12 | KCNJ12 | 1.4 | 1.5 | 0.6 | 1.2 | 119.3 | 131.4 | 49.3 | GO:0006813 |

| | | | | | | | | | |
|---|---|---|---|---|---|---|---|---|---|
| FXYD Domain Containing Ion Transport Regulator 6 | FXYD6 | 0.2 | 3.0 | 0.1 | 1.1 | 22.1 | 273.3 | 4.6 | GO:0006813 |
| FXYD Domain Containing Ion Transport Regulator 1 | FXYD1 | 1.9 | 0.0 | 1.3 | 1.0 | 179.0 | 0.0 | 121.0 | GO:0006813 |
| Potassium Two Pore Domain Channel Subfamily K Member 5 | KCNK5 | 1.5 | 1.1 | 0.3 | 1.0 | 152.5 | 115.2 | 32.3 | GO:0006813 |
| Potassium Inwardly Rectifying Channel Subfamily J Member 3 | KCNJ3 | 0.4 | 0.0 | 2.5 | 0.9 | 38.3 | 0.0 | 261.7 | GO:0006813 |
| Cyclic Nucleotide-Gated Channel Alpha 3 | CNGA3 | 1.2 | 1.4 | 0.2 | 0.9 | 125.6 | 151.6 | 22.7 | GO:0006812 |
| Calcium Homeostasis Modulator Family Member 5 | CALHM5 | 1.6 | 0.6 | 0.4 | 0.8 | 187.8 | 65.0 | 47.2 | GO:0005261 |
| ATPase Na+/K+ Transporting Subunit Alpha 2 | ATP1A2 | 2.1 | 0.1 | 0.3 | 0.8 | 253.8 | 9.5 | 36.8 | GO:0006813 |
| Potassium Inwardly Rectifying Channel Subfamily J Member 15 | KCNJ15 | 0.9 | 0.3 | 0.2 | 0.5 | 193.6 | 61.7 | 44.7 | GO:0006813 |
| Potassium Voltage-Gated Channel Subfamily A Regulatory Beta Subunit 3 | KCNAB3 | 0.5 | 0.0 | 0.4 | 0.3 | 155.2 | 10.3 | 134.5 | GO:0006813 |
| Potassium Calcium-Activated Channel Subfamily M Regulatory Beta Subunit 1 | KCNMB1 | 0.1 | 0.6 | 0.0 | 0.2 | 28.4 | 255.4 | 16.2 | GO:0006813 |
| Transmembrane Protein 63C | TMEM63C | 0.0 | 0.1 | 0.0 | 0.1 | 33.3 | 200.0 | 66.7 | GO:0006812 |
| Calbindin 2 | CALB2 | 0.0 | 0.1 | 0.0 | 0.0 | 0.0 | 250.0 | 50.0 | GO:0005921 |

***Supplementary Table 11*** *– Transcript counts (normalised as TPM) for putative neuroprotective targets identified in Figs. 6B and C. This list was ordered by fold change, with downregulated (i.e. increased in unaffected twin) terms then ranked by decreasing p-value. A gene list was developed from the top 5 significantly downregulated GO terms in this list: GO:0045862 (positive regulation of proteolysis), GO:0031089 (platelet dense granule lumen), GO:0030021 (extracellular matrix structural constituent conferring compression resistance), GO:1903077 (negative regulation of protein localization to plasma membrane), and GO:0030667 (secretory granule membrane). The average of the counts from the unaffected twin and affected twin early biopsy were calculated for each transcript. The percentage expression of this average was calculated for each sample and transcripts were ordered by increasing negative log2FC. Key =* ***U % (Unaffected Twin), AE % (Affected Twin Early)****.*

| Symbol | Unaffected Twin | Affected Twin Early | Average | U % | AE % | log2FC | GO Term | GO Name | GO P-Value |
|---|---|---|---|---|---|---|---|---|---|
| MMP14 | 286.15 | 0.26 | 143.2 | 199.8 | 0.2 | -10.1 | GO:0045862 | positive regulation of proteolysis | 0.003 |
| CD14 | 16.88 | 0.03 | 8.5 | 199.6 | 0.4 | -9.1 | GO:0030667 | secretory granule membrane | 0.001 |
| PKP1 | 6.7 | 0.05 | 3.4 | 198.5 | 1.5 | -7.0 | GO:0030667 | secretory granule membrane | 0.001 |
| DCN | 184 | 1.96 | 93.0 | 197.9 | 2.1 | -6.6 | GO:0030021 | extracellular matrix structural constituent conferring compression resistance | 0.010 |
| CXCR2 | 4.14 | 0.07 | 2.1 | 196.7 | 3.3 | -5.9 | GO:0030667 | secretory granule membrane | 0.001 |
| SELENOP | 3.75 | 0.12 | 1.9 | 193.8 | 6.2 | -5.0 | GO:0031089 | platelet dense granule lumen | 0.008 |
| NUPR1 | 73.57 | 3.92 | 38.7 | 189.9 | 10.1 | -4.2 | GO:0045862 | positive regulation of proteolysis | 0.003 |
| RARRES2 | 528.44 | 31.61 | 280.0 | 188.7 | 11.3 | -4.1 | GO:0031089 | platelet dense granule lumen | 0.008 |
| CASP1 | 4.47 | 0.27 | 2.4 | 188.6 | 11.4 | -4.1 | GO:0045862 | positive regulation of proteolysis | 0.003 |
| SIRPA | 36.52 | 2.52 | 19.5 | 187.1 | 12.9 | -3.9 | GO:0030667 | secretory granule membrane | 0.001 |
| BGN | 14.61 | 1.03 | 7.8 | 186.8 | 13.2 | -3.8 | GO:0030021 | extracellular matrix structural constituent | 0.010 |

| | | | | | | | | | |
|---|---|---|---|---|---|---|---|---|---|
| | | | | | | | | conferring compression resistance | |
| SRC | 38.8 | 2.77 | 20.8 | 186.7 | 13.3 | -3.8 | GO:0045862 | positive regulation of proteolysis | 0.003 |
| CLEC3B | 2498.97 | 196.94 | 1348.0 | 185.4 | 14.6 | -3.7 | GO:0045862, GO:0031089 | positive regulation of proteolysis, platelet dense granule lumen | 0.003, 0.008 |
| FMOD | 46.61 | 3.95 | 25.3 | 184.4 | 15.6 | -3.6 | GO:0030021 | extracellular matrix structural constituent conferring compression resistance | 0.010 |
| NGFR | 4.51 | 0.4 | 2.5 | 183.7 | 16.3 | -3.5 | GO:0045862 | positive regulation of proteolysis | 0.003 |
| LRRC15 | 39.7 | 3.73 | 21.7 | 182.8 | 17.2 | -3.4 | GO:1903077 | negative regulation of protein localization to plasma membrane | 0.011 |
| IRAG2 | 13.65 | 1.72 | 7.7 | 177.6 | 22.4 | -3.0 | GO:0030667 | secretory granule membrane | 0.001 |
| ECM1 | 549.58 | 71.4 | 310.5 | 177.0 | 23.0 | -2.9 | GO:0031089 | platelet dense granule lumen | 0.008 |
| PRELP | 10.58 | 1.39 | 6.0 | 176.8 | 23.2 | -2.9 | GO:0030021 | extracellular matrix structural constituent conferring compression resistance | 0.010 |
| CASP10 | 1.31 | 0.18 | 0.7 | 175.8 | 24.2 | -2.8 | GO:0045862 | positive regulation of proteolysis | 0.003 |
| LUM | 24.05 | 3.51 | 13.8 | 174.5 | 25.5 | -2.8 | GO:0030021 | extracellular matrix structural constituent conferring compression resistance | 0.010 |
| TNF | 1.03 | 0.16 | 0.6 | 173.1 | 26.9 | -2.7 | GO:0045862 | positive regulation of proteolysis | 0.003 |

| | | | | | | | | | |
|---|---|---|---|---|---|---|---|---|---|
| UBR4 | 43.89 | 6.94 | 25.4 | 172.7 | 27.3 | -2.7 | GO:0030667 | secretory granule membrane | 0.001 |
| BST2 | 6.95 | 1.12 | 4.0 | 172.2 | 27.8 | -2.6 | GO:0030667 | secretory granule membrane | 0.001 |
| ALDH3B1 | 25.52 | 4.15 | 14.8 | 172.0 | 28.0 | -2.6 | GO:0030667 | secretory granule membrane | 0.001 |
| ASPN | 224.02 | 37.56 | 130.8 | 171.3 | 28.7 | -2.6 | GO:0030021 | extracellular matrix structural constituent conferring compression resistance | 0.010 |
| HLA-B | 2173 | 385.47 | 1279.2 | 169.9 | 30.1 | -2.5 | GO:0030667 | secretory granule membrane | 0.001 |
| PCOLCE | 893.36 | 158.48 | 525.9 | 169.9 | 30.1 | -2.5 | GO:0045862 | positive regulation of proteolysis | 0.003 |
| RNF180 | 1.25 | 0.23 | 0.7 | 168.9 | 31.1 | -2.5 | GO:0045862 | positive regulation of proteolysis | 0.003 |
| IL1B | 5.97 | 1.12 | 3.5 | 168.4 | 31.6 | -2.4 | GO:0045862 | positive regulation of proteolysis | 0.003 |
| NLRP1 | 2.96 | 0.58 | 1.8 | 167.2 | 32.8 | -2.4 | GO:0045862 | positive regulation of proteolysis | 0.003 |
| TICAM2 | 1.49 | 0.3 | 0.9 | 166.5 | 33.5 | -2.3 | GO:0030667 | secretory granule membrane | 0.001 |
| IFI16 | 28.93 | 5.95 | 17.4 | 165.9 | 34.1 | -2.3 | GO:0045862 | positive regulation of proteolysis | 0.003 |
| MAP3K5 | 0.82 | 0.17 | 0.5 | 165.7 | 34.3 | -2.3 | GO:0045862 | positive regulation of proteolysis | 0.003 |
| IL33 | 1.52 | 0.32 | 0.9 | 165.2 | 34.8 | -2.3 | GO:0045862 | positive regulation of proteolysis | 0.003 |
| AXIN2 | 0.7 | 0.15 | 0.4 | 164.7 | 35.3 | -2.2 | GO:0045862 | positive regulation of proteolysis | 0.003 |
| MOXD1 | 53.45 | 11.55 | 32.5 | 164.5 | 35.5 | -2.2 | GO:0030667 | secretory granule membrane | 0.001 |
| ANPEP | 41.33 | 9.21 | 25.3 | 163.6 | 36.4 | -2.2 | GO:0030667 | secretory granule membrane | 0.001 |

| | | | | | | | | | |
|---|---|---|---|---|---|---|---|---|---|
| GSN | 290.85 | 64.99 | 177.9 | 163.5 | 36.5 | -2.2 | GO:0045862 | positive regulation of proteolysis | 0.003 |
| PCOLCE2 | 140.12 | 33.59 | 86.9 | 161.3 | 38.7 | -2.1 | GO:0045862 | positive regulation of proteolysis | 0.003 |
| SERPINB6 | 1175.51 | 298.17 | 736.8 | 159.5 | 40.5 | -2.0 | GO:0030667 | secretory granule membrane | 0.001 |
| TNFRSF1B | 2.52 | 0.65 | 1.6 | 159.0 | 41.0 | -2.0 | GO:0030667 | secretory granule membrane | 0.001 |
| GPER1 | 17.36 | 4.49 | 10.9 | 158.9 | 41.1 | -2.0 | GO:0045862 | positive regulation of proteolysis | 0.003 |
| TNFRSF1B | 2.52 | 0.65 | 1.6 | 159.0 | 41.0 | -2.0 | GO:0045862 | positive regulation of proteolysis | 0.003 |
| ITPR1 | 13.13 | 3.47 | 8.3 | 158.2 | 41.8 | -1.9 | GO:0030667 | secretory granule membrane | 0.001 |
| HLA-C | 1407.21 | 388.71 | 898.0 | 156.7 | 43.3 | -1.9 | GO:0030667 | secretory granule membrane | 0.001 |
| CD55 | 178.61 | 55.48 | 117.0 | 152.6 | 47.4 | -1.7 | GO:0030667 | secretory granule membrane | 0.001 |
| RGMA | 1.91 | 0.59 | 1.3 | 152.8 | 47.2 | -1.7 | GO:0045862 | positive regulation of proteolysis | 0.003 |
| CPE | 416.09 | 130 | 273.0 | 152.4 | 47.6 | -1.7 | GO:0030667 | secretory granule membrane | 0.001 |
| VSIR | 34.42 | 10.77 | 22.6 | 152.3 | 47.7 | -1.7 | GO:0045862 | positive regulation of proteolysis | 0.003 |
| CD68 | 84.9 | 27.75 | 56.3 | 150.7 | 49.3 | -1.6 | GO:0030667 | secretory granule membrane | 0.001 |
| RHBDD1 | 5.34 | 1.78 | 3.6 | 150.0 | 50.0 | -1.6 | GO:0045862 | positive regulation of proteolysis | 0.003 |
| PID1 | 11.11 | 3.81 | 7.5 | 148.9 | 51.1 | -1.5 | GO:1903077 | negative regulation of protein localization to plasma membrane | 0.011 |
| ANXA7 | 94.99 | 33.17 | 64.1 | 148.2 | 51.8 | -1.5 | GO:0030667 | secretory granule membrane | 0.001 |

| | | | | | | | | | |
|---|---|---|---|---|---|---|---|---|---|
| CFLAR | 3.33 | 1.18 | 2.3 | 147.7 | 52.3 | -1.5 | GO:0045862 | positive regulation of proteolysis | 0.003 |
| PSMC2 | 63.91 | 22.83 | 43.4 | 147.4 | 52.6 | -1.5 | GO:0045862 | positive regulation of proteolysis | 0.003 |
| FBLN1 | 30.23 | 11.62 | 20.9 | 144.5 | 55.5 | -1.4 | GO:0045862 | positive regulation of proteolysis | 0.003 |
| CADPS | 1.23 | 0.48 | 0.9 | 143.9 | 56.1 | -1.4 | GO:0030667 | secretory granule membrane | 0.001 |
| PSME2 | 140.36 | 57.03 | 98.7 | 142.2 | 57.8 | -1.3 | GO:0045862 | positive regulation of proteolysis | 0.003 |
| HSPA1A | 29.84 | 12.19 | 21.0 | 142.0 | 58.0 | -1.3 | GO:0045862 | positive regulation of proteolysis | 0.003 |
| CD58 | 166.48 | 68.09 | 117.3 | 141.9 | 58.1 | -1.3 | GO:0030667 | secretory granule membrane | 0.001 |
| MAGEF1 | 66.2 | 27.63 | 46.9 | 141.1 | 58.9 | -1.3 | GO:0045862 | positive regulation of proteolysis | 0.003 |
| FBXW7 | 18 | 7.76 | 12.9 | 139.8 | 60.2 | -1.2 | GO:0045862 | positive regulation of proteolysis | 0.003 |
| CYB561 | 4.51 | 1.97 | 3.2 | 139.2 | 60.8 | -1.2 | GO:0030667 | secretory granule membrane | 0.001 |
| VPS13C | 2.12 | 0.96 | 1.5 | 137.7 | 62.3 | -1.1 | GO:0030667 | secretory granule membrane | 0.001 |
| LHFPL2 | 20.35 | 9.23 | 14.8 | 137.6 | 62.4 | -1.1 | GO:0030667 | secretory granule membrane | 0.001 |
| DET1 | 1.75 | 0.8 | 1.3 | 137.3 | 62.7 | -1.1 | GO:0045862 | positive regulation of proteolysis | 0.003 |
| GBA1 | 280.81 | 128.29 | 204.6 | 137.3 | 62.7 | -1.1 | GO:0045862 | positive regulation of proteolysis | 0.003 |
| RHOF | 16.38 | 7.61 | 12.0 | 136.6 | 63.4 | -1.1 | GO:0030667 | secretory granule membrane | 0.001 |
| SPACA6 | 0.76 | 0.36 | 0.6 | 135.7 | 64.3 | -1.1 | GO:0030667 | secretory granule membrane | 0.001 |
| HLA-H | 79.7 | 37.37 | 58.5 | 136.2 | 63.8 | -1.1 | GO:0030667 | secretory granule membrane | 0.001 |

| | | | | | | | | | |
|---|---|---|---|---|---|---|---|---|---|
| SMAD3 | 7.91 | 3.73 | 5.8 | 135.9 | 64.1 | -1.1 | GO:0045862 | positive regulation of proteolysis | 0.003 |
| GABARAP | 336 | 161.41 | 248.7 | 135.1 | 64.9 | -1.1 | GO:0045862 | positive regulation of proteolysis | 0.003 |
| ZFAND2A | 42.1 | 21.04 | 31.6 | 133.4 | 66.6 | -1.0 | GO:0045862 | positive regulation of proteolysis | 0.003 |

***Supplementary Table 12*** *– iNPC-derived astrocytes produce cytoplasmic inclusions positive for TDP-43. Following blinded manual quantification, astrocytes from the Unaffected Twin had the highest proportion of cells with TDP-43. Values = mean from three biological experiments.*

| Cell Line | Cells with TDP-43 inclusions (%) |
|---|---|
| healthy control | 2.1 |
| C9-ALS | 4.5 |
| affected twin early | 7.5 |
| affected twin late | 2.9 |
| unaffected father | 5.0 |
| unaffected twin | 13.6 |

### Supplementary Figures

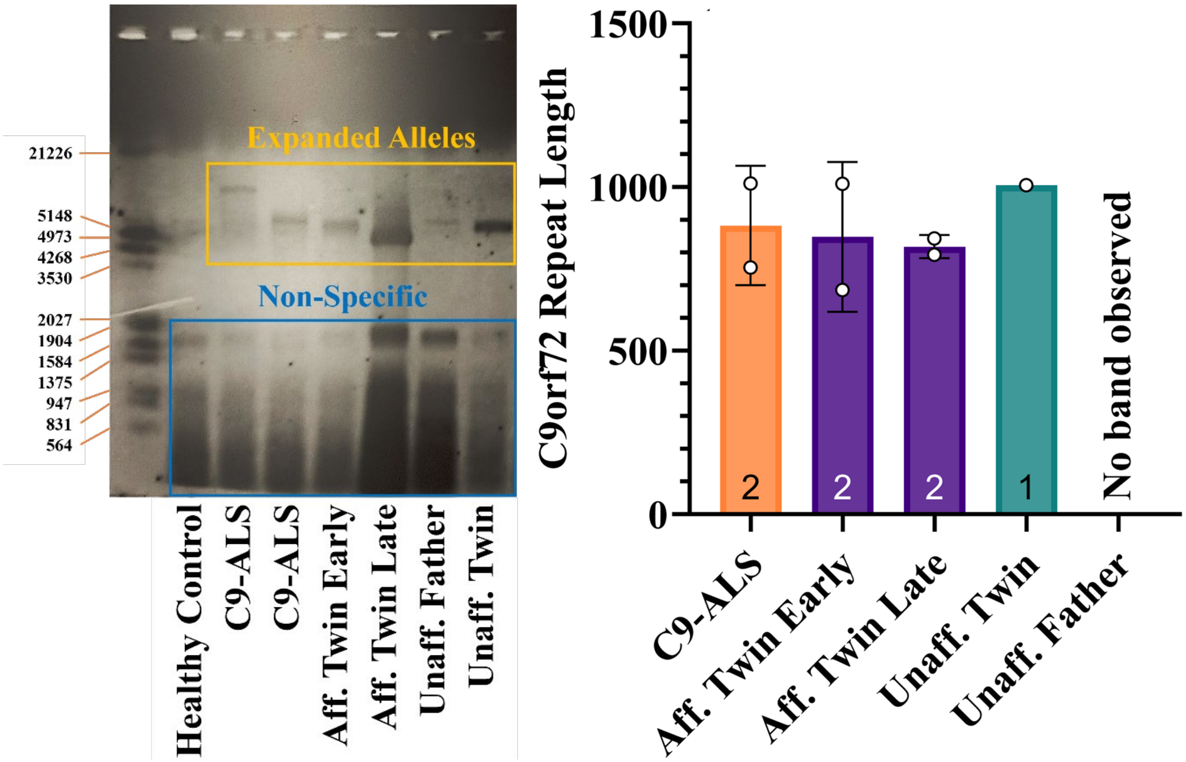


***Supplementary Figure 1*** - *C9orf72 hexanucleotide repeat expansion size was determined using Southern blotting, performed on gDNA extracted from directly converted iNPCs from the family. An expanded allele was established in iNPCs generated by both symptomatic (early and late biopsy) and asymptomatic twins (gold box), while no clear expanded band was observed in the asymptomatic father (left). Repeat length was quantified, showing comparable sizing in the twins, as well as unrelated C9-ALS iNPCs (right). Two biological repeats per line (except unaffected twin n = 1).*

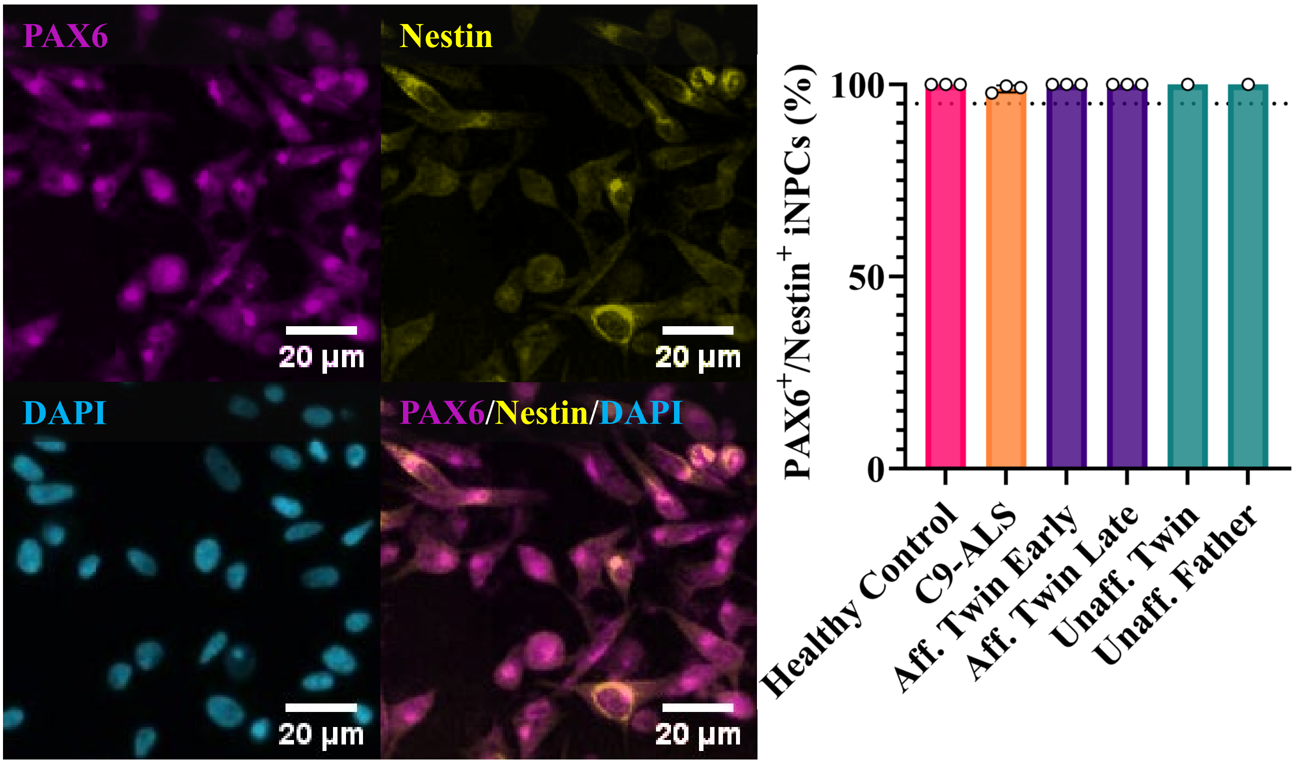


***Supplementary Figure 2*** *– Representative images of affected twin late iNPCs expressing neural progenitor markers PAX6 and Nestin (left). When quantified, all iNPC lines used in this study co-expressed PAX6 and Nestin in >95% of cells (dotted line = 95% percentile). Three biological repeats per line (except unaffected twin and unaffected father n = 1).*

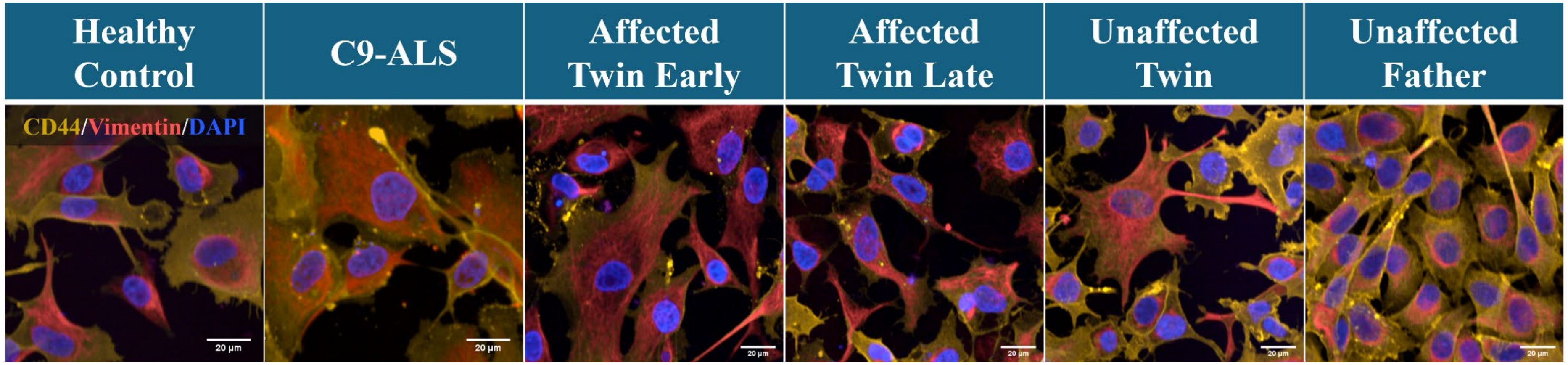


***Supplementary Figure 3*** *– Following differentiation, iAstrocytes were highly enriched for astrocyte quality control markers CD44 (yellow) and Vimentin (red).*

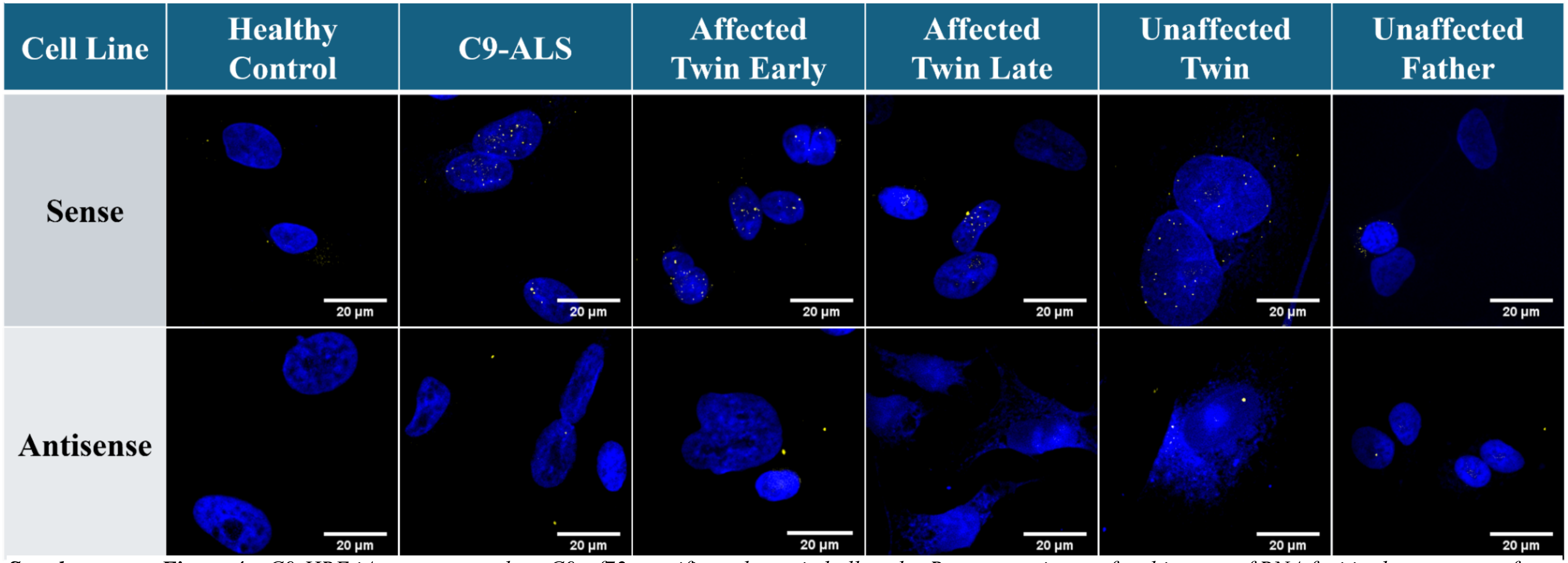


***Supplementary Figure 4*** *– C9-HRE iAstrocytes produce C9orf72-specific pathogenic hallmarks. Representative confocal images of RNA foci in the astrocytes from the pedigree. Images captured on Nikon A1 confocal microscope at 60x magnification and 1.5x zoom.*

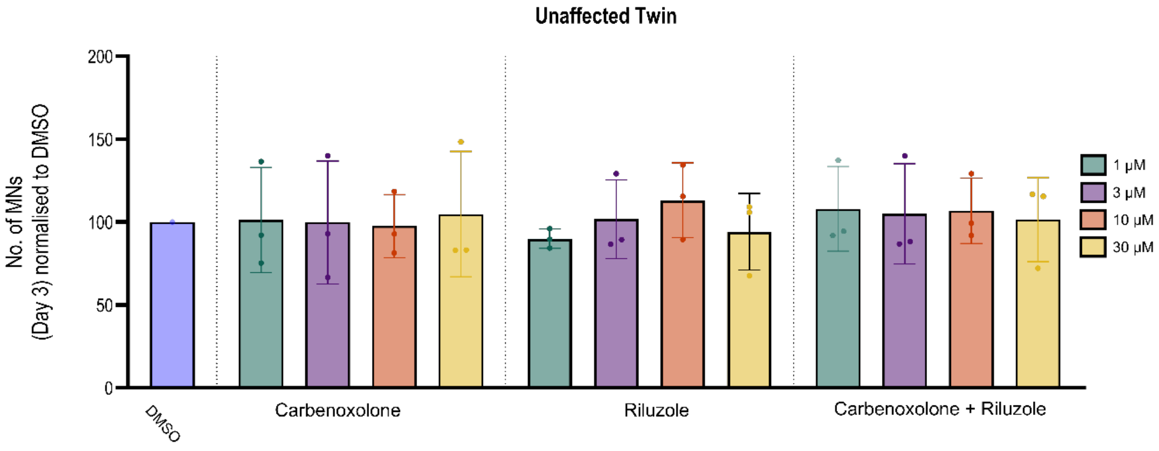


***Supplementary Figure 5*** *– Dose response curve of surviving HB9-GFP*$^{+}$ *MN following 72h co-culture with unaffected twin astrocytes treated with either DMSO, Carbenoxolone (1-30 µM), Riluzole (1-30 µM), or Carbenoxolone + Riluzole (1-30 µM of each). All doses of each condition were comparable to the DMSO baseline. Three biological replicates per condition, values normalised to the DMSO baseline of each biological repeat.*

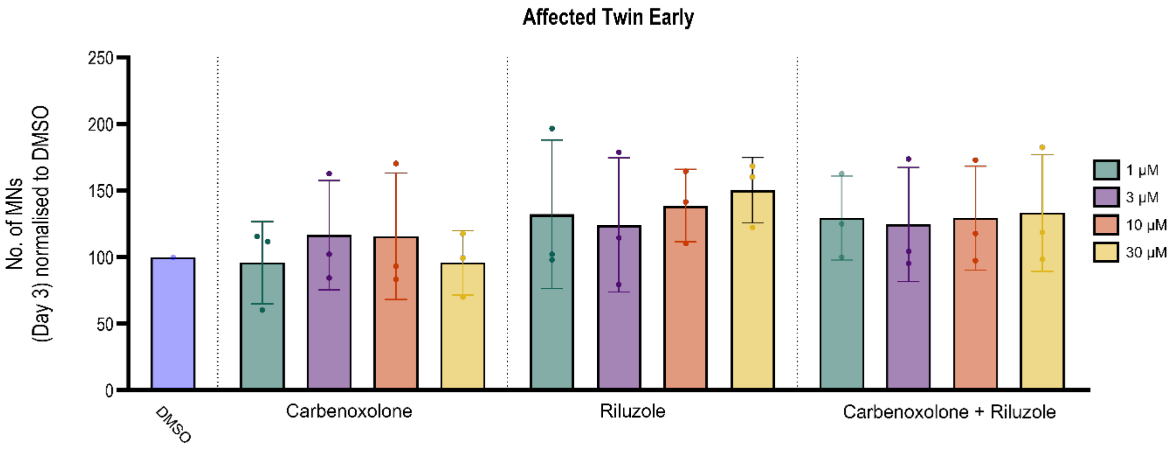


***Supplementary Figure 6*** *– Dose response curve of surviving HB9-GFP*$^{+}$ *MN following 72h co-culture with affected twin early astrocytes treated with either DMSO, Carbenoxolone (1-30 µM), Riluzole (1-30 µM), or Carbenoxolone + Riluzole (1-30 µM of each). A slight but variable rescue is observed following treatment with either 3 or 10 µM Carbenoxolone, with a consistent improvement in survival following treatment with Riluzole or Carbenoxolone and Riluzole in combination. Three biological replicates per condition, values normalised to the DMSO baseline of each biological repeat.*

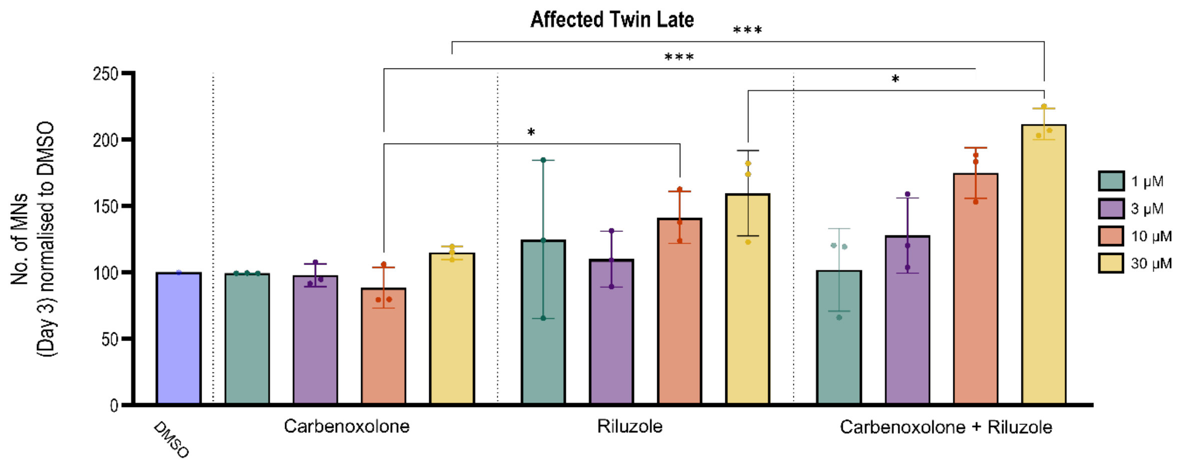


***Supplementary Figure 7*** *– Dose response curve of surviving HB9-GFP*$^{+}$ *MN following 72h co-culture with affected twin early astrocytes treated with either DMSO, Carbenoxolone (1-30 µM), Riluzole (1-30 µM), or Carbenoxolone + Riluzole (1-30 µM of each). Minimal rescue was observed following treatment with Carbenoxolone, but motor neuron surivival was improved with Riluzole treatment with the greatest effect at 10 or 30 µM. Co-dosing of Carbenoxolone and Riluzole at either 10 or 30 µM further increased this rescue. Significance test with one-way ANOVA with Sidak's multiple comparisons test (10 µM CBX vs. RIL vs. CBX + RIL; 30 µM CBX vs. RIL vs. CBX + RIL). ***, P <0.001. *, P <0.05. Three biological replicates per condition, values normalised to the DMSO baseline of each biological repeat.*

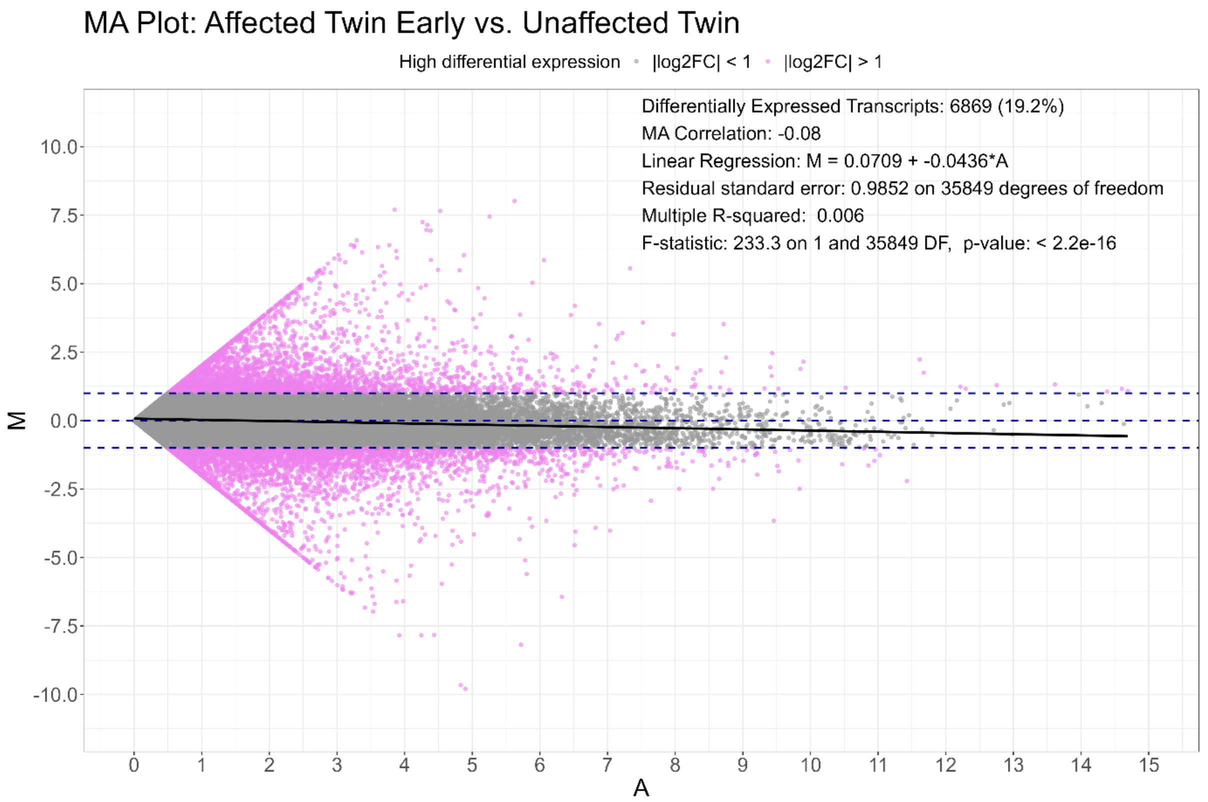


***Supplementary Figure 8*** *– MA Plot comparing TPM normalised transcript expression between affected twin early and unaffected twin. M represents Log2FC with a positive value equating to increased expression in affected twin early and reduced expression an increase in unaffected twin. A represents the TPM expression counts for each transcript averaged between the two samples. Differentially expressed transcripts with an absolute Log2FC >1 are noted in pink. The near zero value of the MA correlation and linear regression confirms that the dataset is not significantly biased towards either sample and therefore is suitable for downstream analysis.*

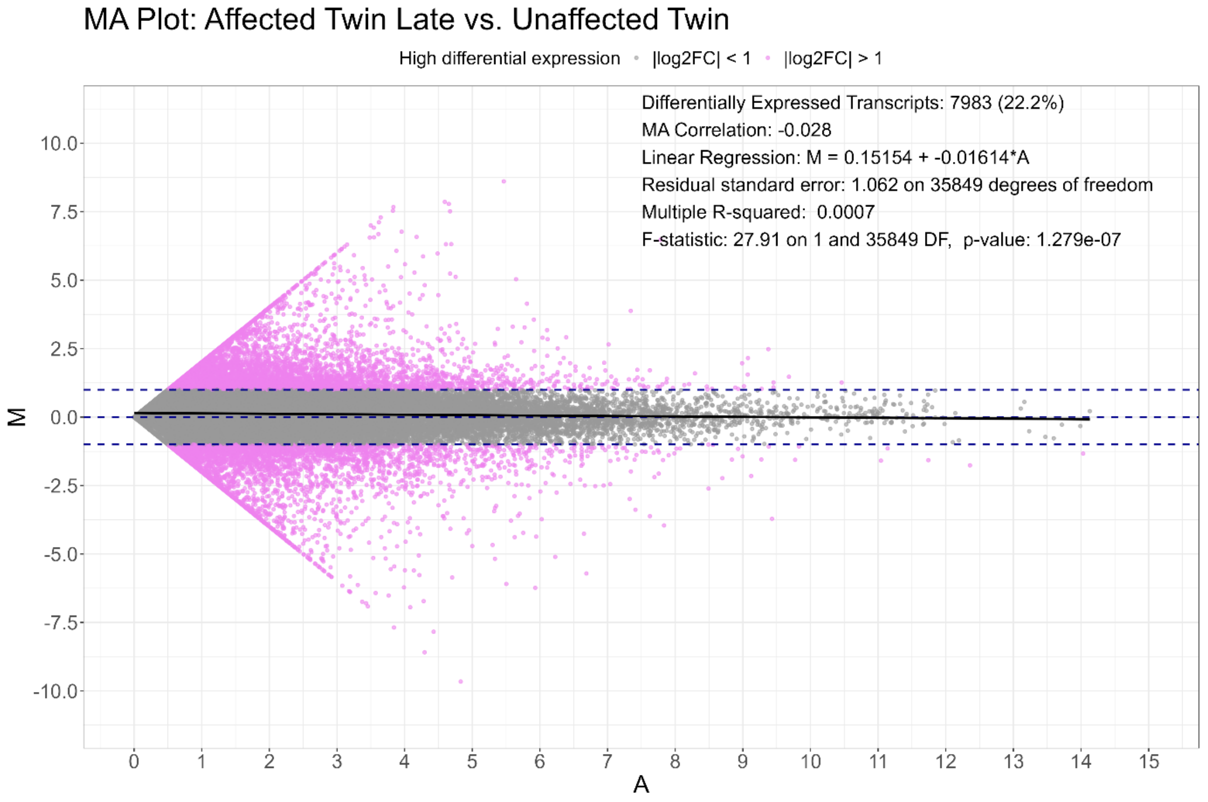


***Supplementary Figure 9*** *– MA Plot comparing TPM normalised transcript expression between affected twin late and unaffected twin. M represents Log2FC with a positive value equating to increased expression in affected twin late and reduced expression an increase in unaffected twin. A represents the TPM expression counts for each transcript averaged between the two samples. Differentially expressed transcripts with an absolute Log2FC >1 are noted in pink. The near zero value of the MA correlation and linear regression confirms that the dataset is not significantly biased towards either sample and therefore is suitable for downstream analysis.*

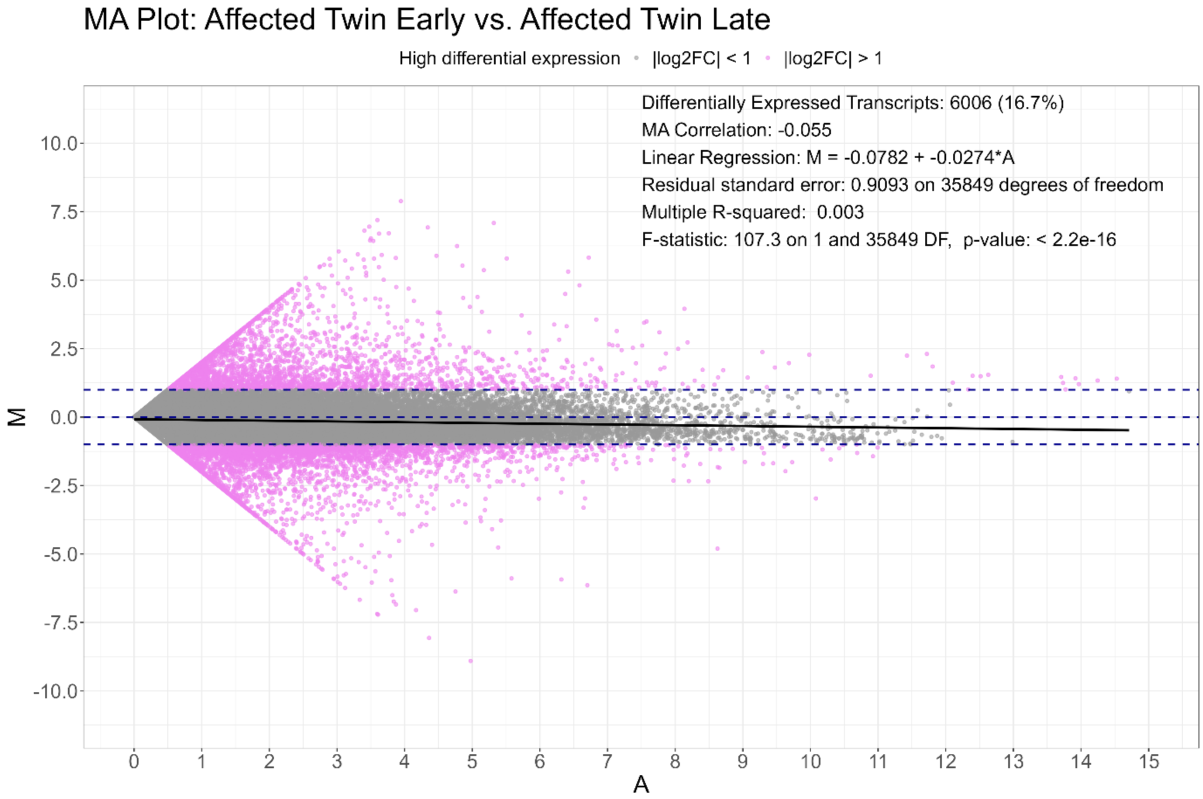


***Supplementary Figure 10*** *– MA Plot comparing TPM normalised transcript expression between affected twin early and affected twin late. M represents Log2FC with a positive value equating to increased expression in affected twin early and reduced expression an increase in affected twin late. A represents the TPM expression counts for each transcript averaged between the two samples. Differentially expressed transcripts with an absolute Log2FC >1 are noted in pink. The near zero value of the MA correlation and linear regression confirms that the dataset is not significantly biased towards either sample and therefore is suitable for downstream analysis.*

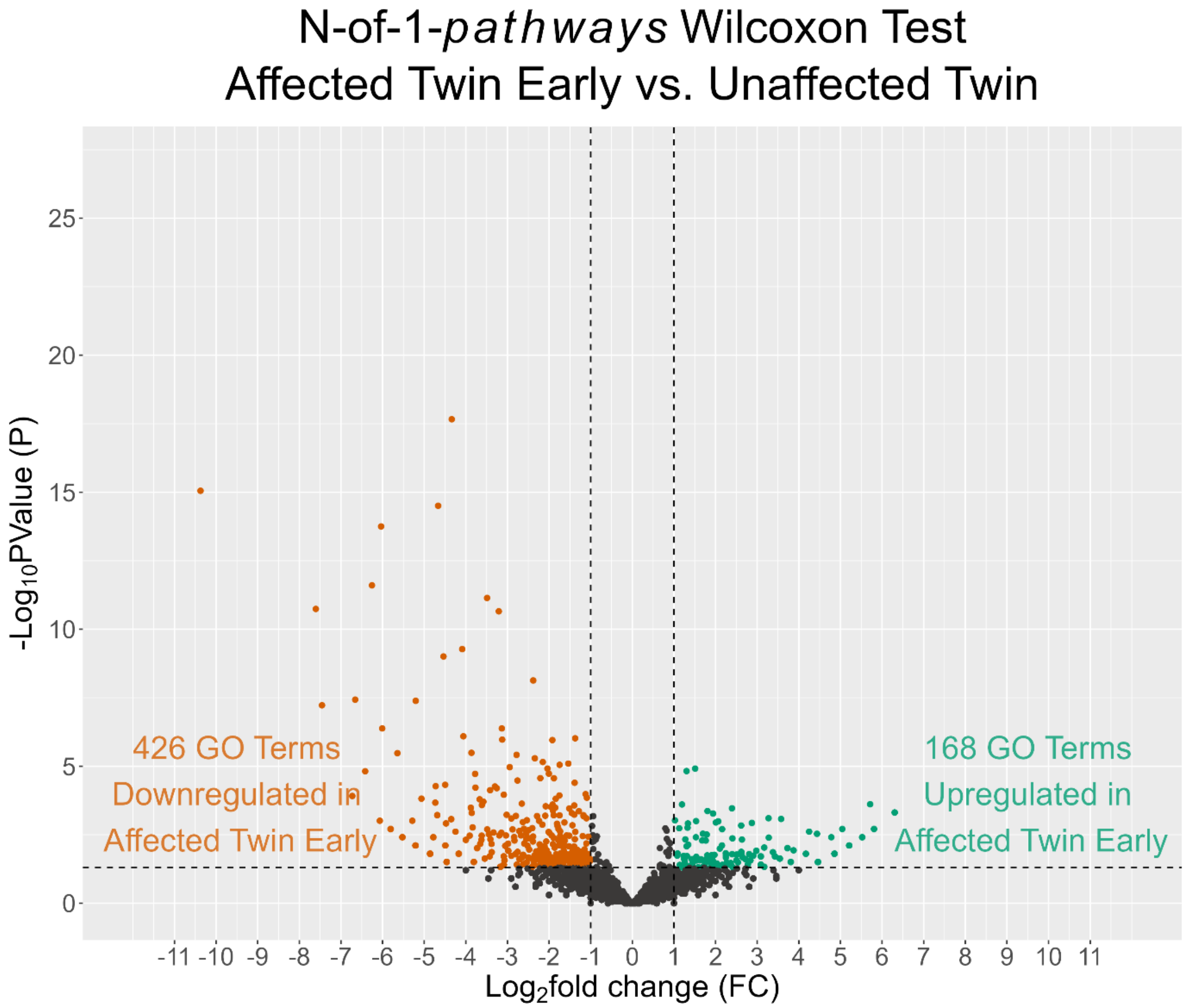


***Supplementary Figure 11*** *– Significantly dysregulated GO clusters identified with N-of-1-pathways Wilcoxon Test between the affected twin early and unaffected twin biopsies (contrast) following GRASPS Translatome RNA sequencing. In the affected twin early astrocytes 168 GO terms were significantly upregulated and 426 downregulated. The 2.5-fold increase in downregulated GO terms in the affected twin early suggests an impairment in canonical translation and global astrocytic function from asymptomatic to early symptomatic disease.*

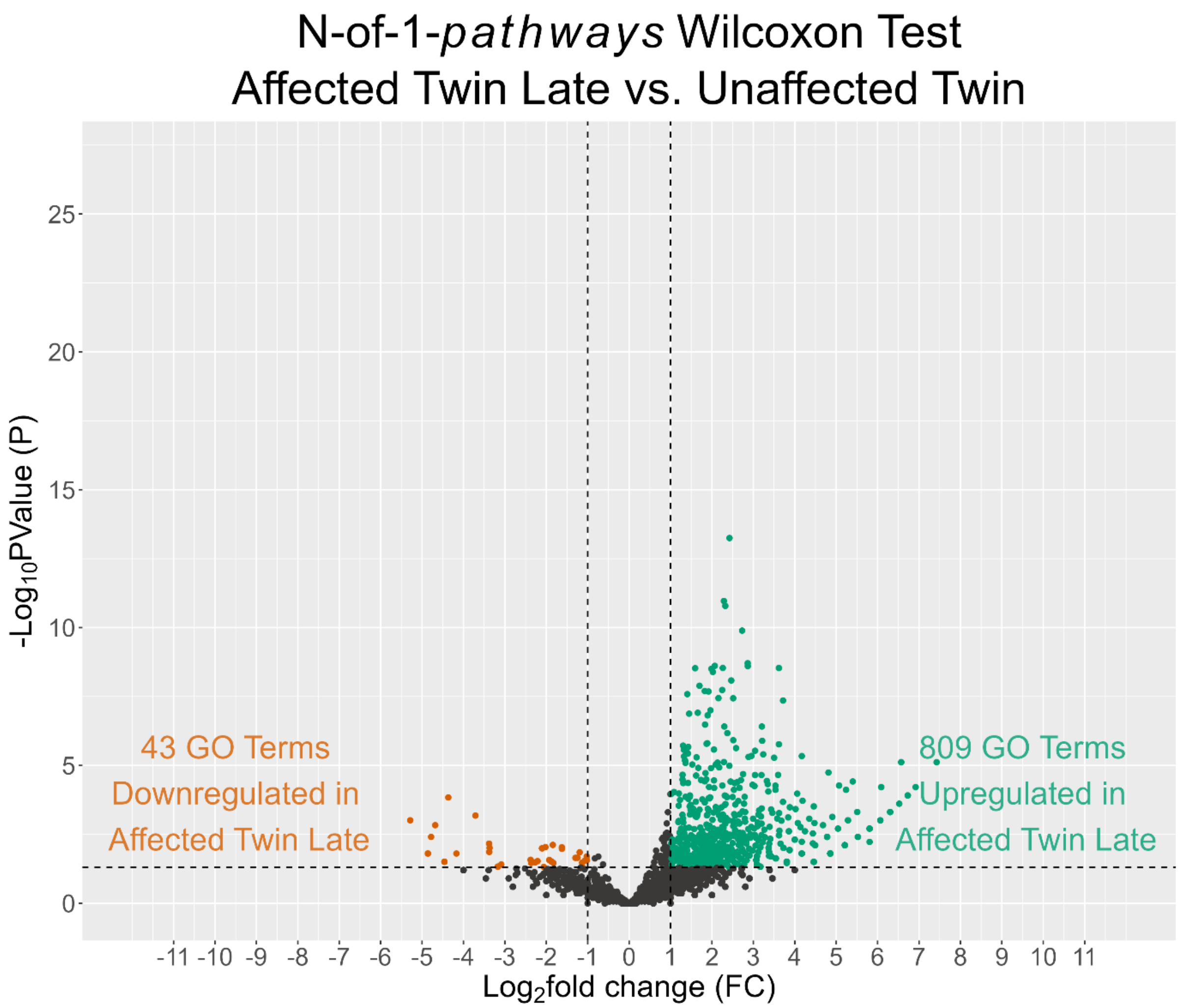


***Supplementary Figure 12*** *– Significantly dysregulated GO clusters identified with N-of-1-pathways Wilcoxon Test between the affected twin late and unaffected twin biopsies (contrast) following GRASPS Translatome RNA sequencing. In affected twin late astrocytes 809 GO terms were significantly upregulated, and 43 terms were downregulated. The 20-fold enrichment in upregulated GO terms in the late biopsy suggests that by end stage, the translational repression is reversed and increased over the asymptomatic baseline.*

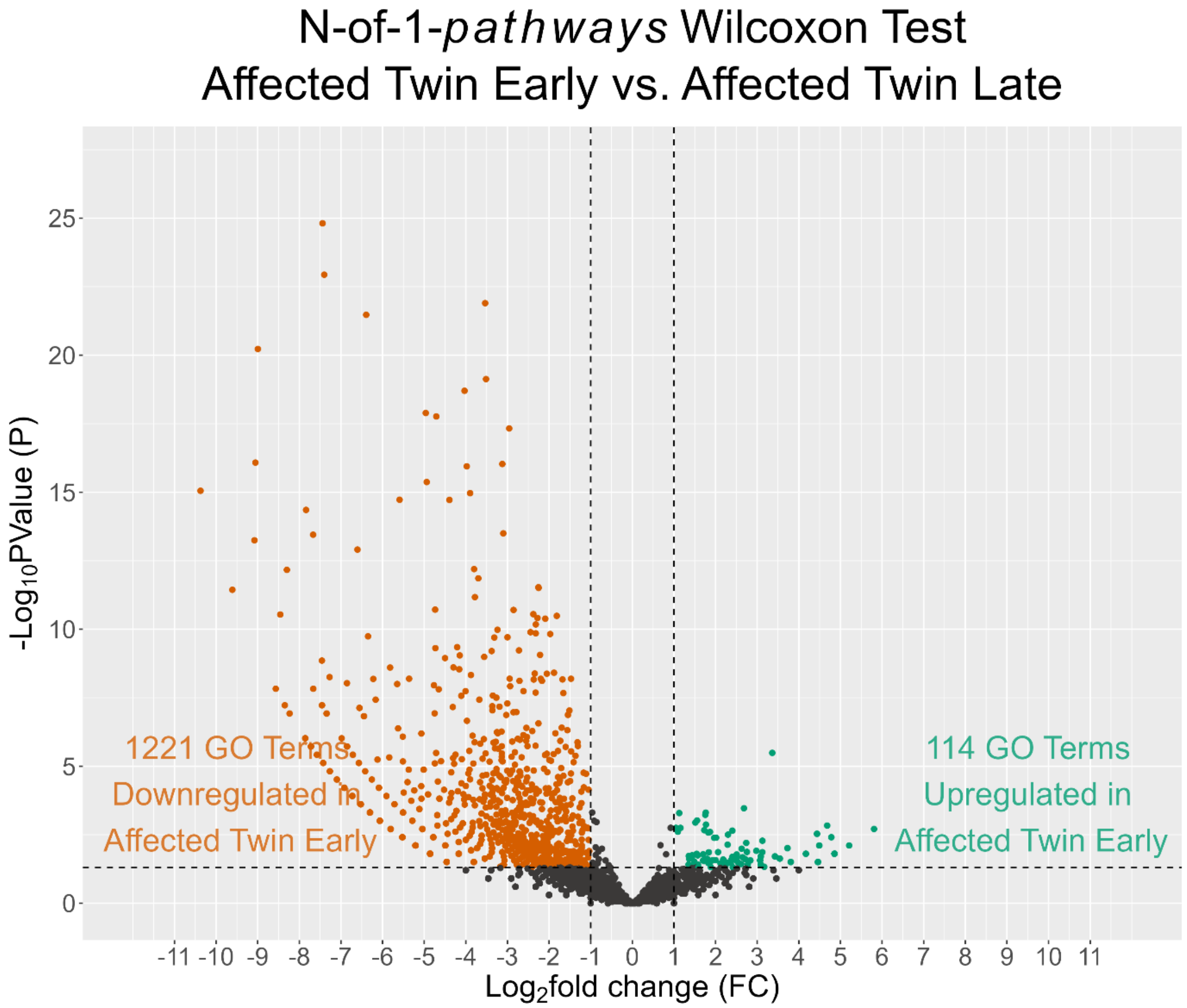


***Supplementary Figure 13** – Significantly dysregulated GO clusters identified with N-of-1-pathways Wilcoxon Test between the affected twin early and affected twin late biopsies (contrast) following GRASPS Translatome RNA sequencing. In affected twin early astrocytes 114 GO terms were significantly upregulated, and 1221 terms were downregulated. This 10-fold enrichment of downregulated terms further supports the suggestion that the early symptomatic translational repression is reversed by end stage.*

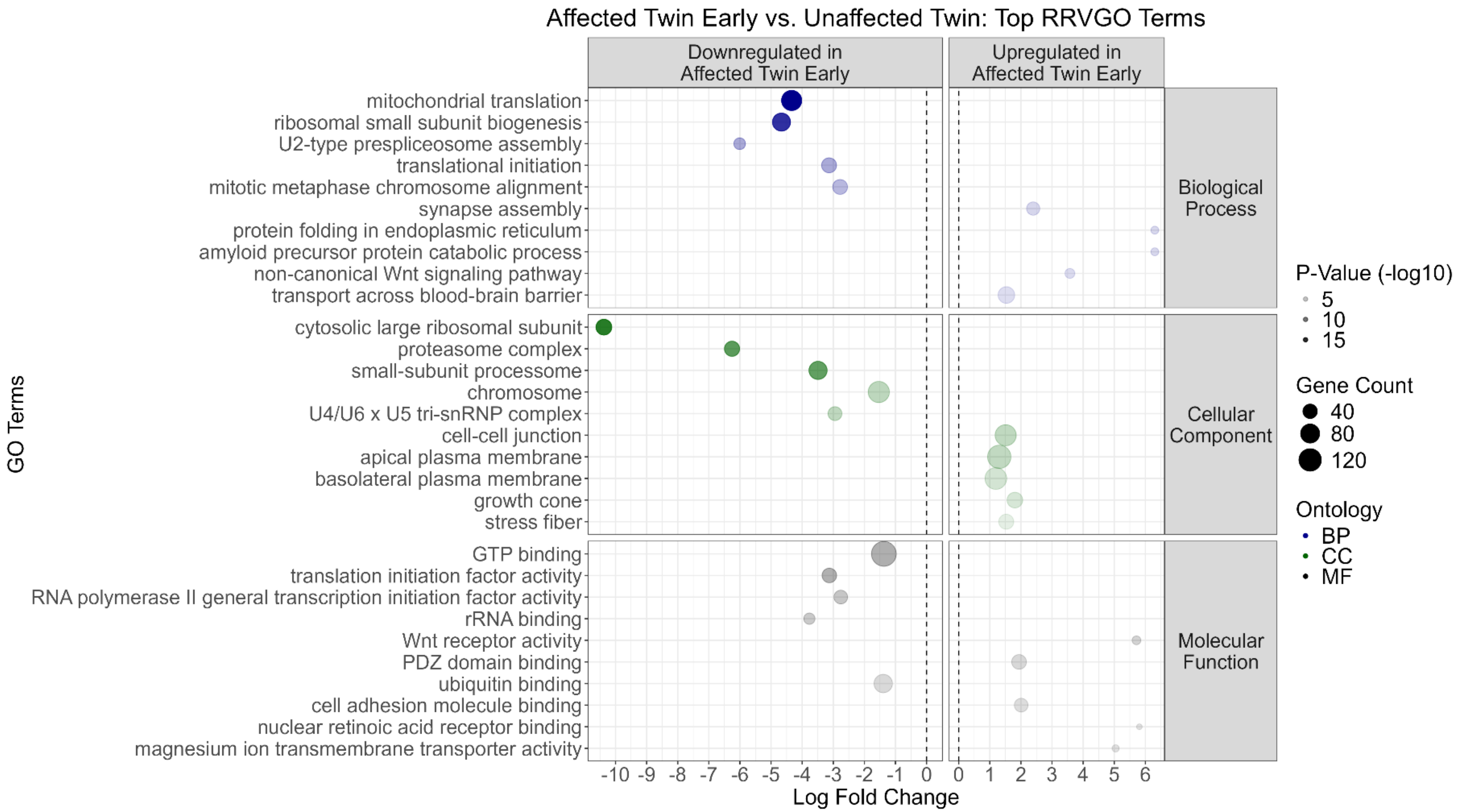


***Supplementary Figure 14*** *– Top GO terms identified between affected twin early and unaffected twin arranged by increasing significance (-log10 P-Value). The top 5 GO terms from each ontology (Biological Process, Cellular Component, and Molecular Function) with positive (upregulated in affected twin early) or negative (downregulated in affected twin early) fold change are displayed (for a total of 30 GO terms). All five of the most significantly dysregulated terms for Biological Process and Cellular Component, and 4/5 from Molecular Function are downregulated in affected twin early, further confirming the shift in astrocytic function identified in Supplementary Fig. 6.*

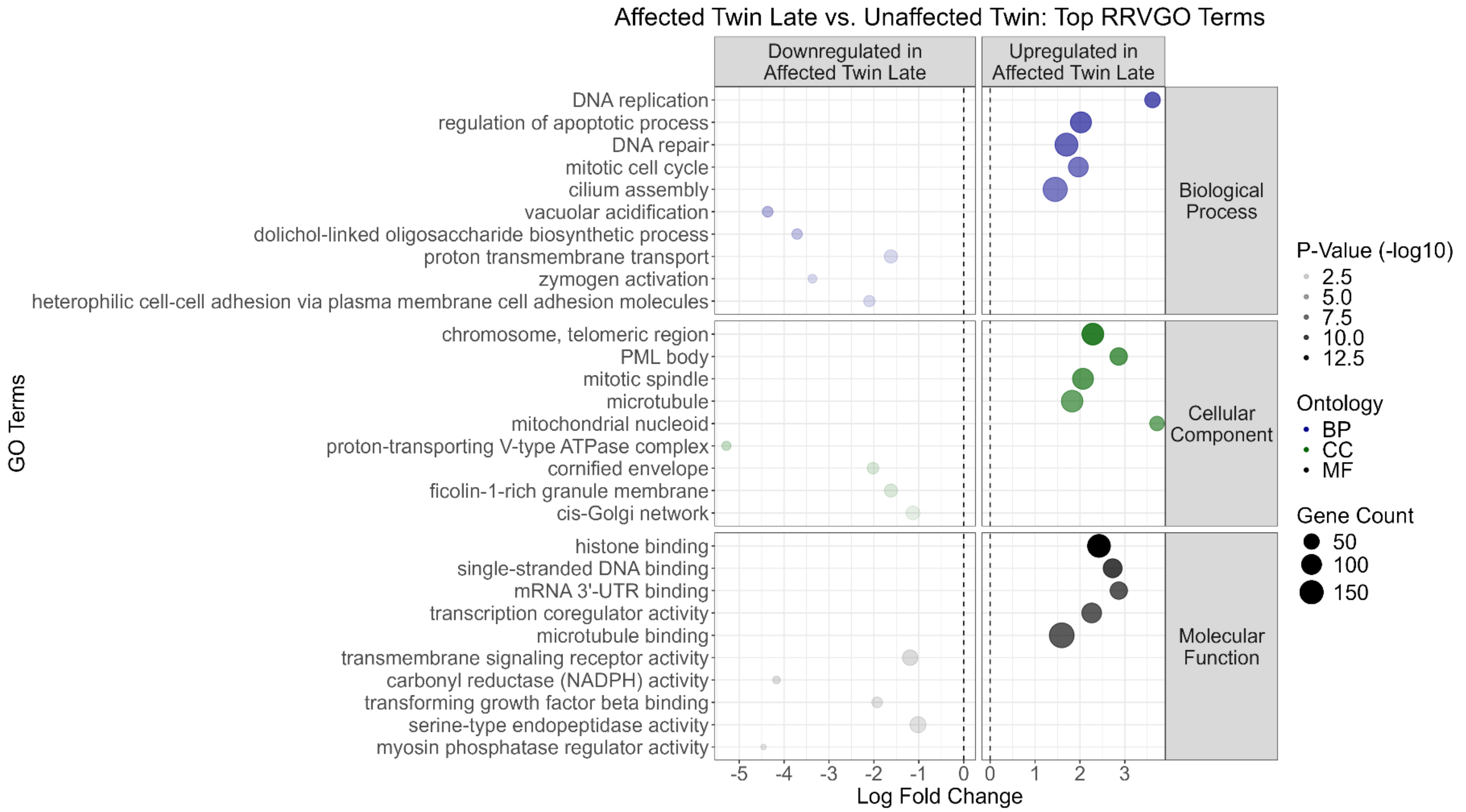


***Supplementary Figure 15*** *– Top GO terms identified between affected twin late and unaffected twin arranged by increasing significance (-log10 P-Value). The top 5 GO terms from each ontology (Biological Process, Cellular Component, and Molecular Function) with positive (upregulated in affected twin late) or negative (downregulated in affected twin late) fold change are displayed (for a total of 29 GO terms - only 4 terms upregulated in Unaffected Twin were associated with Cellular Component). The five most significantly dysregulated GO terms for each ontology are upregulated in affected twin late.*

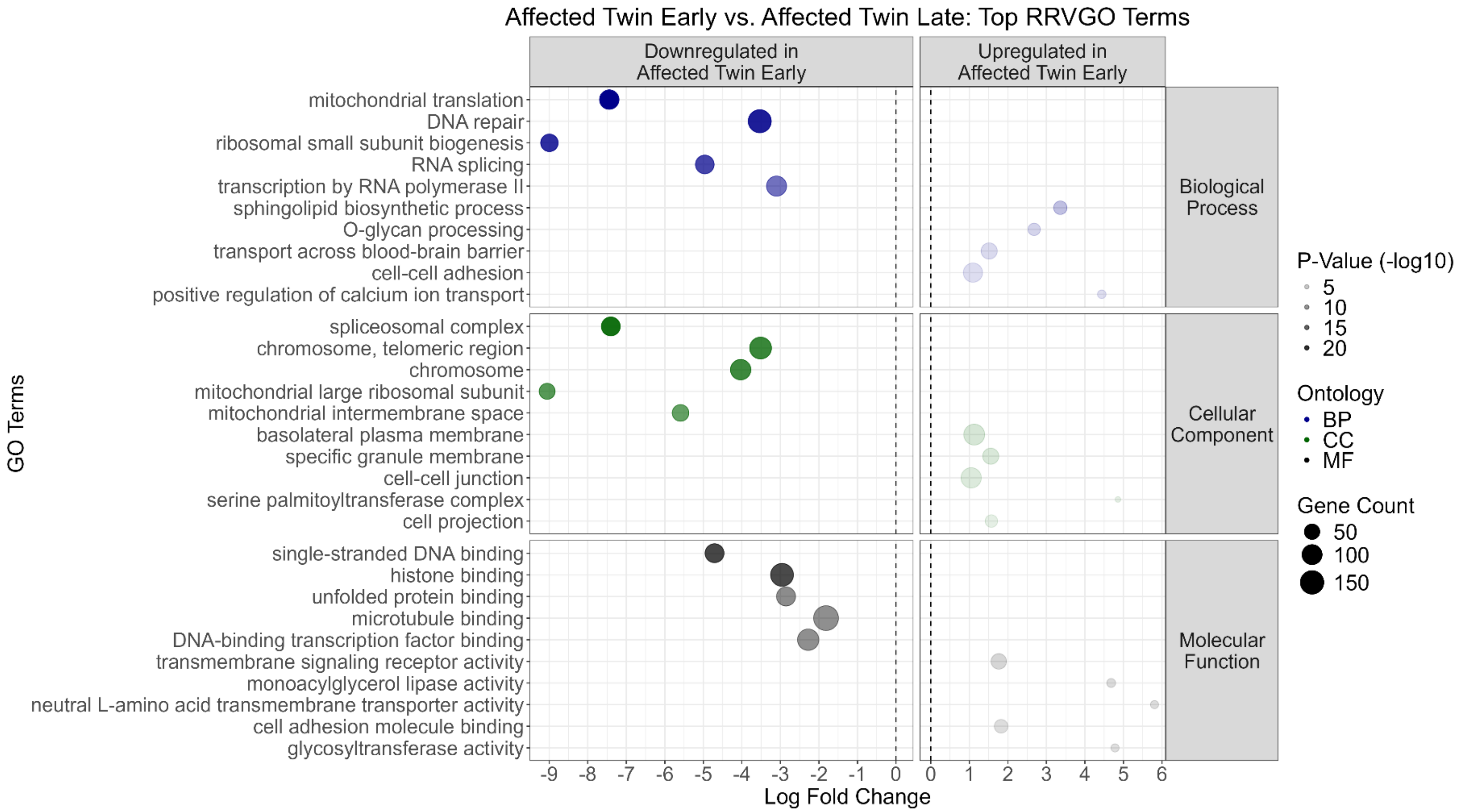


***Supplementary Figure 16*** *– Top GO terms identified between affected twin early and affected twin late arranged by increasing significance (-log10 P-Value). The top 5 GO terms from each ontology (Biological Process, Cellular Component, and Molecular Function) with positive (upregulated in affected twin early) or negative (downregulated in affected twin late) fold change are displayed (for a total of 30 GO terms). The five most significantly dysregulated GO terms for each ontology are downregulated in affected twin early.*

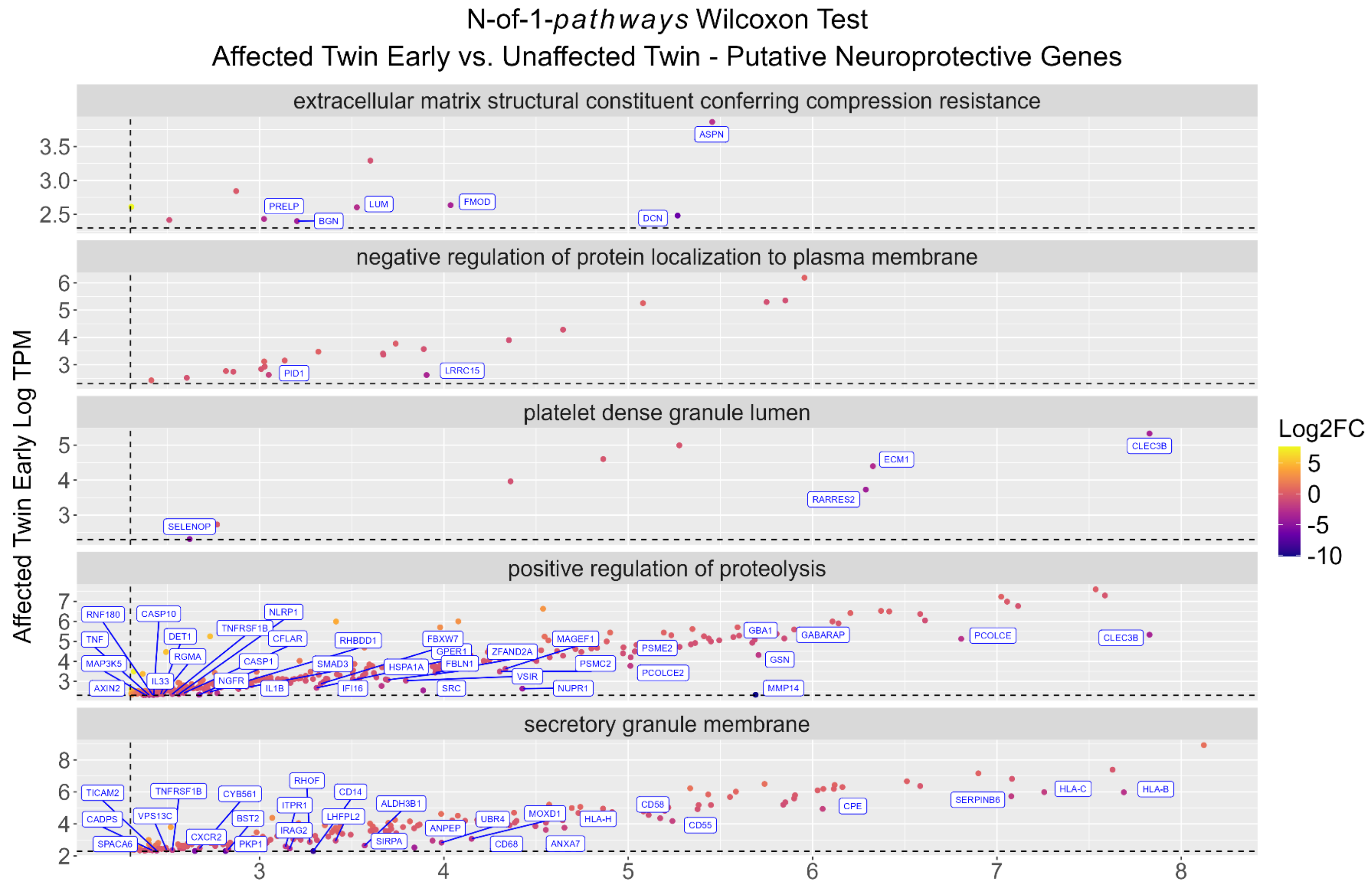


***Supplementary Figure 17*** *– Genes identified from GO terms dysregulated between asymptomatic and early symptomatic astrocytes. The 60 downregulated GO terms in Fig. 6C were hypothesised as a source of putative neuroprotective targets. Following RRVGO condensing, 36 parent terms remained and the contributing transcripts were extracted from the top 5 most significant terms. 75 transcripts with negative log2FC were extracted and labelled on scatter plots of the contributing GO terms (X-axis: Log TPM of affected twin early; Y-axis: Log TPM of unaffected twin, gradient of points: Log2FC).*

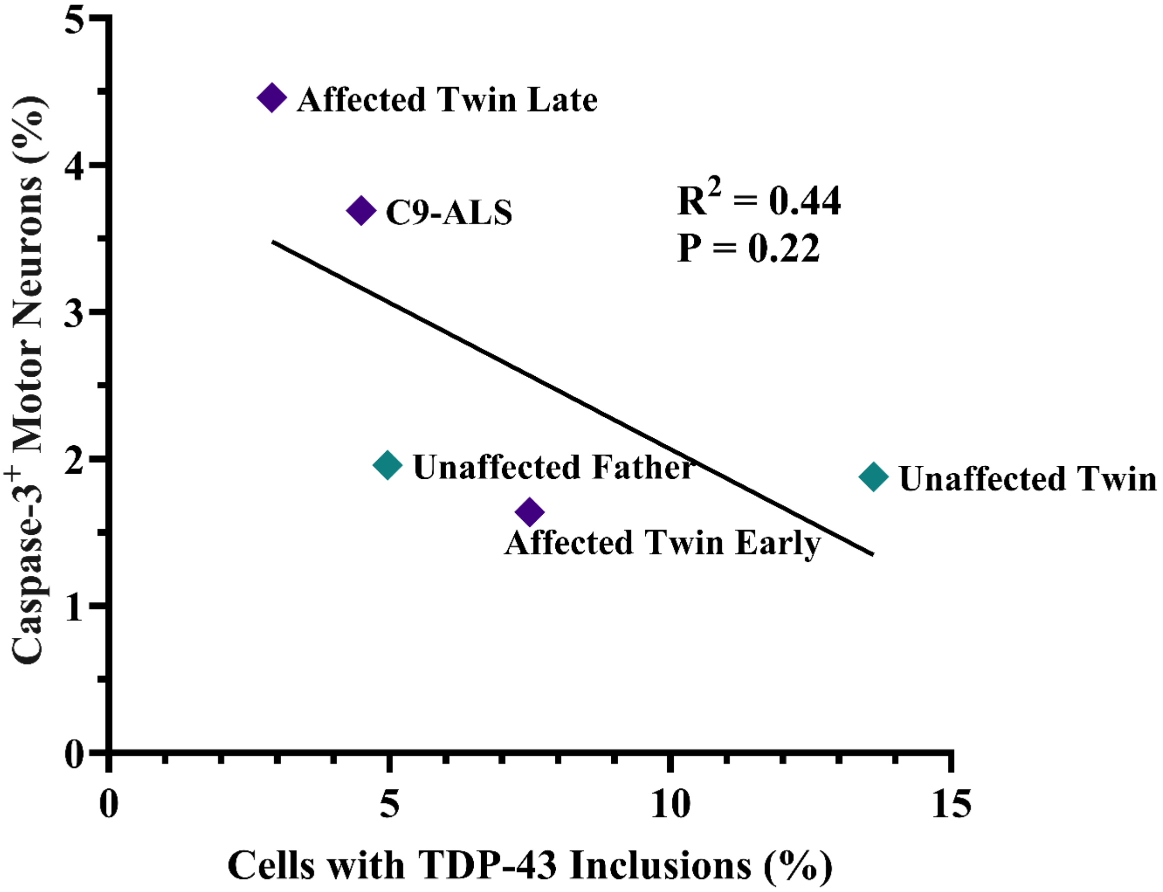


***Supplementary Figure 18*** *– Correlation plot showing the percentage of astrocytes containing cytoplasmic TDP-43 aggregates (X-axis) against astrocyte-induced motor neuron toxicity (Y-axis). Within cells carrying C9-HRE, a non-significant negative correlation is observed, whereby motor neuron toxicity decreases with increasing TDP-43 aggregation. Points marked by ♦*

#### Supplementary References


1. Sambrook J, Fritsch EF, Maniatis T. Molecular cloning: a laboratory manual (2nd ed.). 1989.

2. McKenna A, Hanna M, Banks E, et al. The Genome Analysis Toolkit: a MapReduce framework for analyzing next-generation DNA sequencing data. *Genome Res*. 2010;20(9):1297-303.

3. Karczewski KJ, Francioli LC, Tiao G, et al. The mutational constraint spectrum quantified from variation in 141,456 humans. *Nature*. 2020:581:434-443.

4. McLaren W, Gil L, Hunt SE, et al. The Ensembl Variant Effect Predictor. *Genome Biol*. 2016;17(1):122.

5. Pattamatta A, Nguyen L, Olafson HR, et al. Repeat length increases disease penetrance and severity in C9orf72 ALS/FTD BAC transgenic mice. *Hum Mol Genet*. 2021;29(24):3900-3918.

6. Beck J, Poulter M, Hensman D, et al. Large C9orf72 hexanucleotide repeat expansions are seen in multiple neurodegenerative syndromes and are more frequent than expected in the UK population. *Am J Hum Genet*. 2013;92(3):345-53.

7. Li H. Minimap2: pairwise alignment for nucleotide sequences, *Bioinformatics*. 2018;34(18):3094-3100.

8. Giesselmann P, Brändl B, Raimondeau E, et al. Analysis of short tandem repeat expansions and their methylation state with nanopore sequencing. *Nat Biotechnol*. 2019;37(12):1478-1481.

9. Shepheard SR, Parker MD, Cooper-Knock J, et al. Value of systematic genetic screening of patients with amyotrophic lateral sclerosis. *J Neurol Neurosurg Psychiatry*. 2021;92(5):510-518.

10. Iacoangeli A, Al Khleifat A, Sproviero W, et al. ALSgeneScanner: a pipeline for the analysis and interpretation of DNA sequencing data of ALS patients. *Amyotroph Lateral Scler Frontotemporal Degener*. 2019;20(3-4):207-215.

11. Chatterji S, Pachter L. Reference based annotation with GeneMapper. *Genome Biol*. 2006;7(4):R29.

12. Cleary EM, Pal S, Azam T, et al. Improved PCR based methods for detecting C9orf72 hexanucleotide repeat expansions. *Mol Cell Probes*. 2016;30(4):218-224.

13. Sproviero W, Shatunov A, Stahl D, et al. ATXN2 trinucleotide repeat length correlates with risk of ALS. *Neurobiol Aging*. 2017;51(178):e1-e9.

14. Magnani D, Chandran S, Wyllie DJA, Livesey MR. In Vitro Generation and Electrophysiological Characterization of OPCs and Oligodendrocytes from Human Pluripotent Stem Cells. *Methods Mol Biol*. 2019:1936:65-77.

15. Cooper-Knock J, Higginbottom A, Stopford MJ, et al. Antisense RNA foci in the motor neurons of C9ORF72-ALS patients are associated with TDP-43 proteinopathy. *Acta Neuropathol*. 2015;130(1):63-75.

16. Cooper-Knock J, Walsh MJ, Higginbottom A, et al. Sequestration of multiple RNA recognition motif-containing proteins by C9orf72 repeat expansions. *Brain*. 2014;137(7):2040-51.

17. Hautbergue G, Castelli L, Ferraiuolo L, et al. SRSF1-dependent nuclear export inhibition of *C9ORF72* repeat transcripts prevents neurodegeneration and associated motor deficits. *Nat Commun*. 2017;8:16063.

18. Quaegebeur A, Glaria I, Lashley T, et al. Soluble and insoluble dipeptide repeat protein measurements in *C9orf72*-frontotemporal dementia brains show regional differential solubility and correlation of poly-GR with clinical severity. *Acta Neuropathol Commun*. 2020;8:184.

19. Stopford MJ, Allen SP, Ferraiuolo L. A High-throughput and Pathophysiologically Relevant Astrocyte-motor Neuron Co-culture Assay for Amyotrophic Lateral Sclerosis Therapeutic Discovery. *Bio Protoc*. 2019;9(17):e3353.

20. Du ZW, Chen H, Liu H, et al. Generation and expansion of highly pure motor neuron progenitors from human pluripotent stem cells. *Nat Commun*. 2015;6:6626.

21. Besson H, Harwood C, Ekelund U, et al. Validation of the historical adulthood physical activity questionnaire (HAPAQ) against objective measurements of physical activity. *Int J Behav Nutr Phys Act*. 2010;7:54.

22. Lin Y, Dodd J, Cutillo L, et al. GRASPS: a simple-to-operate translatome technology reveals omics-hidden disease-associated pathways in TDP-43-related amyotrophic lateral sclerosis. *bioRxiv*. [Preprint] doi:10.1101/2024.03.04.583294.

23. Martin M. Cutadapt removes adapter sequences from high-throughput sequencing reads. *EMBnet.journal*. 2011;17(1):10-12.

24. Patro R, Duggal G, Love M, Irizarry R, Kingsford C. Salmon provides fast and bias-aware quantification of transcript expression. *Nat Methods*. 2017;14(4):417-419.

25. Pimentel H, Bray N, Puente S, Melsted P, Pachter L. Differential analysis of RNA-seq incorporating quantification uncertainty. *Nat Methods*. 2017;14(7):687-690.

26. Gardeux V, Achour I, Li J, et al. 'N-of-1-pathways' unveils personal deregulated mechanisms from a single pair of RNA-Seq samples: towards precision medicine. *J Am Med Inform Assoc*. 2014;21(6):1015-1025.

27. Schissler AG, Gardeux V, Li Q, et al. Dynamic changes of RNA-sequencing expression for precision medicine: N-of-1-pathways Mahalanobis distance within pathways of single subjects predicts breast cancer survival. *Bioinformatics*. 2015;31(12):i293-302.

28. Ashburner M, Ball CA, Blake JA, et al. Gene ontology: tool for the unification of biology. The Gene Ontology Consortium. *Nat Genet*. 2000;25(1):25-29.

29. Gene Ontology Consortium. The Gene Ontology knowledgebase in 2026. *Nucleic Acids Res*. 2026;54(1):1779-1792.
